## Supplemental Material for "Generalization of contextual fear is sex-specifically affected by high salt intake"

**S1 Table. Three-way repeated measures ANOVAs on context fear 'training' for control no shock mice across Experiments.**

S1A Table

| Experiment 1 | No Shock – Context Fear Training |
|---|---|
| Sex | F(1,30)=0.036 p=0.850 partial $\eta^2$=0.001 |
| Diet | F(1,30)=1.003 p=0.325 partial $\eta^2$=0.032 |
| Time | F(2.92,87.68)=3.688 **p=0.016** partial $\eta^2$=**0.109** |
| Time × Sex | F(2.92,87.68)=2.055 p=0.114 partial $\eta^2$=0.064 |
| Time × Diet | F(2.92,87.68)=0.653 p=0.579 partial $\eta^2$=0.021 |
| Sex × Diet | F(1,30)=0.000 p=0.990 partial $\eta^2$=0.000 |
| Time × Sex × Diet | F(2.92,87.68)=2.404 p=0.074 partial $\eta^2$=0.074 |

S1B Table

| Experiment 2 | No Shock – Context Fear Training |
|---|---|
| Sex | F(1,30)=0.516 p=0.478 partial $\eta^2$=0.017 |
| Diet | F(1,30)=0.012 p=0.915 partial $\eta^2$=0.000 |
| Time | F(3.05,91.62)=6.466 **p<0.001** partial $\eta^2$=**0.177** |
| Time × Sex | F(3.05,91.62)=1.258 p=0.294 partial $\eta^2$=0.040 |
| Time × Diet | F(3.05,91.62)=1.032 p=0.383 partial $\eta^2$=0.033 |
| Sex × Diet | F(1,30)=3.135 p=0.087 partial $\eta^2$=0.095 |
| Time × Sex × Diet | F(3.05,91.62)=0.710 p=0.551 partial $\eta^2$=0.023 |

S1C Table

| Experiment 3 | No Shock – Context Fear Training |
|---|---|
| Sex | F(1,27)=11.10 p=0.003 partial $\eta^2$=0.291 |
| Diet | F(1,27)=0.245 p=0.625 partial $\eta^2$=0.009 |
| Time | F(2.44,65.88)=3.016 p=0.046 partial $\eta^2$=0.100 |
| Time × Sex | F(2.44,65.88)=3.091 **p=0.042** partial $\eta^2$=**0.103** |
| Time × Diet | F(2.44,65.88)=0.668 p=0.545 partial $\eta^2$=0.024 |
| Sex × Diet | F(1,27)=2.753 p=0.109 partial $\eta^2$=0.093 |
| Time × Sex × Diet | F(2.44,65.88)=1.590 p=0.207 partial $\eta^2$=0.056 |

S1 Figure

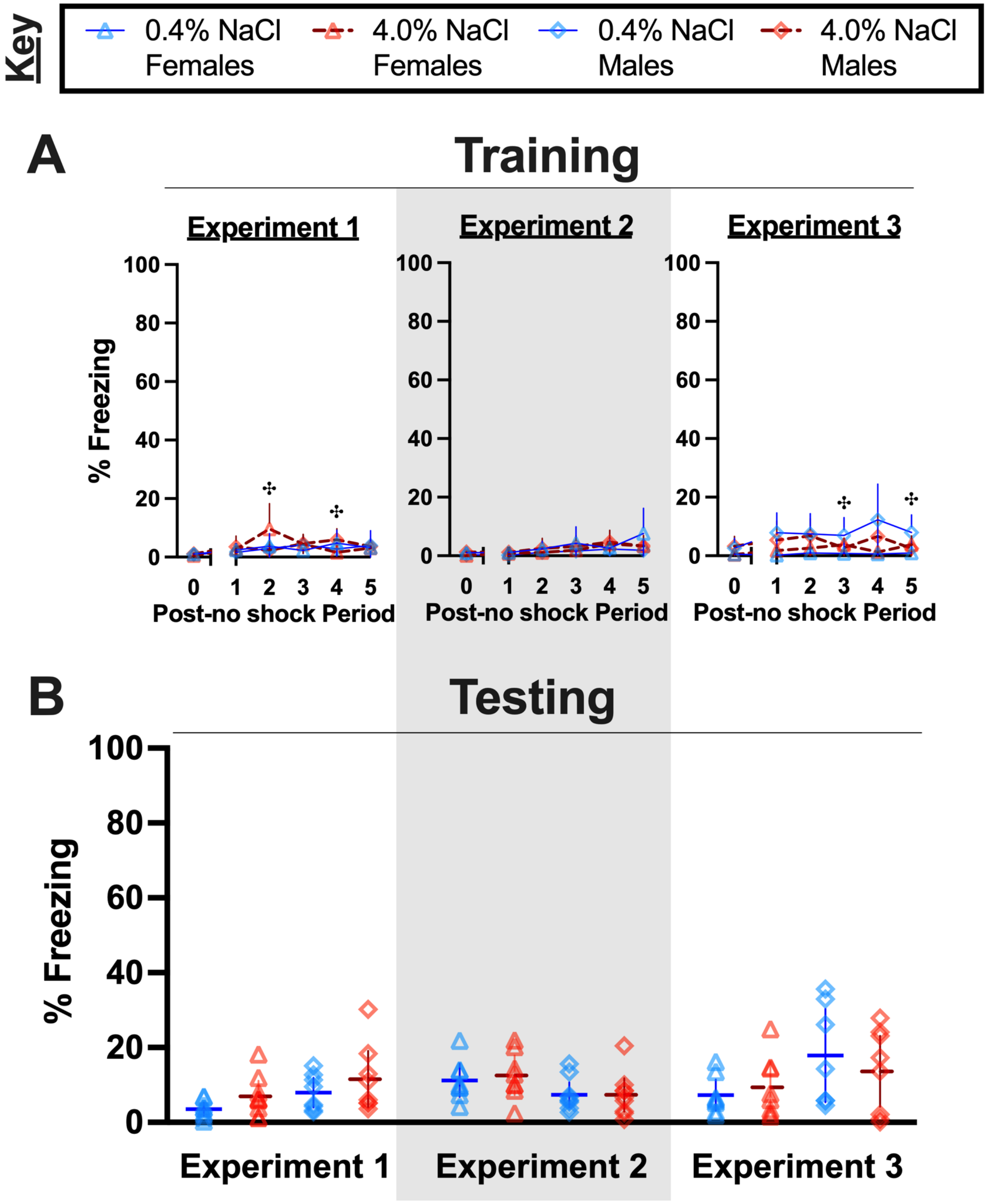

Key
0.4% NaCl Females
4.0% NaCl Females
0.4% NaCl Males
4.0% NaCl Males
A
Training
Experiment 1
Experiment 2
Experiment 3
% Freezing
100
80
60
40
20
0
0 1 2 3 4 5
Post-no shock Period
B
Testing
% Freezing
100
80
60
40
20
0
Experiment 1
Experiment 2
Experiment 3

**S1 Fig. Fear behavior in no shock control mice across Experiments.**

Females represented by triangles, males by diamonds; 0.4% NaCl represented by blue symbols and solid lines, 4.0% NaCl represented by red symbols and dashed lines. A) Percent freezing during ‘training’ of control mice exposed to the Training Context, but receiving no foot shocks for the duration of the 6 min ‘training’ session. B) Percent freezing during testing of control no shock mice exposed to the Training Context for minutes 2 through 6 of a 10 min testing session. Experiment 1: 0.4% NaCl females, n=9; 4.0% NaCl females, n=9; 0.4% NaCl males, n=8; 4.0% NaCl males, n=8. Experiment 2 (grey shading): 0.4% NaCl females, n=8; 4.0% NaCl females, n=8; 0.4% NaCl males, n=9; 4.0% NaCl males, n=9. Experiment 3: 0.4% NaCl females, n=8; 4.0% NaCl females, n=8; 0.4% NaCl males, n=7; 4.0% NaCl males, n=8. Data are graphed as mean ± 95% confidence interval. ‡indicates $p<0.05$ difference between males consuming 0.4% and 4.0% NaCl diets at indicated time points.

**S2 Table. Three-way ANOVAs on context fear testing of control no shock mice across Experiments.**

| Context Fear Expression | No Shock Groups Across Experiments | | |
|---|---|---|---|
| Sex | $F(1,87)=2.852$ | $p=0.095$ | partial $\eta^2=0.032$ |
| Diet | $F(1,87)=0.497$ | $p=0.483$ | partial $\eta^2=0.006$ |
| Experiment | $F(2,87)=3.187$ | $p=0.046$ | partial $\eta^2=0.068$ |
| Sex × Diet | $F(1,87)=0.748$ | $p=0.390$ | partial $\eta^2=0.009$ |
| Sex × Experiment | $F(2,87)=6.137$ | **$p=0.003$** | partial $\eta^2=$**0.124** |
| Diet × Experiment | $F(2,87)=0.839$ | $p=0.436$ | partial $\eta^2=0.019$ |
| Sex × Diet × Experiment | $F(2,87)=0.447$ | $p=0.641$ | partial $\eta^2=0.010$ |

S2 Figure

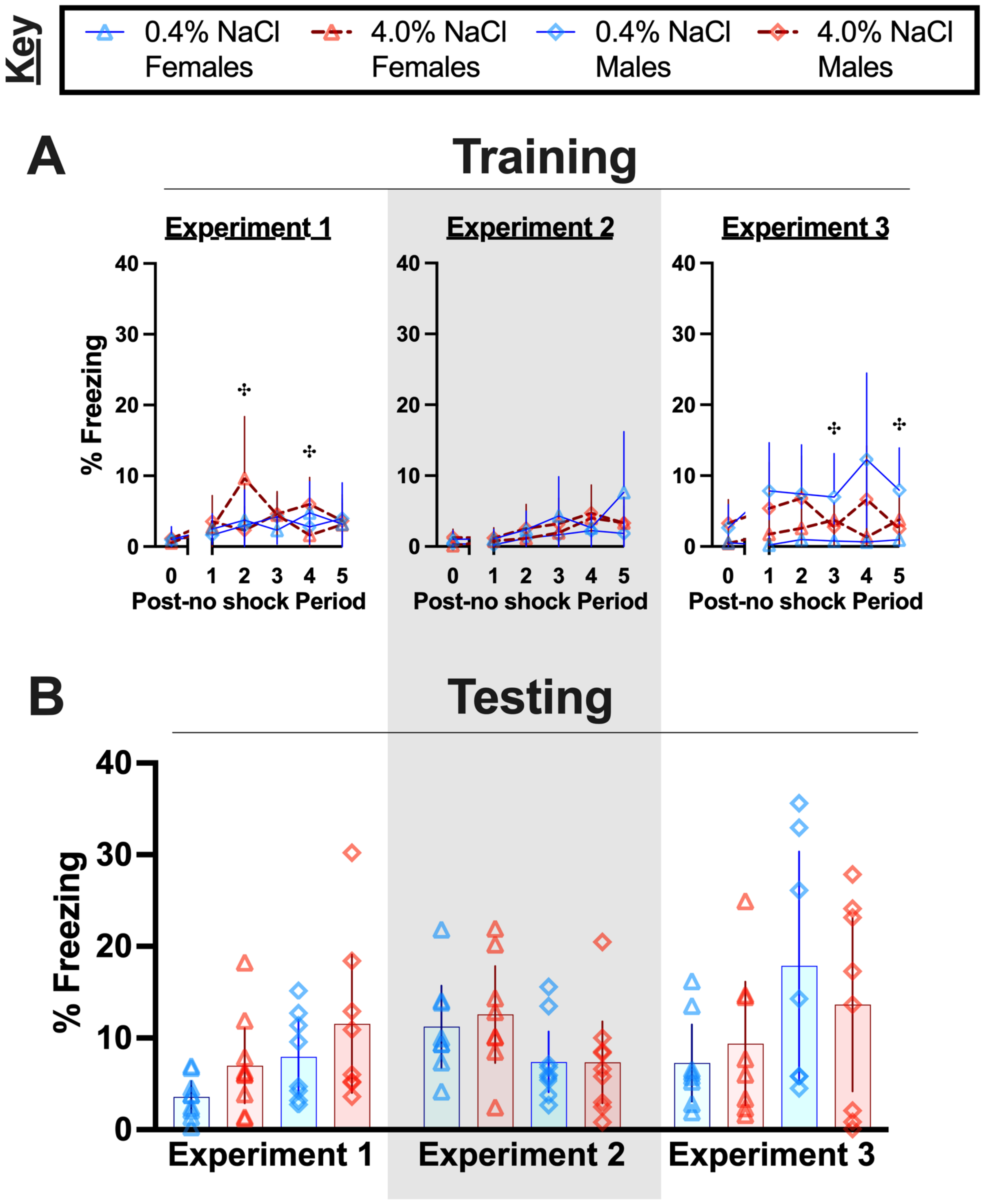
Key
0.4% NaCl Females
4.0% NaCl Females
0.4% NaCl Males
4.0% NaCl Males
A
Training
Experiment 1
Experiment 2
Experiment 3
% Freezing
Post-no shock Period
B
Testing
% Freezing
Experiment 1
Experiment 2
Experiment 3

**S2 Fig. Fear behavior in no shock control mice across Experiments with shortened Y axes**.

Females represented by triangles, males by diamonds; 0.4% NaCl represented by blue symbols and solid lines, 4.0% NaCl represented by red symbols and dashed lines. A) Percent freezing during 'training' of control mice exposed to the Training Context, but receiving no foot shocks for the duration of the 6 min 'training' session. B) Percent freezing during testing of control no shock mice exposed to the Training Context for minutes 2 through 6 of a 10 min testing session. Experiment 1: 0.4% NaCl females, n=9; 4.0% NaCl females, n=9; 0.4% NaCl males, n=8; 4.0% NaCl males, n=8. Experiment 2 (grey shading): 0.4% NaCl females, n=8; 4.0% NaCl females, n=8; 0.4% NaCl males, n=9; 4.0% NaCl males, n=9. Experiment 3: 0.4% NaCl females, n=8; 4.0% NaCl females, n=8; 0.4% NaCl males, n=7; 4.0% NaCl males, n=8. Data are graphed as mean ± 95% confidence interval. $^{‡}$indicates $p<0.05$ difference between males consuming 0.4% and 4.0% NaCl diets at indicated time points.

**S3 Table. Three-way repeated measures ANOVAs on context fear training for context fear conditioned mice of both sexes in Experiment 1.**

S3A Table

| **Females** | **Experiment 1 – Context Fear Training** |
|---|---|
| Diet | $F(1,29)=0.145$ $p=0.706$ partial $\eta^2=0.005$ |
| Context | $F(1,29)=0.009$ $p=0.927$ partial $\eta^2=0.000$ |
| Time | $F(4.00,115.9)=111.3$ $\mathbf{p<0.001}$ partial $\eta^2=\mathbf{0.793}$ |
| Time × Diet | $F(4.00,115.9)=0.377$ $p=0.824$ partial $\eta^2=0.013$ |
| Time × Context | $F(4.00,115.9)=0.844$ $p=0.500$ partial $\eta^2=0.028$ |
| Diet × Context | $F(1,29)=0.024$ $p=0.879$ partial $\eta^2=0.001$ |
| Time × Diet × Context | $F(4.00,115.9)=0.849$ $p=0.497$ partial $\eta^2=0.028$ |

S3B Table

| **Males** | **Experiment 1 – Context Fear Training** |
|---|---|
| Diet | $F(1,28)=0.538$ $p=0.470$ partial $\eta^2=0.019$ |
| Context | $F(1,28)=0.033$ $p=0.857$ partial $\eta^2=0.001$ |
| Time | $F(3.56,99.68)=59.79$ $\mathbf{p<0.001}$ partial $\eta^2=\mathbf{0.681}$ |
| Time × Diet | $F(3.56,99.68)=0.167$ $p=0.941$ partial $\eta^2=0.006$ |
| Time × Context | $F(3.56,99.68)=0.271$ $p=0.877$ partial $\eta^2=0.010$ |
| Diet × Context | $F(1,28)=0.013$ $p=0.909$ partial $\eta^2=0.000$ |
| Time × Diet × Context | $F(3.56,99.68)=0.266$ $p=0.880$ partial $\eta^2=0.009$ |

**S4 Table. Three-way repeated measures ANOVAs on context fear training for context fear conditioned mice of both sexes in Experiment 2.**

S4A Table

| **Females** | **Experiment 2 – Context Fear Training** |
|---|---|
| Diet | F(1,29)=0.083 p=0.775 partial $\eta^2$=0.003 |
| Context | F(1,29)=0.097 p=0.758 partial $\eta^2$=0.003 |
| Time | F(4.11,119.1)=121.2 **p<0.001** partial $\eta^2$=**0.807** |
| Time × Diet | F(4.11,119.1)=0.354 p=0.845 partial $\eta^2$=0.012 |
| Time × Context | F(4.11,119.1)=0.259 p=0.908 partial $\eta^2$=0.009 |
| Diet × Context | F(1,29)=0.104 p=0.750 partial $\eta^2$=0.004 |
| Time × Diet × Context | F(4.11,119.1)=0.537 p=0.714 partial $\eta^2$=0.018 |

S4 Table

| **Males** | **Experiment 2 – Context Fear Training** |
|---|---|
| Diet | F(1,31)=2.080 p=0.159 partial $\eta^2$=0.063 |
| Context | F(1,31)=0.111 p=0.741 partial $\eta^2$=0.004 |
| Time | F(3.26,101.0)=103.1 **p<0.001** partial $\eta^2$=**0.769** |
| Time × Diet | F(3.26,101.0)=2.166 p=0.091 partial $\eta^2$=0.065 |
| Time × Context | F(3.26,101.0)=0.402 p=0.768 partial $\eta^2$=0.013 |
| Diet × Context | F(1,31)=1.332 p=0.257 partial $\eta^2$=0.041 |
| Time × Diet × Context | F(3.26,101.0)=1.512 p=0.213 partial $\eta^2$=0.047 |

**S5 Table. Three-way repeated measures ANOVAs on context fear training for context fear conditioned mice of both sexes in Experiment 3.**

S5A Table

| **Females** | **Experiment 3 – Context Fear Training** |
|---|---|
| Diet | $F(1,30)=0.777$ $p=0.385$ partial $\eta^2=0.025$ |
| Context | $F(1,30)=0.514$ $p=0.479$ partial $\eta^2=0.017$ |
| Time | $F(3.82,114.7)=100.9$ $\mathbf{p<0.001}$ partial $\eta^2=\mathbf{0.771}$ |
| Time × Diet | $F(3.82,114.7)=2.287$ $p=0.067$ partial $\eta^2=0.071$ |
| Time × Context | $F(3.82,114.7)=1.008$ $p=0.404$ partial $\eta^2=0.033$ |
| Diet × Context | $F(1,30)=0.250$ $p=0.621$ partial $\eta^2=0.008$ |
| Time × Diet × Context | $F(3.82,114.7)=0.185$ $p=0.940$ partial $\eta^2=0.006$ |

S5B Table

| **Males** | **Experiment 3 – Context Fear Training** |
|---|---|
| Diet | $F(1,27)=1.070$ $p=0.310$ partial $\eta^2=0.038$ |
| Context | $F(1,27)=0.241$ $p=0.628$ partial $\eta^2=0.009$ |
| Time | $F(2.84,76.71)=79.15$ $\mathbf{p<0.001}$ partial $\eta^2=\mathbf{0.746}$ |
| Time × Diet | $F(2.84,76.71)=0.332$ $p=0.792$ partial $\eta^2=0.012$ |
| Time × Context | $F(2.84,76.71)=0.146$ $p=0.924$ partial $\eta^2=0.005$ |
| Diet × Context | $F(1,27)=0.533$ $p=0.472$ partial $\eta^2=0.019$ |
| Time × Diet × Context | $F(2.84,76.71)=0.413$ $p=0.733$ partial $\eta^2=0.015$ |

**S6 Table. Three-way repeated measures ANOVAs on full 10 min time course of context fear testing for control no shock mice across Experiments.**

| Experiment 1 | No Shock – Context Fear Testing | | |
|---|---|---|---|
| Sex | $F(1,30)=5.414$ | **$p=0.027$** | partial $\eta^2=$**0.153** |
| Diet | $F(1,30)=1.972$ | $p=0.170$ | partial $\eta^2=0.062$ |
| Time | $F(8.43,252.8)=1.005$ | $p=0.453$ | partial $\eta^2=0.032$ |
| Time × Sex | $F(8.43,252.8)=0.602$ | $p=0.784$ | partial $\eta^2=0.020$ |
| Time × Diet | $F(8.43,252.8)=0.706$ | $p=0.693$ | partial $\eta^2=0.023$ |
| Sex × Diet | $F(1,30)=0.138$ | $p=0.713$ | partial $\eta^2=0.005$ |
| Time × Sex × Diet | $F(8.43,252.8)=1.022$ | $p=0.421$ | partial $\eta^2=0.033$ |

| Experiment 2 | No Shock – Context Fear Testing | | |
|---|---|---|---|
| Sex | $F(1,30)=5.647$ | **$p=0.024$** | partial $\eta^2=$**0.158** |
| Diet | $F(1,30)=0.000$ | $p=0.992$ | partial $\eta^2=0.000$ |
| Time | $F(7.15,214.5)=3.614$ | **$p<0.001$** | partial $\eta^2=$**0.108** |
| Time × Sex | $F(7.15,214.5)=1.753$ | $p=0.097$ | partial $\eta^2=0.055$ |
| Time × Diet | $F(7.15,214.5)=1.166$ | $p=0.323$ | partial $\eta^2=0.037$ |
| Sex × Diet | $F(1,30)=0.317$ | $p=0.578$ | partial $\eta^2=0.010$ |
| Time × Sex × Diet | $F(7.15,214.5)=0.863$ | $p=0.538$ | partial $\eta^2=0.028$ |

| Experiment 3 | No Shock – Context Fear Testing | | |
|---|---|---|---|
| Sex | $F(1,27)=3.799$ | $p=0.062$ | partial $\eta^2=0.123$ |
| Diet | $F(1,27)=0.002$ | $p=0.966$ | partial $\eta^2=0.000$ |
| Time | $F(7.48,202.0)=1.440$ | $p=0.186$ | partial $\eta^2=0.051$ |
| Time × Sex | $F(7.48,202.0)=1.459$ | $p=0.179$ | partial $\eta^2=0.051$ |
| Time × Diet | $F(7.48,202.0)=0.549$ | $p=0.807$ | partial $\eta^2=0.020$ |
| Sex × Diet | $F(1,27)=0.890$ | $p=0.354$ | partial $\eta^2=0.032$ |
| Time × Sex × Diet | $F(7.48,202.0)=0.459$ | $p=0.874$ | partial $\eta^2=0.017$ |

S3 Figure

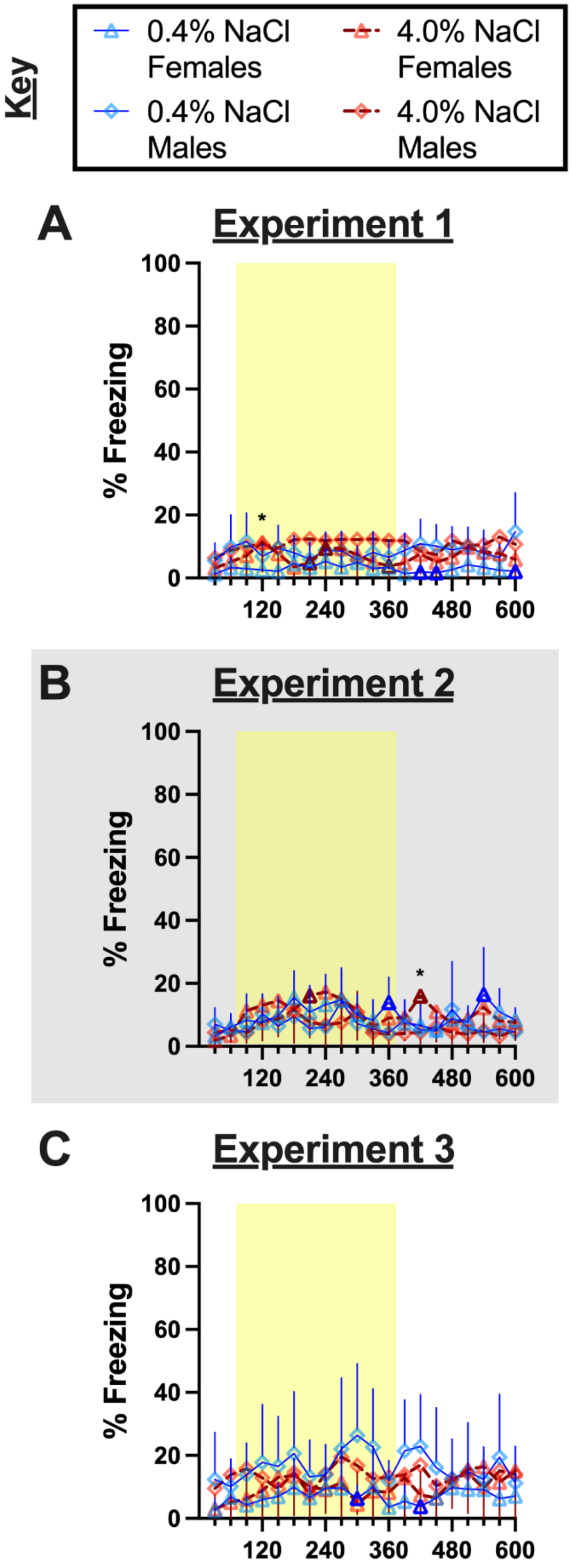
Key
0.4% NaCl Females
4.0% NaCl Females
0.4% NaCl Males
4.0% NaCl Males
A
Experiment 1
% Freezing
100
80
60
40
20
0
120
240
360
480
600
*
B
Experiment 2
% Freezing
100
80
60
40
20
0
120
240
360
480
600
*
C
Experiment 3
% Freezing
100
80
60
40
20
0
120
240
360
480
600

**S3 Fig. Time course of context fear expression testing in no shock groups**. Females represented by triangles, males by diamonds; 0.4% NaCl represented by blue symbols and solid lines, 4.0% NaCl represented by red symbols and dashed lines. Percent freezing for each 30 s bin of the 10 min testing session is graphed for all mice. Yellow shading indicates minutes 2 through 6, which were averaged and analyzed as our original measure of context fear expression. Testing occurred A) 48 h after training in Experiment 1, during which mice underwent two weeks of diet manipulation; B) 48 h after training in Experiment 2 (grey shading), during which mice underwent six weeks of diet manipulation; and C) four weeks after training in Experiment 3, during which mice underwent six total weeks of diet manipulation (training occurred after two weeks of diet manipulation). Experiment 1: 0.4% NaCl females, n=9; 4.0% NaCl females, n=9; 0.4% NaCl males, n=8; 4.0% NaCl males, n=8. Experiment 2 (grey shading): 0.4% NaCl females, n=8; 4.0% NaCl females, n=8; 0.4% NaCl males, n=9; 4.0% NaCl males, n=9. Experiment 3: 0.4% NaCl females, n=8; 4.0% NaCl females, n=8; 0.4% NaCl males, n=7; 4.0% NaCl males, n=8. Data are graphed as mean ± 95% confidence interval; pairwise comparisons were made using Bonferroni correction. Thick symbol borders on female data indicate significant ($p<0.05$) differences between females and males for that specific diet. *$p<0.05$ indicates significant differences between females on 0.4% versus 4.0% NaCl.

**S7 Table. Three-way ANOVAs on context fear expression (min 2-6) for context fear conditioned mice across Experiment.**

| | Context Trained Shock Groups | | |
|---|---|---|---|
| **Context Fear Expression** | **Experiment 1** | **Experiment 2** | **Experiment 3** |
| Sex | $F(1,57)=7.538$<br>**p=0.008**<br>partial $\eta^2$=**0.117** | $F(1,60)=22.29$<br>$p<0.001$<br>partial $\eta^2=0.271$ | $F(1,57)=15.19$<br>$p<0.001$<br>partial $\eta^2=0.210$ |
| Diet | $F(1,57)=2.107$<br>$p=0.152$<br>partial $\eta^2=0.036$ | $F(1,60)=6.665$<br>**p=0.012**<br>partial $\eta^2$=**0.100** | $F(1,57)=0.002$<br>$p=0.962$<br>partial $\eta^2=0.000$ |
| Context | $F(1,57)=114.1$<br>**p<0.001**<br>partial $\eta^2$=**0.667** | $F(1,60)=171.5$<br>$p<0.001$<br>partial $\eta^2=0.741$ | $F(1,57)=84.47$<br>$p<0.001$<br>partial $\eta^2=0.597$ |
| Sex × Diet | $F(1,57)=0.191$<br>$p=0.663$<br>partial $\eta^2=0.003$ | $F(1,60)=0.099$<br>$p=0.754$<br>partial $\eta^2=0.002$ | $F(1,57)=5.380$<br>**p=0.024**<br>partial $\eta^2$=**0.086** |
| Sex × Context | $F(1,57)=1.297$<br>$p=0.259$<br>partial $\eta^2=0.022$ | $F(1,60)=4.938$<br>**p=0.030**<br>partial $\eta^2$=**0.076** | $F(1,57)=9.414$<br>**p=0.003**<br>partial $\eta^2$=**0.142** |
| Diet × Context | $F(1,57)=0.486$<br>$p=0.489$<br>partial $\eta^2=0.008$ | $F(1,60)=0.316$<br>$p=0.576$<br>partial $\eta^2=0.005$ | $F(1,57)=0.014$<br>$p=0.905$<br>partial $\eta^2=0.000$ |
| Sex × Diet × Context | $F(1,57)=3.235$<br>$p=0.077$<br>partial $\eta^2=0.054$ | $F(1,60)=1.682$<br>$p=0.200$<br>partial $\eta^2=0.027$ | $F(1,57)=2.642$<br>$p=0.110$<br>partial $\eta^2=0.044$ |

**S8 Table. Three-way repeated measures ANOVAs on full 10 min time course of context fear testing for mice of both sexes in Experiment 1.**

S8A Table

| **Females** | **Experiment 1 – Context Fear Testing** | | |
|---|---|---|---|
| Diet | $F(1,29)=0.539$ | $p=0.469$ | partial $\eta^2=0.018$ |
| Context | $F(1,29)=56.67$ | $p<0.001$ | partial $\eta^2=0.661$ |
| Time | $F(8.41,244.0)=7.535$ | $p<0.001$ | partial $\eta^2=0.206$ |
| Time × Diet | $F(8.41,244.0)=0.675$ | $p=0.721$ | partial $\eta^2=0.023$ |
| Time × Context | $F(8.41,244.0)=2.142$ | **$p=0.030$** | partial $\eta^2=$**0.069** |
| Diet × Context | $F(1,29)=0.491$ | $p=0.489$ | partial $\eta^2=0.017$ |
| Time × Diet × Context | $F(8.41,244.0)=0.771$ | $p=0.635$ | partial $\eta^2=0.026$ |

S8B Table

| **Males** | **Experiment 1 – Context Fear Testing** | | |
|---|---|---|---|
| Diet | $F(1,28)=2.622$ | $p=0.117$ | partial $\eta^2=0.086$ |
| Context | $F(1,28)=39.04$ | $p=0.857$ | partial $\eta^2=0.582$ |
| Time | $F(7.48,209.4)=2.649$ | $p=0.010$ | partial $\eta^2=0.086$ |
| Time × Diet | $F(7.48,209.4)=0.509$ | $p=0.838$ | partial $\eta^2=0.018$ |
| Time × Context | $F(7.48,209.4)=3.909$ | **$p<0.001$** | partial $\eta^2=$**0.122** |
| Diet × Context | $F(1,28)=3.371$ | $p=0.077$ | partial $\eta^2=0.107$ |
| Time × Diet × Context | $F(7.48,209.4)=0.772$ | $p=0.620$ | partial $\eta^2=0.027$ |

**S9 Table. Three-way repeated measures ANOVAs on full 10 min time course of context fear testing for mice of both sexes in Experiment 2.**

S9A Table

| **Females** | **Experiment 2 – Context Fear Testing** |
|---|---|
| Diet | $F(1,29)=0.858$ $p=0.362$ partial $\eta^2=0.029$ |
| Context | $F(1,29)=109.8$ $\mathbf{p<0.001}$ partial $\eta^2=\mathbf{0.791}$ |
| Time | $F(8.06,233.7)=4.180$ $\mathbf{p<0.001}$ partial $\eta^2=\mathbf{0.126}$ |
| Time × Diet | $F(8.06,233.7)=0.873$ $p=0.541$ partial $\eta^2=0.029$ |
| Time × Context | $F(8.06,233.7)=1.429$ $p=0.184$ partial $\eta^2=0.047$ |
| Diet × Context | $F(1,29)=0.363$ $p=0.552$ partial $\eta^2=0.012$ |
| Time × Diet × Context | $F(8.06,233.7)=0.739$ $p=0.658$ partial $\eta^2=0.025$ |

S9B Table

| **Males** | **Experiment 2 – Context Fear Testing** |
|---|---|
| Diet | $F(1,31)=3.042$ $p=0.091$ partial $\eta^2=0.089$ |
| Context | $F(1,31)=52.22$ $p<0.001$ partial $\eta^2=0.628$ |
| Time | $F(6.86,212.6)=6.425$ $p<0.001$ partial $\eta^2=0.172$ |
| Time × Diet | $F(6.86,212.6)=1.542$ $p=0.066$ partial $\eta^2=0.047$ |
| Time × Context | $F(6.86,212.6)=3.902$ $\mathbf{p<0.001}$ partial $\eta^2=\mathbf{0.112}$ |
| Diet × Context | $F(1,31)=1.344$ $p=0.255$ partial $\eta^2=0.042$ |
| Time × Diet × Context | $F(6.86,212.6)=1.188$ $p=0.312$ partial $\eta^2=0.037$ |

**S10 Table. Three-way repeated measures ANOVAs on full 10 min time course of context fear testing for mice of both sexes in Experiment 3.**

S10A Table

| **Females** | **Experiment 3 – Context Fear Testing** | | |
|---|---|---|---|
| Diet | $F(1,30)=0.646$ | $p=0.428$ | partial $\eta^2=0.021$ |
| Context | $F(1,30)=15.64$ | **$p<0.001$** | partial $\eta^2=$**$0.343$** |
| Time | $F(6.57,197.0)=5.706$ | **$p<0.001$** | partial $\eta^2=$**$0.160$** |
| Time × Diet | $F(6.57,197.0)=0.862$ | $p=0.532$ | partial $\eta^2=0.028$ |
| Time × Context | $F(6.57,197.0)=1.029$ | $p=0.410$ | partial $\eta^2=0.033$ |
| Diet × Context | $F(1,30)=1.045$ | $p=0.315$ | partial $\eta^2=0.034$ |
| Time × Diet × Context | $F(6.57,197.0)=0.614$ | $p=0.733$ | partial $\eta^2=0.020$ |

S10B Table

| **Males** | **Experiment 3 – Context Fear Testing** | | |
|---|---|---|---|
| Diet | $F(1,27)=6.657$ | **$p=0.016$** | partial $\eta^2=$**$0.198$** |
| Context | $F(1,27)=109.0$ | **$p<0.001$** | partial $\eta^2=$**$0.801$** |
| Time | $F(6.91,186.7)=2.821$ | **$p=0.008$** | partial $\eta^2=$**$0.095$** |
| Time × Diet | $F(6.91,186.7)=0.980$ | $p=0.447$ | partial $\eta^2=0.035$ |
| Time × Context | $F(6.91,186.7)=1.150$ | $p=0.334$ | partial $\eta^2=0.041$ |
| Diet × Context | $F(1,27)=0.476$ | $p=0.496$ | partial $\eta^2=0.017$ |
| Time × Diet × Context | $F(6.91,186.7)=1.474$ | $p=0.180$ | partial $\eta^2=0.052$ |

S4 Figure

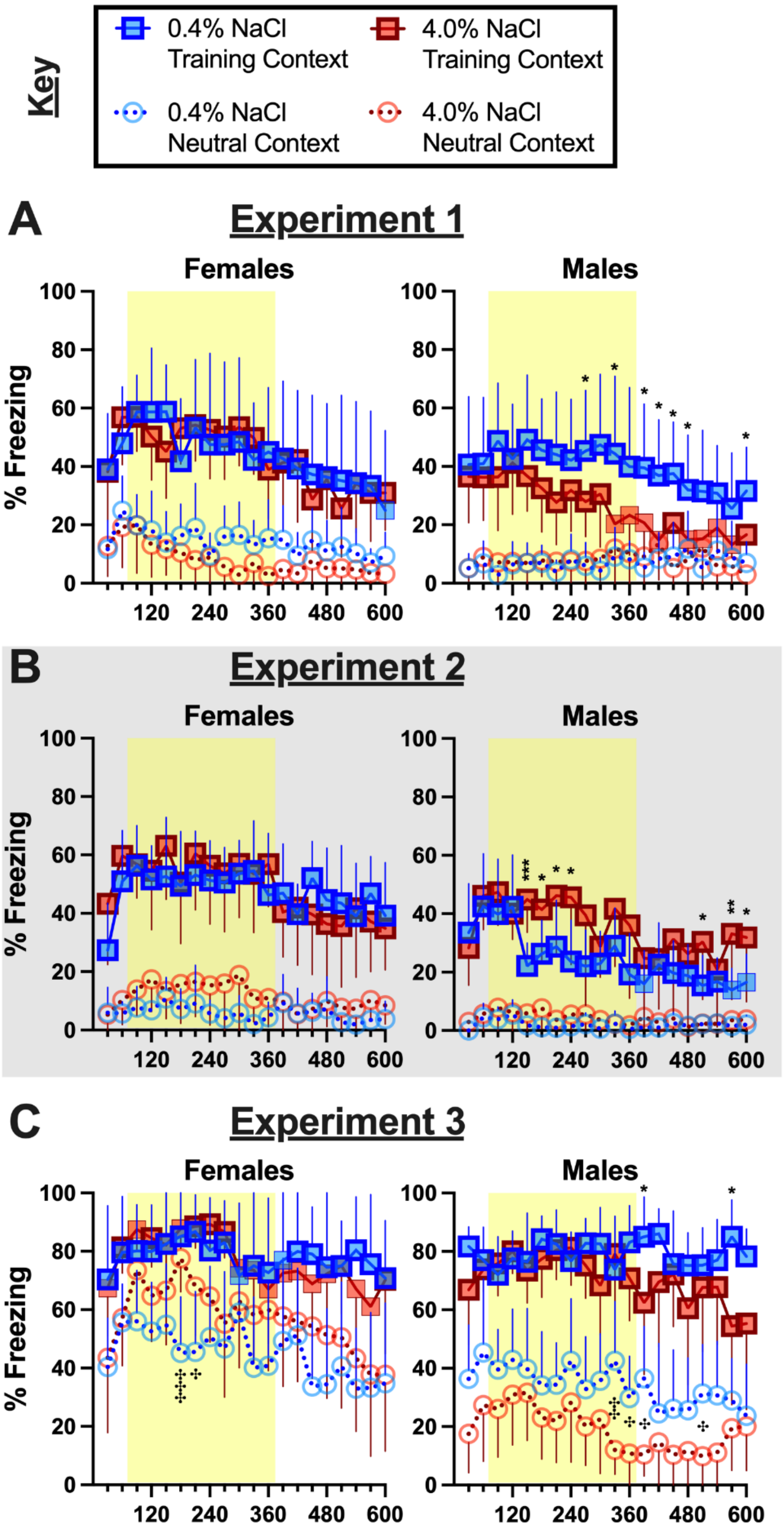

Key
0.4% NaCl Training Context
4.0% NaCl Training Context
0.4% NaCl Neutral Context
4.0% NaCl Neutral Context
A
Experiment 1
B
Experiment 2
C
Experiment 3
Females
Males
% Freezing
0
20
40
60
80
100
120
240
360
480
600

**S4 Fig. Time course of context fear expression testing in shock groups**. Mice assigned to 0.4% NaCl represented by blue symbols, mice assigned to 4.0% NaCl represented by red symbols; mice tested in Training Context represented by squares and solid lines, mice tested in Neutral Context represented by circles and dotted lines. Percent freezing for each 30 s bin of the 10 min testing session is graphed for all mice. Yellow shading indicates minutes 2 through 6, which were averaged and analyzed as our original measure of context fear expression. Testing occurred A) 48 h after training in Experiment 1, during which mice underwent two weeks of diet manipulation; B) 48 h after training in Experiment 2 (grey shading), during which mice underwent six weeks of diet manipulation; and C) four weeks after training in Experiment 3, during which mice underwent six total weeks of diet manipulation (training occurred after two weeks of diet manipulation). Experiment 1: 0.4% NaCl females Training Context, n=7; 0.4% NaCl females Neutral Context, n=9; 4.0% NaCl females Training Context, n=8; 4.0% NaCl females Neutral Context, n=9; 0.4% NaCl males Training Context, n=8; 0.4% NaCl males Neutral Context, n=9; 4.0% NaCl males Training Context, n=7; 4.0% NaCl males Neutral Context, n=8. Experiment 2: 0.4% NaCl females Training Context, n=9; 0.4% NaCl females Neutral Context, n=8; 4.0% NaCl females Training Context, n=8; 4.0% NaCl females Neutral Context, n=8; 0.4% NaCl males Training Context, n=9; 0.4% NaCl males Neutral Context, n=8; 4.0% NaCl males Training Context, n=8; 4.0% NaCl males Neutral Context, n=10. Experiment 3: 0.4% NaCl females Training Context, n=8; 0.4% NaCl females Neutral Context, n=9; 4.0% NaCl females Training Context, n=9; 4.0% NaCl females Neutral Context, n=8; 0.4% NaCl males Training Context, n=7; 0.4% NaCl males Neutral Context, n=8; 4.0% NaCl males Training Context, n=8; 4.0% NaCl males Neutral Context, n=8. Data are graphed as mean ± 95% confidence interval; pairwise comparisons were made using Bonferroni correction. Thick symbol borders on Training Context data indicate significant ($p<0.05$) differences between Training and Neutral Contexts for that specific diet/sex combination. *$p<0.05$, **$p<0.01$, ***$p<0.001$ indicate difference between 0.4% and 4.0% NaCl within Training Context and same sex. $^{\ddagger}p<0.05$, $^{\ddagger\ddagger}p<0.01$, $^{\ddagger\ddagger\ddagger}p<0.001$ indicate difference between 0.4% and 4.0% NaCl within Neutral Context and same sex.

**S11 Table. Three-way ANOVAs on serum osmolality in control no shock mice across Experiments.**

| Osmolality | No Shock Groups Across Experiments | | |
|---|---|---|---|
| Sex | $F(1,89)=0.424$ | $p=0.517$ | partial $\eta^2=0.005$ |
| Diet | $F(1,89)=0.046$ | $p=0.830$ | partial $\eta^2=0.001$ |
| Experiment | $F(1,89)=3.639$ | $p=0.030$ | partial $\eta^2=0.076$ |
| Sex × Diet | $F(1,89)=0.510$ | $p=0.477$ | partial $\eta^2=0.006$ |
| Sex × Experiment | $F(1,89)=4.287$ | **$p=0.017$** | partial $\eta^2=$**0.088** |
| Diet × Experiment | $F(1,89)=1.729$ | $p=0.183$ | partial $\eta^2=0.037$ |
| Sex × Diet × Experiment | $F(1,89)=1.331$ | $p=0.270$ | partial $\eta^2=0.029$ |

S5 Figure

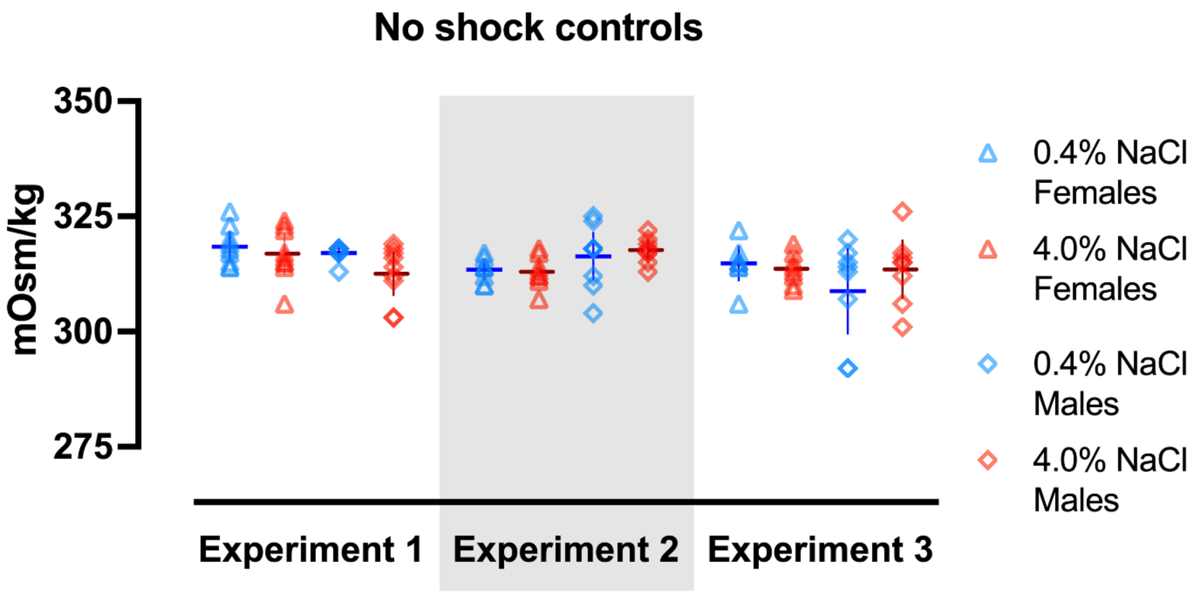


**S5 Fig. Serum osmolality in no shock control mice across Experiments**.

Females represented by triangles, males by diamonds; 0.4% NaCl represented by blue symbols, 4.0% NaCl represented by red symbols. Experiment 1: 0.4% NaCl females, n=9; 4.0% NaCl females, n=9; 0.4% NaCl males, n=8; 4.0% NaCl males, n=9. Experiment 2 (grey shading): 0.4% NaCl females, n=8; 4.0% NaCl females, n=8; 0.4% NaCl males, n=9; 4.0% NaCl males, n=9. Experiment 3: 0.4% NaCl females, n=8; 4.0% NaCl females, n=8; 0.4% NaCl males, n=8; 4.0% NaCl males, n=8. Data are graphed as mean ± 95% confidence interval.

**S12 Table. Three-way ANOVAs on serum osmolality in context fear conditioned mice across Experiments.**

| | Context Trained Shock Groups | | |
|---|---|---|---|
| **Osmolality** | **Experiment 1** | **Experiment 2** | **Experiment 3** |
| Sex | $F(1,59)=1.662$<br>$p=0.202$<br>partial $\eta^2=0.027$ | $F(1,62)=0.096$<br>$p=0.758$<br>partial $\eta^2=0.002$ | $F(1,55)=0.086$<br>$p=0.771$<br>partial $\eta^2=0.002$ |
| Diet | $F(1,59)=001$<br>$p=0.979$<br>partial $\eta^2=0.000$ | $F(1,62)=0.673$<br>$p=0.415$<br>partial $\eta^2=0.011$ | $F(1,55)=0.745$<br>$p=0.392$<br>partial $\eta^2=0.013$ |
| Context | $F(1,59)=1.339$<br>$p=0.252$<br>partial $\eta^2=0.022$ | $F(1,62)=0.005$<br>$p=0.941$<br>partial $\eta^2=0.000$ | $F(1,55)=1.440$<br>$p=0.235$<br>partial $\eta^2=0.026$ |
| Sex × Diet | $F(1,59)=0.012$<br>$p=0.912$<br>partial $\eta^2=0.000$ | $F(1,62)=0.152$<br>$p=0.698$<br>partial $\eta^2=0.002$ | $F(1,55)=0.213$<br>$p=0.646$<br>partial $\eta^2=0.004$ |
| Sex × Context | $F(1,59)=0.482$<br>$p=0.490$<br>partial $\eta^2=0.008$ | $F(1,62)=0.020$<br>$p=0.888$<br>partial $\eta^2=0.000$ | $F(1,55)=0.154$<br>$p=0.696$<br>partial $\eta^2=0.003$ |
| Diet × Context | $F(1,59)=1.903$<br>$p=0.173$<br>partial $\eta^2=0.031$ | $F(1,62)=0.055$<br>$p=0.815$<br>partial $\eta^2=0.001$ | $F(1,55)=0.781$<br>$p=0.381$<br>partial $\eta^2=0.014$ |
| Sex × Diet × Context | $F(1,59)=0.003$<br>$p=0.959$<br>partial $\eta^2=0.000$ | $F(1,62)=1.901$<br>$p=0.173$<br>partial $\eta^2=0.030$ | $F(1,55)=0.856$<br>$p=0.359$<br>partial $\eta^2=0.015$ |

**S13. Three-way ANOVAs on log-transformed serum corticosterone levels in control no shock mice across Experiments.**

| Corticosterone | No Shock Groups Across Experiments | | |
|---|---|---|---|
| Sex | $F(1,81)=12.77$ | $p<0.001$ | partial $\eta^2=0.136$ |
| Diet | $F(1,81)=2.498$ | $p=0.118$ | partial $\eta^2=0.030$ |
| Experiment | $F(2,81)=12.81$ | $p<0.001$ | partial $\eta^2=0.240$ |
| Sex × Diet | $F(1,81)=1.654$ | $p=0.202$ | partial $\eta^2=0.020$ |
| Sex × Experiment | $F(2,81)=3.498$ | **$p=0.035$** | partial $\eta^2=$**0.080** |
| Diet × Experiment | $F(2,81)=0.527$ | $p=0.592$ | partial $\eta^2=0.013$ |
| Sex × Diet × Experiment | $F(2,81)=1.512$ | $p=0.227$ | partial $\eta^2=0.036$ |

S6 Figure

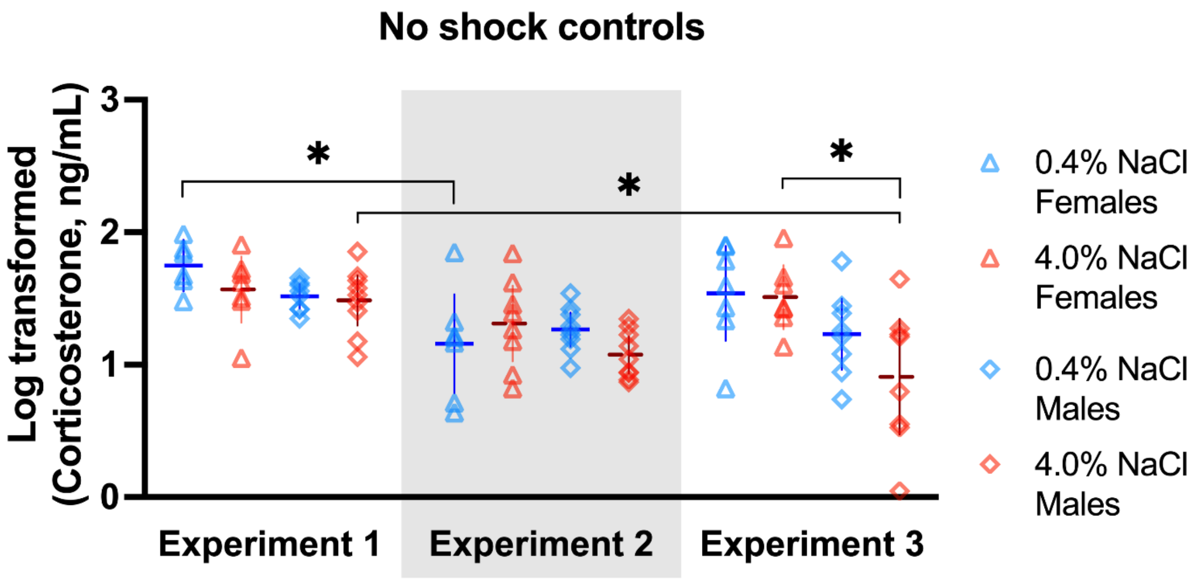


**S6 Fig. Log-transformed serum corticosterone levels of no shock control mice across Experiments**.

Females represented by triangles, males by diamonds; 0.4% NaCl represented by blue symbols, 4.0% NaCl represented by red symbols. Experiment 1: 0.4% NaCl females, n=6; 4.0% NaCl females, n=7; 0.4% NaCl males, n=8; 4.0% NaCl males, n=9. Experiment 2 (grey shading): 0.4% NaCl females, n=7; 4.0% NaCl females, n=8; 0.4% NaCl males, n=9; 4.0% NaCl males, n=9. Experiment 3: 0.4% NaCl females, n=7; 4.0% NaCl females, n=7; 0.4% NaCl males, n=8; 4.0% NaCl males, n=8. Data are graphed as mean ± 95% confidence interval. *p<0.05 indicate difference between marked groups within same diet.

**S14 Table. Three-way ANOVAs on log-transformed serum corticosterone in context fear conditioned mice across Experiments.**

| | Context Trained Shock Groups | | |
|---|---|---|---|
| **Corticosterone** | **Experiment 1** | **Experiment 2** | **Experiment 3** |
| Sex | $F(1,52)=5.924$<br>**p=0.018**<br>partial $\eta^2$=**0.102** | $F(1,59)=0.973$<br>$p=0.328$<br>partial $\eta^2=0.016$ | $F(1,53)=38.87$<br>**p<0.001**<br>partial $\eta^2$=**0.423** |
| Diet | $F(1,52)=1.107$<br>$p=0.298$<br>partial $\eta^2=0.021$ | $F(1,59)=1.540$<br>$p=0.220$<br>partial $\eta^2=0.025$ | $F(1,53)=1.170$<br>$p=0.284$<br>partial $\eta^2=0.022$ |
| Context | $F(1,52)=0.325$<br>$p=0.571$<br>partial $\eta^2=0.006$ | $F(1,59)=5.074$<br>**p=0.028**<br>partial $\eta^2$=**0.079** | $F(1,53)=0.000$<br>$p=0.992$<br>partial $\eta^2=0.000$ |
| Sex × Diet | $F(1,52)=2.815$<br>$p=0.099$<br>partial $\eta^2=0.051$ | $F(1,59)=1.041$<br>$p=0.312$<br>partial $\eta^2=0.017$ | $F(1,53)=0.186$<br>$p=0.668$<br>partial $\eta^2=0.003$ |
| Sex × Context | $F(1,52)=2.355$<br>$p=0.131$<br>partial $\eta^2=0.043$ | $F(1,59)=19.69$<br>$p=0.166$<br>partial $\eta^2=0.032$ | $F(1,53)=0.012$<br>$p=0.913$<br>partial $\eta^2=0.000$ |
| Diet × Context | $F(1,52)=0.033$<br>$p=0.856$<br>partial $\eta^2=0.001$ | $F(1,59)=0.056$<br>$p=0.814$<br>partial $\eta^2=0.001$ | $F(1,53)=0.966$<br>$p=0.330$<br>partial $\eta^2=0.018$ |
| Sex × Diet × Context | $F(1,52)=0.164$<br>$p=0.687$<br>partial $\eta^2=0.003$ | $F(1,59)=0.155$<br>$p=0.695$<br>partial $\eta^2=0.003$ | $F(1,53)=0.204$<br>$p=0.654$<br>partial $\eta^2=0.004$ |

S7 Figure

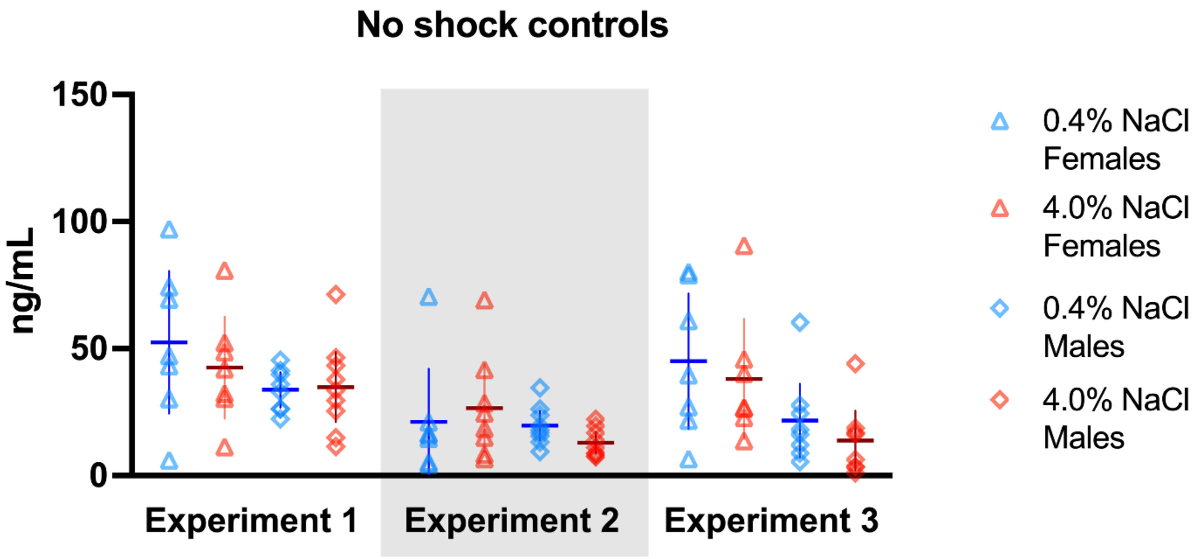


**S7 Fig. Raw (pre-transformed) serum corticosterone levels of no shock control mice across Experiments**.

Females represented by triangles, males by diamonds; 0.4% NaCl represented by blue symbols, 4.0% NaCl represented by red symbols. Experiment 1: 0.4% NaCl females, n=6; 4.0% NaCl females, n=7; 0.4% NaCl males, n=8; 4.0% NaCl males, n=9. Experiment 2 (grey shading): 0.4% NaCl females, n=7; 4.0% NaCl females, n=8; 0.4% NaCl males, n=9; 4.0% NaCl males, n=9. Experiment 3: 0.4% NaCl females, n=7; 4.0% NaCl females, n=7; 0.4% NaCl males, n=8; 4.0% NaCl males, n=8. Data are graphed as mean ± 95% confidence interval. These data were not statistically analyzed – log transformations were applied to normalize data distribution prior to analyzing statistically.

S8 Figure

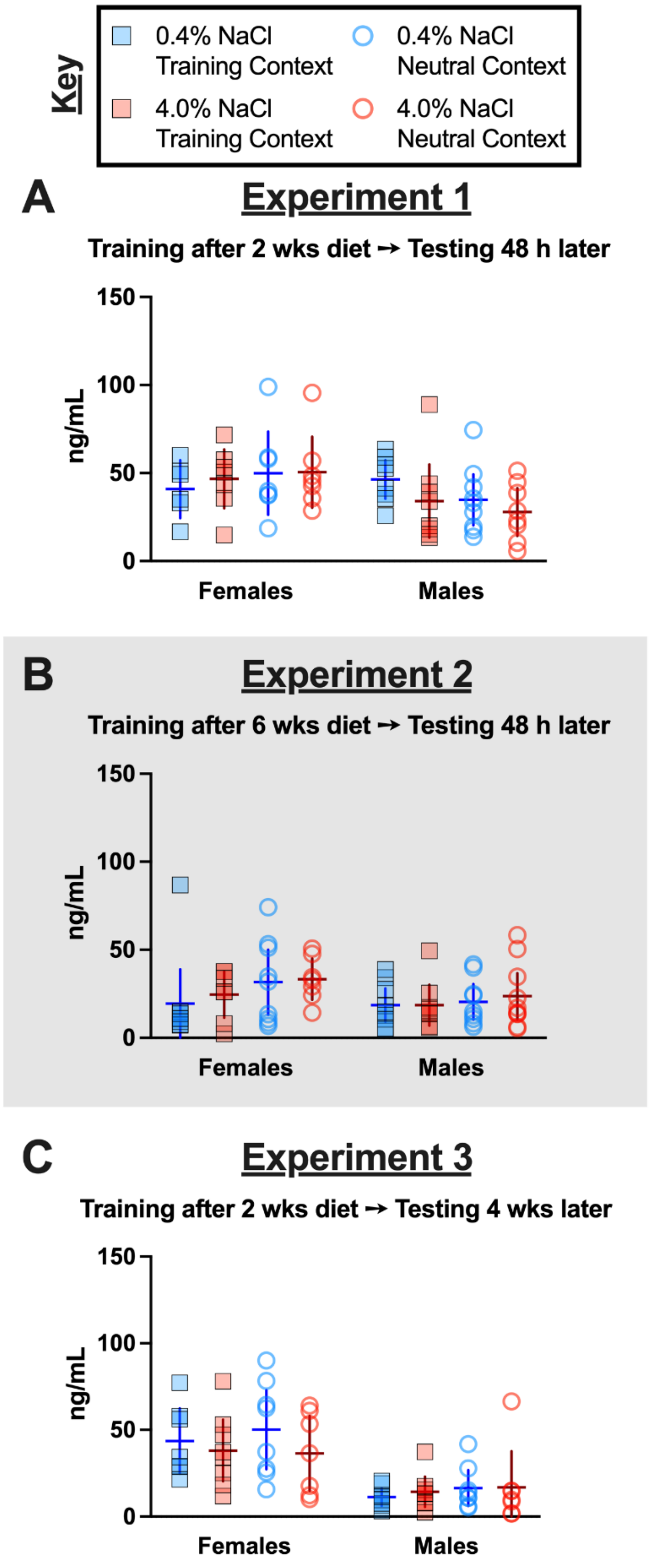
Key
0.4% NaCl Training Context
0.4% NaCl Neutral Context
4.0% NaCl Training Context
4.0% NaCl Neutral Context
A
Experiment 1
Training after 2 wks diet → Testing 48 h later
ng/mL
150
100
50
0
Females
Males
B
Experiment 2
Training after 6 wks diet → Testing 48 h later
ng/mL
150
100
50
0
Females
Males
C
Experiment 3
Training after 2 wks diet → Testing 4 wks later
ng/mL
150
100
50
0
Females
Males

**S8 Fig. Raw (pre-transformed) serum corticosterone levels in fear conditioned mice across Experiments**.

Mice assigned to 0.4% NaCl represented by blue symbols, mice assigned to 4.0% NaCl represented by red symbols; mice tested in Training Context represented by squares and solid lines, mice tested in Neutral Context represented by circles and dotted lines. Log-transformed serum corticosterone levels in context fear conditioned mice in A) Experiment 1, B) Experiment 2 (grey shading), and C) Experiment 3. Experiment 1: 0.4% NaCl females Training Context, n=6; 0.4% NaCl females Neutral Context, n=7; 4.0% NaCl females Training Context, n=7; 4.0% NaCl females Neutral Context, n=7; 0.4% NaCl males Training Context, n=8; 0.4% NaCl males Neutral Context, n=9; 4.0% NaCl males Training Context, n=8; 4.0% NaCl males Neutral Context, n=8. Experiment 2: 0.4% NaCl females Training Context, n=8; 0.4% NaCl females Neutral Context, n=9; 4.0% NaCl females Training Context, n=7; 4.0% NaCl females Neutral Context, n=7; 0.4% NaCl males Training Context, n=9; 0.4% NaCl males Neutral Context, n=9; 4.0% NaCl males Training Context, n=8; 4.0% NaCl males Neutral Context, n=10. Experiment 3: 0.4% NaCl females Training Context, n=7; 0.4% NaCl females Neutral Context, n=8; 4.0% NaCl females Training Context, n=8; 4.0% NaCl females Neutral Context, n=7; 0.4% NaCl males Training Context, n=8; 0.4% NaCl males Neutral Context, n=8; 4.0% NaCl males Training Context, n=8; 4.0% NaCl males Neutral Context, n=7. Data are graphed as mean ± 95% confidence interval. These data were not statistically analyzed – log transformations were applied to normalize data distribution prior to analyzing statistically.

**S15 Table. Three-way repeated measures ANOVAs on weekly average water consumption per day for context fear conditioned mice across Experiments.**

S15A Table

| Females | Experiment 1 – Water/day | | |
|---|---|---|---|
| Diet | F(1,30)=50.89 | **p<0.001** | partial $\eta^2$=**0.629** |
| Context | F(1,30)=0.361 | p=0.553 | partial $\eta^2$=0.012 |
| Time | F(1.64,49.08)=0.680 | p=0.483 | partial $\eta^2$=0.022 |
| Time × Diet | F(1.64,49.08)=0.225 | p=0.755 | partial $\eta^2$=0.007 |
| Time × Context | F(1.64,49.08)=0.011 | p=0.977 | partial $\eta^2$=0.000 |
| Diet × Context | F(1,30)=0.000 | p=0.985 | partial $\eta^2$=0.000 |
| Time × Diet × Context | F(1.64,49.08)=0.613 | p=0.514 | partial $\eta^2$=0.020 |

S15B Table

| Males | Experiment 1 – Water/day | | |
|---|---|---|---|
| Diet | F(1,29)=13.40 | **p<0.001** | partial $\eta^2$=**0.316** |
| Context | F(1,29)=1.042 | p=0.316 | partial $\eta^2$=0.035 |
| Time | F(1.27,36.92)=9.485 | **p=0.002** | partial $\eta^2$=**0.246** |
| Time × Diet | F(1.27,36.92)=2.030 | p=0.159 | partial $\eta^2$=0.065 |
| Time × Context | F(1.27,36.92)=0.411 | p=0.574 | partial $\eta^2$=0.014 |
| Diet × Context | F(1,29)=1.036 | p=0.317 | partial $\eta^2$=0.035 |
| Time × Diet × Context | F(1.27,36.92)=0.595 | p=0.484 | partial $\eta^2$=0.020 |

S15C Table

| Females | Experiment 2 – Water/day | | |
|---|---|---|---|
| Diet | F(1,30)=312.2 | p<0.001 | partial $\eta^2$=0.912 |
| Context | F(1,30)=0.026 | p=0.874 | partial $\eta^2$=0.001 |
| Time | F(4.11,123.3)=4.981 | p<0.001 | partial $\eta^2$=0.142 |
| Time × Diet | F(4.11,123.3)=5.091 | **p<0.001** | partial $\eta^2$=**0.145** |
| Time × Context | F(4.11,123.3)=1.342 | p=0.258 | partial $\eta^2$=0.043 |
| Diet × Context | F(1,30)=3.278 | p=0.080 | partial $\eta^2$=0.099 |
| Time × Diet × Context | F(4.11,123.3)=0.991 | p=0.417 | partial $\eta^2$=0.032 |

S15D Table

| Males | Experiment 2 – Water/day | | |
|---|---|---|---|
| Diet | F(1,32)=41.90 | p<0.001 | partial $\eta^2$=0.567 |
| Context | F(1,32)=0.533 | p=0.471 | partial $\eta^2$=0.016 |
| Time | F(3.25,103.9)=17.41 | p<0.001 | partial $\eta^2$=0.352 |

| | | | |
|---|---|---|---|
| Time × Diet | F(3.25,103.9)=3.360 | **p=0.019** | partial $\eta^2$=**0.095** |
| Time × Context | F(3.25,103.9)=0.454 | p=0.730 | partial $\eta^2$=0.014 |
| Diet × Context | F(1,32)=0.765 | p=0.388 | partial $\eta^2$=0.023 |
| Time × Diet × Context | F(3.25,103.9)=0.938 | p=0.431 | partial $\eta^2$=0.028 |

S15E Table

| **Females** | **Experiment 3 – Water/day** | | |
|---|---|---|---|
| Diet | F(1,30)=91.88 | **p<0.001** | partial $\eta^2$=**0.754** |
| Context | F(1,30)=0.009 | p=0.927 | partial $\eta^2$=0.000 |
| Time | F(3.16,94.64)=13.26 | **p<0.001** | partial $\eta^2$=**0.306** |
| Time × Diet | F(3.16,94.64)=1.647 | p=0.182 | partial $\eta^2$=0.052 |
| Time × Context | F(3.16,94.64)=0.249 | p=0.871 | partial $\eta^2$=0.008 |
| Diet × Context | F(1,30)=0.039 | p=0.845 | partial $\eta^2$=0.001 |
| Time × Diet × Context | F(3.16,94.64)=0.163 | p=0.928 | partial $\eta^2$=0.005 |

S15F Table

| **Males** | **Experiment 3 – Water/day** | | |
|---|---|---|---|
| Diet | F(1,27)=45.38 | **p<0.001** | partial $\eta^2$=**0.627** |
| Context | F(1,27)=0.201 | p=0.657 | partial $\eta^2$=0.007 |
| Time | F(3.73,100.8)=17.83 | **p<0.001** | partial $\eta^2$=**0.398** |
| Time × Diet | F(3.73,100.8)=2.515 | p=0.050 | partial $\eta^2$=0.085 |
| Time × Context | F(3.73,100.8)=0.952 | p=0.433 | partial $\eta^2$=0.034 |
| Diet × Context | F(1,27)=0.035 | p=0.853 | partial $\eta^2$=0.001 |
| Time × Diet × Context | F(3.73,100.8)=0.445 | p=0.763 | partial $\eta^2$=0.016 |

S9 Figure

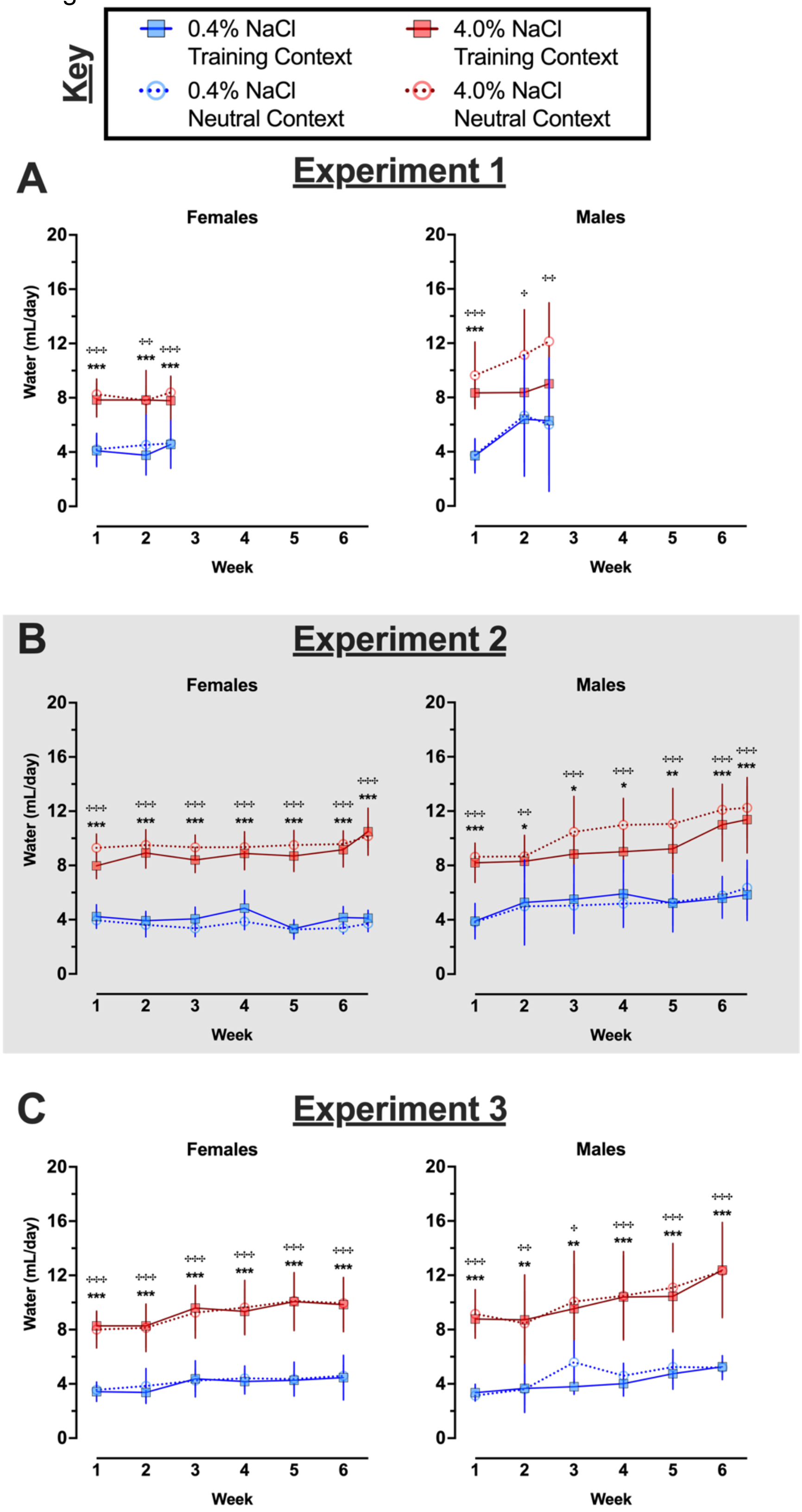

Key
0.4% NaCl Training Context
4.0% NaCl Training Context
0.4% NaCl Neutral Context
4.0% NaCl Neutral Context
A
Experiment 1
Females
Males
Water (mL/day)
Week
B
Experiment 2
Females
Males
Water (mL/day)
Week
C
Experiment 3
Females
Males
Water (mL/day)
Week

**S9 Fig. Average water consumed per day by context fear conditioned mice across Experiments.**

Mice assigned to 0.4% NaCl represented by blue symbols, mice assigned to 4.0% NaCl represented by red symbols; mice to be tested in Training Context represented by squares and solid lines, mice to be tested in Neutral Context represented by circles and dotted lines. Water consumption was measured twice weekly, and full weeks were averaged; partial weeks at the conclusion of A) Experiment 1 and B) Experiment 2 (grey shading) are included in the graphs. Some data loss occurred on the very last weighing day for a subset of animals in C) Experiment 3, thus graphs and repeated measures statistical analyses for Experiment 3 consumption cease at week 6 to maximize inclusion of mice in repeated measures analyses. Experiment 1: 0.4% NaCl females Training Context, n=8; 0.4% NaCl females Neutral Context, n=9; 4.0% NaCl females Training Context, n=8; 4.0% NaCl females Neutral Context, n=9; 0.4% NaCl males Training Context, n=8; 0.4% NaCl males Neutral Context, n=9; 4.0% NaCl males Training Context, n=8; 4.0% NaCl males Neutral Context, n=8. Experiment 2: 0.4% NaCl females Training Context, n=9; 0.4% NaCl females Neutral Context, n=9; 4.0% NaCl females Training Context, n=8; 4.0% NaCl females Neutral Context, n=8; 0.4% NaCl males Training Context, n=9; 0.4% NaCl males Neutral Context, n=9; 4.0% NaCl males Training Context, n=8; 4.0% NaCl males Neutral Context, n=10. Experiment 3: 0.4% NaCl females Training Context, n=8; 0.4% NaCl females Neutral Context, n=9; 4.0% NaCl females Training Context, n=9; 4.0% NaCl females Neutral Context, n=8; 0.4% NaCl males Training Context, n=8; 0.4% NaCl males Neutral Context, n=7; 4.0% NaCl males Training Context, n=8; 4.0% NaCl males Neutral Context, n=8. Data are graphed as mean ± 95% confidence interval. *$p<0.05$, **$p<0.01$, ***$p<0.001$ indicate difference between mice within the same sex consuming 0.4% NaCl versus 4.0% NaCl and tested in Training Context. $^{+}p<0.05$, $^{++}p<0.01$, $^{+++}p<0.001$ indicate difference

between mice within the same sex consuming 0.4% NaCl versus 4.0% NaCl and tested in Neutral Context.

**S16 Table. Three-way repeated measures ANOVAs on weekly average food consumption per day for context fear conditioned mice across Experiments.**

S16A Table

| Females | Experiment 1 – Food/day | | |
|---|---|---|---|
| Diet | $F(1,30)=6.785$ | **$p=0.014$** | partial $\eta^2$=**0.184** |
| Context | $F(1,30)=0.099$ | $p=0.756$ | partial $\eta^2=0.003$ |
| Time | $F(1.67,50.14)=0.650$ | $p=0.500$ | partial $\eta^2=0.021$ |
| Time × Diet | $F(1.67,50.14)=0.578$ | $p=0.535$ | partial $\eta^2=0.019$ |
| Time × Context | $F(1.67,50.14)=1.233$ | $p=0.295$ | partial $\eta^2=0.039$ |
| Diet × Context | $F(1,30)=0.237$ | $p=0.630$ | partial $\eta^2=0.008$ |
| Time × Diet × Context | $F(1.67,50.14)=0.809$ | $p=0.431$ | partial $\eta^2=0.026$ |

S16B Table

| Males | Experiment 1 – Food/day | | |
|---|---|---|---|
| Diet | $F(1,29)=1.427$ | $p=0.242$ | partial $\eta^2=0.047$ |
| Context | $F(1,29)=0.013$ | $p=0.910$ | partial $\eta^2=0.000$ |
| Time | $F(1.85,53.57)=1.825$ | $p=0.174$ | partial $\eta^2=0.059$ |
| Time × Diet | $F(1.85,53.57)=0.332$ | $p=0.702$ | partial $\eta^2=0.011$ |
| Time × Context | $F(1.85,53.57)=0.142$ | $p=0.852$ | partial $\eta^2=0.005$ |
| Diet × Context | $F(1,29)=0.003$ | $p=0.958$ | partial $\eta^2=0.000$ |
| Time × Diet × Context | $F(1.85,53.57)=0.866$ | $p=0.419$ | partial $\eta^2=0.029$ |

S16C Table

| Females | Experiment 2 – Food/day | | |
|---|---|---|---|
| Diet | $F(1,30)=41.57$ | **$p<0.001$** | partial $\eta^2$=**0.581** |
| Context | $F(1,30)=0.015$ | $p=0.904$ | partial $\eta^2=0.000$ |
| Time | $F(3.72,111.5)=13.02$ | **$p<0.001$** | partial $\eta^2$=**0.303** |
| Time × Diet | $F(3.72,111.5)=0.188$ | $p=0.936$ | partial $\eta^2=0.006$ |
| Time × Context | $F(3.72,111.5)=1.323$ | $p=0.267$ | partial $\eta^2=0.042$ |
| Diet × Context | $F(1,30)=0.032$ | $p=0.859$ | partial $\eta^2=0.001$ |
| Time × Diet × Context | $F(3.72,111.5)=0.433$ | $p=0.771$ | partial $\eta^2=0.014$ |

S16D Table

| Males | Experiment 2 – Food/day | | |
|---|---|---|---|
| Diet | $F(1,32)=3.836$ | $p=0.059$ | partial $\eta^2=0.107$ |
| Context | $F(1,32)=0.128$ | $p=0.722$ | partial $\eta^2=0.004$ |
| Time | $F(4.32,138.3)=10.30$ | **$p<0.001$** | partial $\eta^2$=**0.243** |
| Time × Diet | $F(4.32,138.3)=1.317$ | $p=0.265$ | partial $\eta^2=0.040$ |
| Time × Context | $F(4.32,138.3)=0.451$ | $p=0.786$ | partial $\eta^2=0.014$ |

| | | | |
|---|---|---|---|
| Diet × Context | F(1,32)=0.672 | p=0.418 | partial $\eta^2$=0.021 |
| Time × Diet × Context | F(4.32,138.3)=0.773 | p=0.554 | partial $\eta^2$=0.024 |

S16E Table

| **Females** | **Experiment 3 – Food/day** | | |
|---|---|---|---|
| Diet | F(1,30)=16.65 | **p<0.001** | partial $\eta^2$=**0.357** |
| Context | F(1,30)=1.131 | p=0.296 | partial $\eta^2$=0.036 |
| Time | F(2.78,83.39)=5.015 | **p=0.004** | partial $\eta^2$=**0.143** |
| Time × Diet | F(2.78,83.39)=1.584 | p=0.202 | partial $\eta^2$=0.050 |
| Time × Context | F(2.78,83.39)=0.457 | p=0.699 | partial $\eta^2$=0.015 |
| Diet × Context | F(1,30)=0.022 | p=0.884 | partial $\eta^2$=0.001 |
| Time × Diet × Context | F(2.78,83.39)=0.436 | p=0.713 | partial $\eta^2$=0.014 |

S16F Table

| **Males** | **Experiment 3 – Food/day** | | |
|---|---|---|---|
| Diet | F(1,28)=7.673 | **p=0.010** | partial $\eta^2$=**0.215** |
| Context | F(1,28)=0.017 | p=0.897 | partial $\eta^2$=0.001 |
| Time | F(3.40,95.20)=5.087 | **p=0.002** | partial $\eta^2$=**0.154** |
| Time × Diet | F(3.40,95.20)=0.981 | p=0.412 | partial $\eta^2$=0.034 |
| Time × Context | F(3.40,95.20)=1.256 | p=0.294 | partial $\eta^2$=0.043 |
| Diet × Context | F(1,28)=0.482 | p=0.482 | partial $\eta^2$=0.017 |
| Time × Diet × Context | F(3.40,95.20)=0.546 | p=0.674 | partial $\eta^2$=0.019 |

S10 Figure

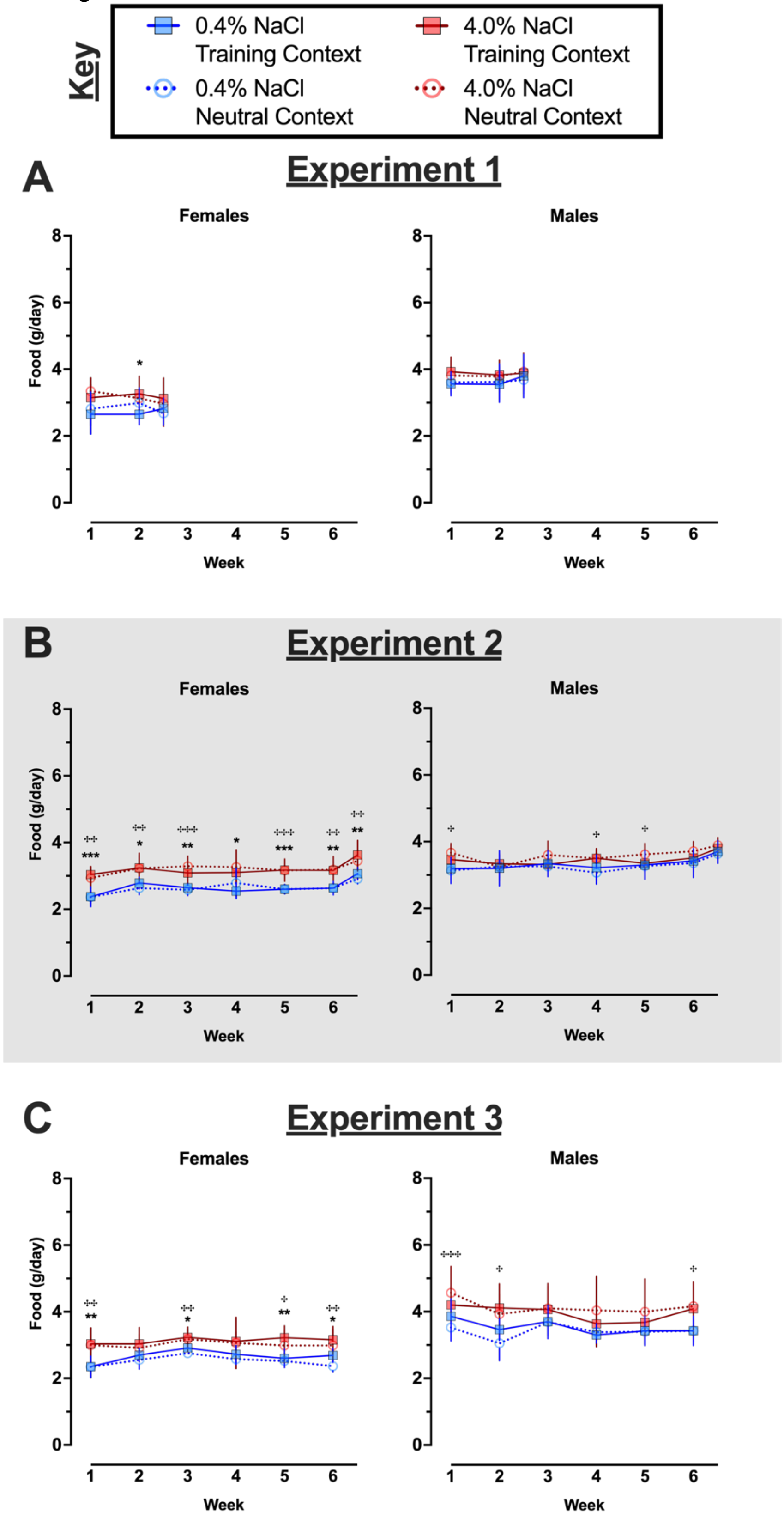

Key
0.4% NaCl Training Context
4.0% NaCl Training Context
0.4% NaCl Neutral Context
4.0% NaCl Neutral Context
A
Experiment 1
Females
Males
Food (g/day)
Week
B
Experiment 2
Females
Males
Food (g/day)
Week
C
Experiment 3
Females
Males
Food (g/day)
Week

**S10 Fig. Average food consumed per day by context fear conditioned mice across Experiments.**

Mice assigned to 0.4% NaCl represented by blue symbols, mice assigned to 4.0% NaCl represented by red symbols; mice to be tested in Training Context represented by squares and solid lines, mice to be tested in Neutral Context represented by circles and dotted lines. Food consumption was measured twice weekly, and full weeks were averaged; partial weeks at the conclusion of A) Experiment 1 and B) Experiment 2 (grey shading) are included in the graphs. Some data loss occurred on the very last weighing day for a subset of animals in C) Experiment 3, thus graphs and repeated measures statistical analyses for Experiment 3 consumption cease at week 6 to maximize inclusion of mice in repeated measures analyses. Experiment 1: 0.4% NaCl females Training Context, n=8; 0.4% NaCl females Neutral Context, n=9; 4.0% NaCl females Training Context, n=8; 4.0% NaCl females Neutral Context, n=9; 0.4% NaCl males Training Context, n=8; 0.4% NaCl males Neutral Context, n=9; 4.0% NaCl males Training Context, n=8; 4.0% NaCl males Neutral Context, n=8. Experiment 2: 0.4% NaCl females Training Context, n=9; 0.4% NaCl females Neutral Context, n=9; 4.0% NaCl females Training Context, n=8; 4.0% NaCl females Neutral Context, n=8; 0.4% NaCl males Training Context, n=9; 0.4% NaCl males Neutral Context, n=9; 4.0% NaCl males Training Context, n=8; 4.0% NaCl males Neutral Context, n=10. Experiment 3: 0.4% NaCl females Training Context, n=8; 0.4% NaCl females Neutral Context, n=9; 4.0% NaCl females Training Context, n=9; 4.0% NaCl females Neutral Context, n=8; 0.4% NaCl males Training Context, n=8; 0.4% NaCl males Neutral Context, n=8; 4.0% NaCl males Training Context, n=8; 4.0% NaCl males Neutral Context, n=8. Data are graphed as mean ± 95% confidence interval. *p<0.05, **p<0.01, ***p<0.001 indicate difference between mice within the same sex consuming 0.4% NaCl versus 4.0% NaCl and tested in Training Context. $^{+}$p<0.05, $^{++}$p<0.01, $^{+++}$p<0.001 indicate difference

between mice within the same sex consuming 0.4% NaCl versus 4.0% NaCl and tested in Neutral Context.

**S17 Table. Three-way repeated measures ANOVAs on weekly body weight change for context fear conditioned mice across Experiments.**

S17A Table

| Females | Experiment 1 – Body Weight Change | | |
|---|---|---|---|
| Diet | F(1,30)=1.586 | p=0.218 | partial $\eta^2$=0.050 |
| Context | F(1,30)=0.731 | p=0.399 | partial $\eta^2$=0.024 |
| Time | F(1.73,51.83)=82.14 | p<0.001 | partial $\eta^2$=0.732 |
| Time × Diet | F(1.73,51.83)=5.233 | **p=0.011** | partial $\eta^2$=**0.149** |
| Time × Context | F(1.73,51.83)=3.272 | p=0.053 | partial $\eta^2$=0.098 |
| Diet × Context | F(1,30)=0.675 | p=0.418 | partial $\eta^2$=0.022 |
| Time × Diet × Context | F(1.73,51.83)=0.437 | p=0.619 | partial $\eta^2$=0.014 |

S17B Table

| Males | Experiment 1 – Body Weight Change | | |
|---|---|---|---|
| Diet | F(1,29)=0.281 | p=0.600 | partial $\eta^2$=0.010 |
| Context | F(1,29)=0.998 | p=0.326 | partial $\eta^2$=0.033 |
| Time | F(1.86,53.91)=30.34 | p<0.001 | partial $\eta^2$=0.511 |
| Time × Diet | F(1.86,53.91)=0.330 | p=0.705 | partial $\eta^2$=0.011 |
| Time × Context | F(1.86,53.91)=0.308 | p=0.720 | partial $\eta^2$=0.011 |
| Diet × Context | F(1,29)=0.294 | p=0.592 | partial $\eta^2$=0.010 |
| Time × Diet × Context | F(1.86,53.91)=3.318 | **p=0.047** | partial $\eta^2$=**0.103** |

S17C Table

| Females | Experiment 2 – Body Weight Change | | |
|---|---|---|---|
| Diet | F(1,30)=6.764 | **p=0.014** | partial $\eta^2$=**0.184** |
| Context | F(1,30)=1.677 | p=0.205 | partial $\eta^2$=0.053 |
| Time | F(4.06,121.8)=60.14 | **p<0.001** | partial $\eta^2$=**0.667** |
| Time × Diet | F(4.06,121.8)=1.821 | p=0.128 | partial $\eta^2$=0.057 |
| Time × Context | F(4.06,121.8)=0.594 | p=0.670 | partial $\eta^2$=0.019 |
| Diet × Context | F(1,30)=0.216 | p=0.646 | partial $\eta^2$=0.007 |
| Time × Diet × Context | F(4.06,121.8)=0.017 | p=0.999 | partial $\eta^2$=0.001 |

S17D Table

| Males | Experiment 2 – Body Weight Change | | |
|---|---|---|---|
| Diet | F(1,32)=2.211 | p=0.147 | partial $\eta^2$=0.065 |
| Context | F(1,32)=0.806 | p=0.376 | partial $\eta^2$=0.025 |
| Time | F(4.63,148.3)=7.269 | **p<0.001** | partial $\eta^2$=**0.185** |
| Time × Diet | F(4.63,148.3)=2.292 | p=0.053 | partial $\eta^2$=0.067 |
| Time × Context | F(4.63,148.3)=0.580 | p=0.702 | partial $\eta^2$=0.018 |

| | | | |
|---|---|---|---|
| Diet × Context | F(1,32)=0.257 | p=0.616 | partial $\eta^2$=0.008 |
| Time × Diet × Context | F(4.63,148.3)=1.694 | p=0.145 | partial $\eta^2$=0.050 |

S17E Table

| Females | Experiment 3 – Body Weight Change | | |
|---|---|---|---|
| Diet | F(1,30)=1.695 | p=0.203 | partial $\eta^2$=0.053 |
| Context | F(1,30)=0.127 | p=0.724 | partial $\eta^2$=0.004 |
| Time | F(4.26,127.8)=74.03 | p<0.001 | partial $\eta^2$=0.712 |
| Time × Diet | F(4.26,127.8)=3.422 | **p=0.009** | partial $\eta^2$=**0.102** |
| Time × Context | F(4.26,127.8)=0.055 | p=0.996 | partial $\eta^2$=0.002 |
| Diet × Context | F(1,30)=1.156 | p=0.291 | partial $\eta^2$=0.037 |
| Time × Diet × Context | F(4.26,127.8)=2.173 | p=0.072 | partial $\eta^2$=0.068 |

S17F Table

| Males | Experiment 3 – Body Weight Change | | |
|---|---|---|---|
| Diet | F(1,28)=1.021 | p=0.321 | partial $\eta^2$=0.035 |
| Context | F(1,28)=0.231 | p=0.634 | partial $\eta^2$=0.008 |
| Time | F(3.86,107.9)=6.806 | **p<0.001** | partial $\eta^2$=**0.196** |
| Time × Diet | F(3.86,107.9)=1.575 | p=0.188 | partial $\eta^2$=0.053 |
| Time × Context | F(3.86,107.9)=1.271 | p=0.287 | partial $\eta^2$=0.043 |
| Diet × Context | F(1,28)=0.002 | p=0.962 | partial $\eta^2$=0.000 |
| Time × Diet × Context | F(3.86,107.9)=1.244 | p=0.297 | partial $\eta^2$=0.043 |

S11 Figure

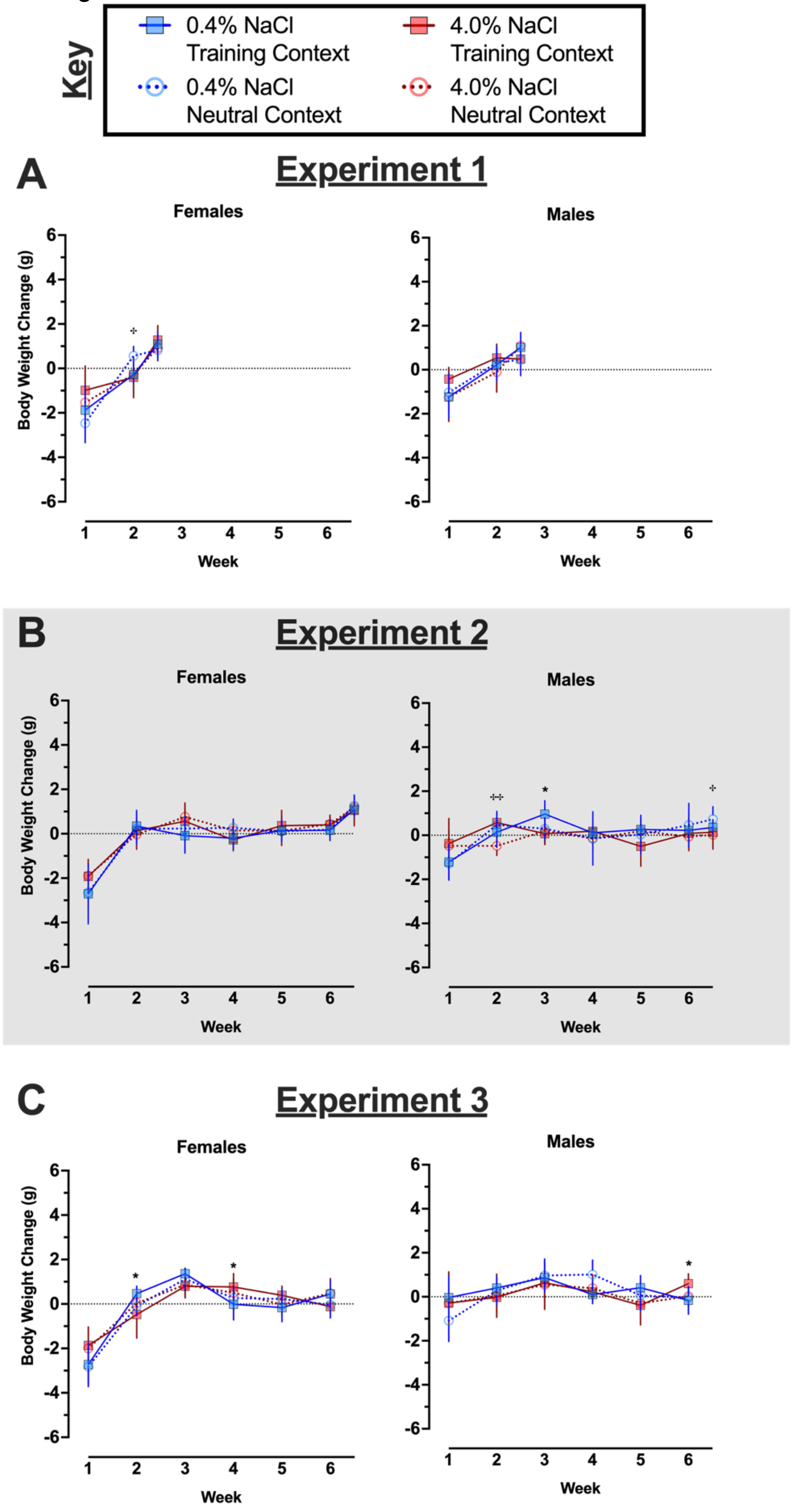

Key
0.4% NaCl Training Context
4.0% NaCl Training Context
0.4% NaCl Neutral Context
4.0% NaCl Neutral Context
A
Experiment 1
Females
Males
Body Weight Change (g)
Week
B
Experiment 2
Females
Males
Body Weight Change (g)
Week
C
Experiment 3
Females
Males
Body Weight Change (g)
Week

**S11 Fig. Weekly body weight change in context fear conditioned mice across Experiments.**

Mice assigned to 0.4% NaCl represented by blue symbols, mice assigned to 4.0% NaCl represented by red symbols; mice to be tested in Training Context represented by squares and solid lines, mice to be tested in Neutral Context represented by circles and dotted lines. Body weights were measured twice weekly, changes were calculated across full weeks. Body weight changes for partial weeks at the conclusion of A) Experiment 1 and B) Experiment 2 (grey shading) are included in the graphs. Some data loss occurred on the very last weighing day for a subset of animals in C) Experiment 3, thus graphs and repeated measures statistical analyses for Experiment 3 body weight changes cease at week 6 to maximize inclusion of mice in repeated measures analyses. Experiment 1: 0.4% NaCl females Training Context, n=8; 0.4% NaCl females Neutral Context, n=9; 4.0% NaCl females Training Context, n=8; 4.0% NaCl females Neutral Context, n=9; 0.4% NaCl males Training Context, n=8; 0.4% NaCl males Neutral Context, n=9; 4.0% NaCl males Training Context, n=8; 4.0% NaCl males Neutral Context, n=8. Experiment 2: 0.4% NaCl females Training Context, n=9; 0.4% NaCl females Neutral Context, n=9; 4.0% NaCl females Training Context, n=8; 4.0% NaCl females Neutral Context, n=8; 0.4% NaCl males Training Context, n=9; 0.4% NaCl males Neutral Context, n=9; 4.0% NaCl males Training Context, n=8; 4.0% NaCl males Neutral Context, n=10. Experiment 3: 0.4% NaCl females Training Context, n=8; 0.4% NaCl females Neutral Context, n=9; 4.0% NaCl females Training Context, n=9; 4.0% NaCl females Neutral Context, n=8; 0.4% NaCl males Training Context, n=8; 0.4% NaCl males Neutral Context, n=8; 4.0% NaCl males Training Context, n=8; 4.0% NaCl males Neutral Context, n=8. Data are graphed as mean ± 95% confidence interval. *$p<0.05$ indicates difference between mice within the same sex consuming 0.4% NaCl versus 4.0% NaCl and tested in Training Context. $^{+}p<0.05$, $^{++}p<0.01$, indicate difference between mice within the same sex consuming 0.4% NaCl versus 4.0% NaCl and tested in Neutral Context.

**S18 Table. Three-way repeated measures ANOVAs on twice weekly body weight measurements of context fear conditioned mice across Experiments.**

S18A Table

| **Females** | **Experiment 1 – Body Weight** | | |
|---|---|---|---|
| Diet | F(1,30)=0.388 | p=0.538 | partial $\eta^2$=0.013 |
| Context | F(1,30)=2.823 | p=0.103 | partial $\eta^2$=0.086 |
| Time | F(3.24,97.25)=30.28 | **p<0.001** | partial $\eta^2$=**0.502** |
| Time × Diet | F(3.24,97.25)=1.859 | p=0.137 | partial $\eta^2$=0.058 |
| Time × Context | F(3.24,97.25)=0.767 | p=0.524 | partial $\eta^2$=0.025 |
| Diet × Context | F(1,30)=0.015 | p=0.904 | partial $\eta^2$=0.000 |
| Time × Diet × Context | F(3.24,97.25)=1.035 | p=0.384 | partial $\eta^2$=0.033 |

S18B Table

| **Males** | **Experiment 1 – Body Weight** | | |
|---|---|---|---|
| Diet | F(1,29)=0.316 | p=0.579 | partial $\eta^2$=0.011 |
| Context | F(1,29)=0.061 | p=0.806 | partial $\eta^2$=0.002 |
| Time | F(2.43,70.56)=9.973 | **p<0.001** | partial $\eta^2$=**0.256** |
| Time × Diet | F(2.43,70.56)=0.357 | p=0.742 | partial $\eta^2$=0.012 |
| Time × Context | F(2.43,70.56)=1.979 | p=0.137 | partial $\eta^2$=0.064 |
| Diet × Context | F(1,29)=0.076 | p=0.785 | partial $\eta^2$=0.003 |
| Time × Diet × Context | F(2.43,70.56)=0.962 | p=0.401 | partial $\eta^2$=0.032 |

S18C Table

| **Females** | **Experiment 2 – Body Weight** | | |
|---|---|---|---|
| Diet | F(1,30)=0.074 | p=0.787 | partial $\eta^2$=0.002 |
| Context | F(1,30)=0.369 | p=0.548 | partial $\eta^2$=0.012 |
| Time | F(4.96,148.9)=40.23 | p<0.001 | partial $\eta^2$=0.573 |
| Time × Diet | F(4.96,148.9)=2.823 | **p=0.018** | partial $\eta^2$=**0.086** |
| Time × Context | F(4.96,148.9)=1.563 | p=0.175 | partial $\eta^2$=0.050 |
| Diet × Context | F(1,30)=1.463 | p=0.236 | partial $\eta^2$=0.046 |
| Time × Diet × Context | F(4.96,148.9)=0.356 | p=0.876 | partial $\eta^2$=0.012 |

S18D Table

| **Males** | **Experiment 2 – Body Weight** | | |
|---|---|---|---|
| Diet | F(1,32)=0.006 | p=0.937 | partial $\eta^2$=0.000 |
| Context | F(1,32)=0.063 | p=0.803 | partial $\eta^2$=0.002 |
| Time | F(2.28,73.02)=4.422 | **p=0.012** | partial $\eta^2$=**0.121** |
| Time × Diet | F(2.28,73.02)=2.370 | p=0.093 | partial $\eta^2$=0.069 |
| Time × Context | F(2.28,73.02)=0.784 | p=0.476 | partial $\eta^2$=0.024 |

| | |
|---|---|
| Diet × Context | $F(1,32)=1.073$ $p=0.308$ partial $\eta^2=0.032$ |
| Time × Diet × Context | $F(2.28,73.02)=0.383$ $p=0.710$ partial $\eta^2=0.012$ |

S18E Table

| **Females** | **Experiment 3 – Body Weight** |
|---|---|
| Diet | $F(1,30)=2.455$ $p=0.128$ partial $\eta^2=0.076$ |
| Context | $F(1,30)=3.497$ $p=0.071$ partial $\eta^2=0.104$ |
| Time | $F(4.57,137.0)=51.87$ $\mathbf{p<0.001}$ partial $\eta^2=\mathbf{0.634}$ |
| Time × Diet | $F(4.57,137.0)=1.496$ $p=0.200$ partial $\eta^2=0.047$ |
| Time × Context | $F(4.57,137.0)=0.288$ $p=0.906$ partial $\eta^2=0.010$ |
| Diet × Context | $F(1,30)=2.245$ $p=0.145$ partial $\eta^2=0.070$ |
| Time × Diet × Context | $F(4.57,137.0)=1.660$ $p=0.154$ partial $\eta^2=0.052$ |

S18F Table

| **Males** | **Experiment 3 – Body Weight** |
|---|---|
| Diet | $F(1,28)=0.120$ $p=0.732$ partial $\eta^2=0.004$ |
| Context | $F(1,28)=1.016$ $p=0.322$ partial $\eta^2=0.035$ |
| Time | $F(2.67,74.86)=12.22$ $p<0.001$ partial $\eta^2=0.304$ |
| Time × Diet | $F(2.67,74.86)=3.019$ $\mathbf{p=0.040}$ partial $\eta^2=\mathbf{0.097}$ |
| Time × Context | $F(2.67,74.86)=0.426$ $p=0.713$ partial $\eta^2=0.015$ |
| Diet × Context | $F(1,28)=0.150$ $p=0.701$ partial $\eta^2=0.005$ |
| Time × Diet × Context | $F(2.67,74.86)=0.422$ $p=0.715$ partial $\eta^2=0.015$ |

S12 Figure

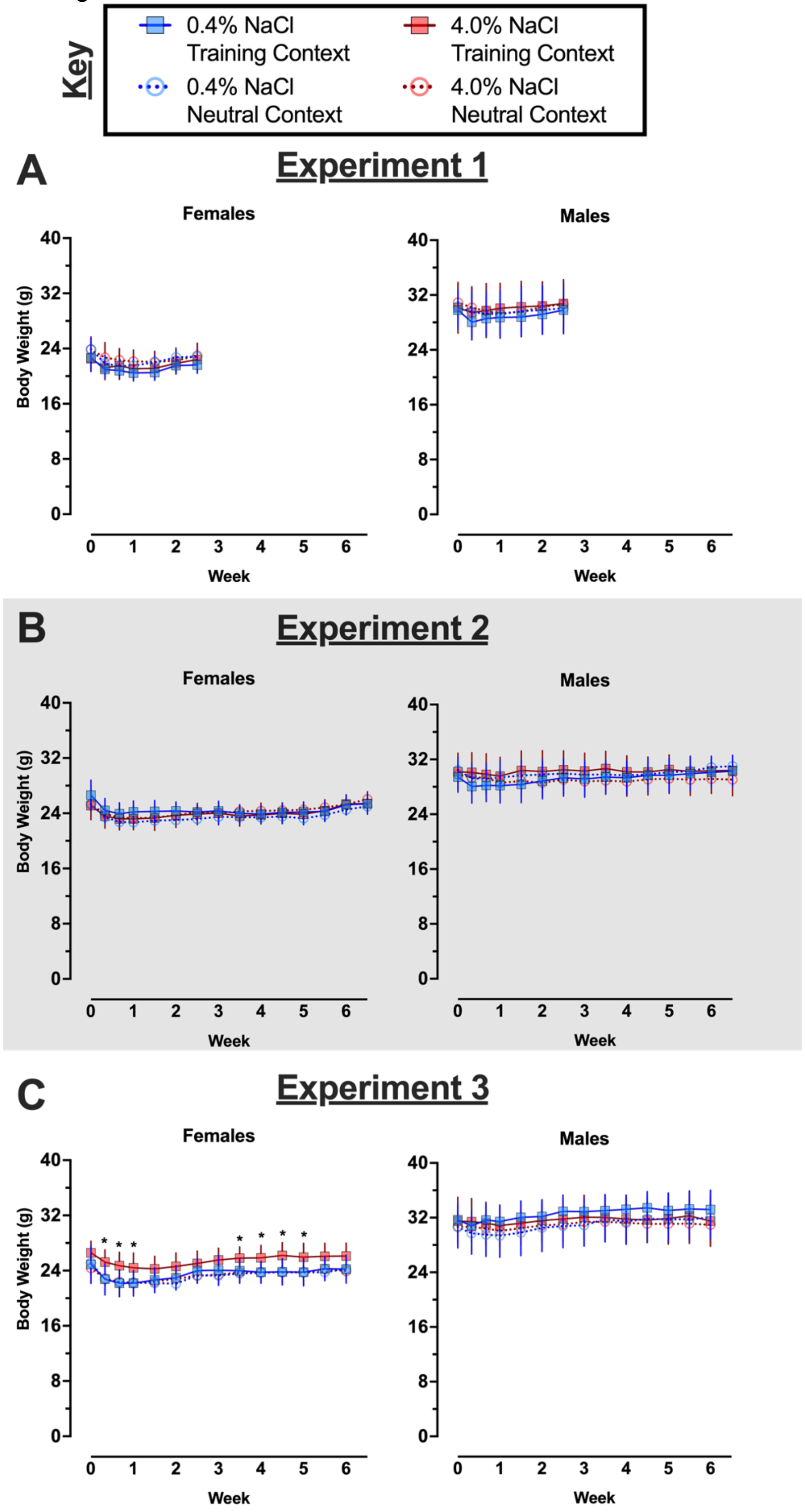
Key
0.4% NaCl Training Context
4.0% NaCl Training Context
0.4% NaCl Neutral Context
4.0% NaCl Neutral Context
A
Experiment 1
Females
Males
Body Weight (g)
Week
B
Experiment 2
Females
Males
Body Weight (g)
Week
C
Experiment 3
Females
Males
Body Weight (g)
Week

**S12 Fig. Body weight of context fear conditioned mice across Experiments.**

Mice assigned to 0.4% NaCl represented by blue symbols, mice assigned to 4.0% NaCl represented by red symbols; mice to be tested in Training Context represented by squares and solid lines, mice to be tested in Neutral Context represented by circles and dotted lines. Body weights were measured twice weekly of mice in A) Experiment 1, B) Experiment 2 (grey shading), and C) Experiment 3. Some data loss occurred on the very last weighing day for a subset of animals in C) Experiment 3, thus graphs and repeated measures statistical analyses for Experiment 3 body weights cease at week 6 to maximize inclusion of mice in repeated measures analyses. Experiment 1: 0.4% NaCl females Training Context, n=8; 0.4% NaCl females Neutral Context, n=9; 4.0% NaCl females Training Context, n=8; 4.0% NaCl females Neutral Context, n=9; 0.4% NaCl males Training Context, n=8; 0.4% NaCl males Neutral Context, n=9; 4.0% NaCl males Training Context, n=8; 4.0% NaCl males Neutral Context, n=8. Experiment 2: 0.4% NaCl females Training Context, n=9; 0.4% NaCl females Neutral Context, n=9; 4.0% NaCl females Training Context, n=8; 4.0% NaCl females Neutral Context, n=8; 0.4% NaCl males Training Context, n=9; 0.4% NaCl males Neutral Context, n=9; 4.0% NaCl males Training Context, n=8; 4.0% NaCl males Neutral Context, n=10. Experiment 3: 0.4% NaCl females Training Context, n=8; 0.4% NaCl females Neutral Context, n=9; 4.0% NaCl females Training Context, n=9; 4.0% NaCl females Neutral Context, n=8; 0.4% NaCl males Training Context, n=8; 0.4% NaCl males Neutral Context, n=8; 4.0% NaCl males Training Context, n=8; 4.0% NaCl males Neutral Context, n=8. Data are graphed as mean ± 95% confidence interval. *$p<0.05$ indicates difference between mice within the same sex consuming 0.4% NaCl versus 4.0% NaCl and tested in Training Context.

**S19 Table. Three-way repeated measures ANOVAs on weekly average water consumption per day for control no shock mice across Experiments.**

S19A Table

| Experiment 1 | Water/day | | |
|---|---|---|---|
| Sex | $F(1,31)=3.245$ | $p=0.081$ | partial $\eta^2=0.095$ |
| Diet | $F(1,31)=50.99$ | **$p<0.001$** | partial $\eta^2=$**0.622** |
| Time | $F(1.67,51.69)=3.219$ | $p=0.057$ | partial $\eta^2=0.094$ |
| Time × Sex | $F(1.67,51.69)=4.157$ | **$p=0.027$** | partial $\eta^2=$**0.118** |
| Time × Diet | $F(1.67,51.69)=1.459$ | $p=0.242$ | partial $\eta^2=0.045$ |
| Sex × Diet | $F(1,31)=3.868$ | $p=0.058$ | partial $\eta^2=0.111$ |
| Time × Sex × Diet | $F(1.67,51.69)=0.910$ | $p=0.393$ | partial $\eta^2=0.029$ |

S19B Table

| Experiment 2 | Water/day | | |
|---|---|---|---|
| Sex | $F(1,29)=9.248$ | $p=0.005$ | partial $\eta^2=0.242$ |
| Diet | $F(1,29)=204.6$ | $p<0.001$ | partial $\eta^2=0.876$ |
| Time | $F(3.98,115.4)=17.93$ | $p<0.001$ | partial $\eta^2=0.382$ |
| Time × Sex | $F(3.98,115.4)=6.510$ | **$p<0.001$** | partial $\eta^2=$**0.183** |
| Time × Diet | $F(3.98,115.4)=5.307$ | **$p<0.001$** | partial $\eta^2=$**0.155** |
| Sex × Diet | $F(1,29)=0.231$ | $p=0.635$ | partial $\eta^2=0.008$ |
| Time × Sex × Diet | $F(3.98,115.4)=0.084$ | $p=0.987$ | partial $\eta^2=0.003$ |

S19C Table

| Experiment 3 | Water/day | | |
|---|---|---|---|
| Sex | $F(1,28)=0.188$ | $p=0.668$ | partial $\eta^2=0.007$ |
| Diet | $F(1,28)=37.59$ | $p<0.001$ | partial $\eta^2=0.573$ |
| Time | $F(3.47,97.21)=5.224$ | $p=0.001$ | partial $\eta^2=0.157$ |
| Time × Sex | $F(3.47,97.21)=0.962$ | $p=0.423$ | partial $\eta^2=0.033$ |
| Time × Diet | $F(3.47,97.21)=3.541$ | $p=0.013$ | partial $\eta^2=0.112$ |
| Sex × Diet | $F(1,28)=3.551$ | $p=0.070$ | partial $\eta^2=0.113$ |
| Time × Sex × Diet | $F(3.47,97.21)=3.387$ | **$p=0.016$** | partial $\eta^2=$**0.108** |

**S20 Table. Three-way repeated measures ANOVAs on weekly average food consumption per day for control no shock mice across Experiments.**

S20A Table

| **Experiment 1** | **Food/day** | | |
|---|---|---|---|
| Sex | $F(1,31)=17.58$ | **$p<0.001$** | partial $\eta^2$=**0.362** |
| Diet | $F(1,31)=8.329$ | **$p=0.007$** | partial $\eta^2$=**0.212** |
| Time | $F(1.56,48.19)=0.003$ | $p=0.990$ | partial $\eta^2=0.000$ |
| Time × Sex | $F(1.56,48.19)=2.401$ | $p=0.113$ | partial $\eta^2=0.072$ |
| Time × Diet | $F(1.56,48.19)=0.223$ | $p=0.745$ | partial $\eta^2=0.007$ |
| Sex × Diet | $F(1,31)=0.130$ | $p=0.721$ | partial $\eta^2=0.004$ |
| Time × Sex × Diet | $F(1.56,48.19)=1.258$ | $p=0.286$ | partial $\eta^2=0.039$ |

S20B Table

| **Experiment 2** | **Food/day** | | |
|---|---|---|---|
| Sex | $F(1,29)=22.47$ | **$p<0.001$** | partial $\eta^2$=**0.437** |
| Diet | $F(1,29)=15.77$ | $p<0.001$ | partial $\eta^2=0.352$ |
| Time | $F(3.08,89.34)=2.245$ | $p=0.087$ | partial $\eta^2=0.072$ |
| Time × Sex | $F(3.08,89.34)=0.476$ | $p=0.705$ | partial $\eta^2=0.016$ |
| Time × Diet | $F(3.08,89.34)=3.463$ | **$p=0.019$** | partial $\eta^2$=**0.107** |
| Sex × Diet | $F(1,29)=0.969$ | $p=0.333$ | partial $\eta^2=0.032$ |
| Time × Sex × Diet | $F(3.08,89.34)=0.617$ | $p=0.610$ | partial $\eta^2=0.021$ |

S20C Table

| **Experiment 3** | **Food/day** | | |
|---|---|---|---|
| Sex | $F(1,28)=24.30$ | **$p<0.001$** | partial $\eta^2$=**0.465** |
| Diet | $F(1,28)=7.983$ | **$p=0.009$** | partial $\eta^2$=**0.222** |
| Time | $F(2.76,77.39)=1.692$ | $p=0.179$ | partial $\eta^2=0.057$ |
| Time × Sex | $F(2.76,77.39)=1.523$ | $p=0.218$ | partial $\eta^2=0.052$ |
| Time × Diet | $F(2.76,77.39)=1.496$ | $p=0.224$ | partial $\eta^2=0.051$ |
| Sex × Diet | $F(1,28)=0.735$ | $p=0.398$ | partial $\eta^2=0.026$ |
| Time × Sex × Diet | $F(2.76,77.39)=1.174$ | $p=0.324$ | partial $\eta^2=0.040$ |

S13 Figure

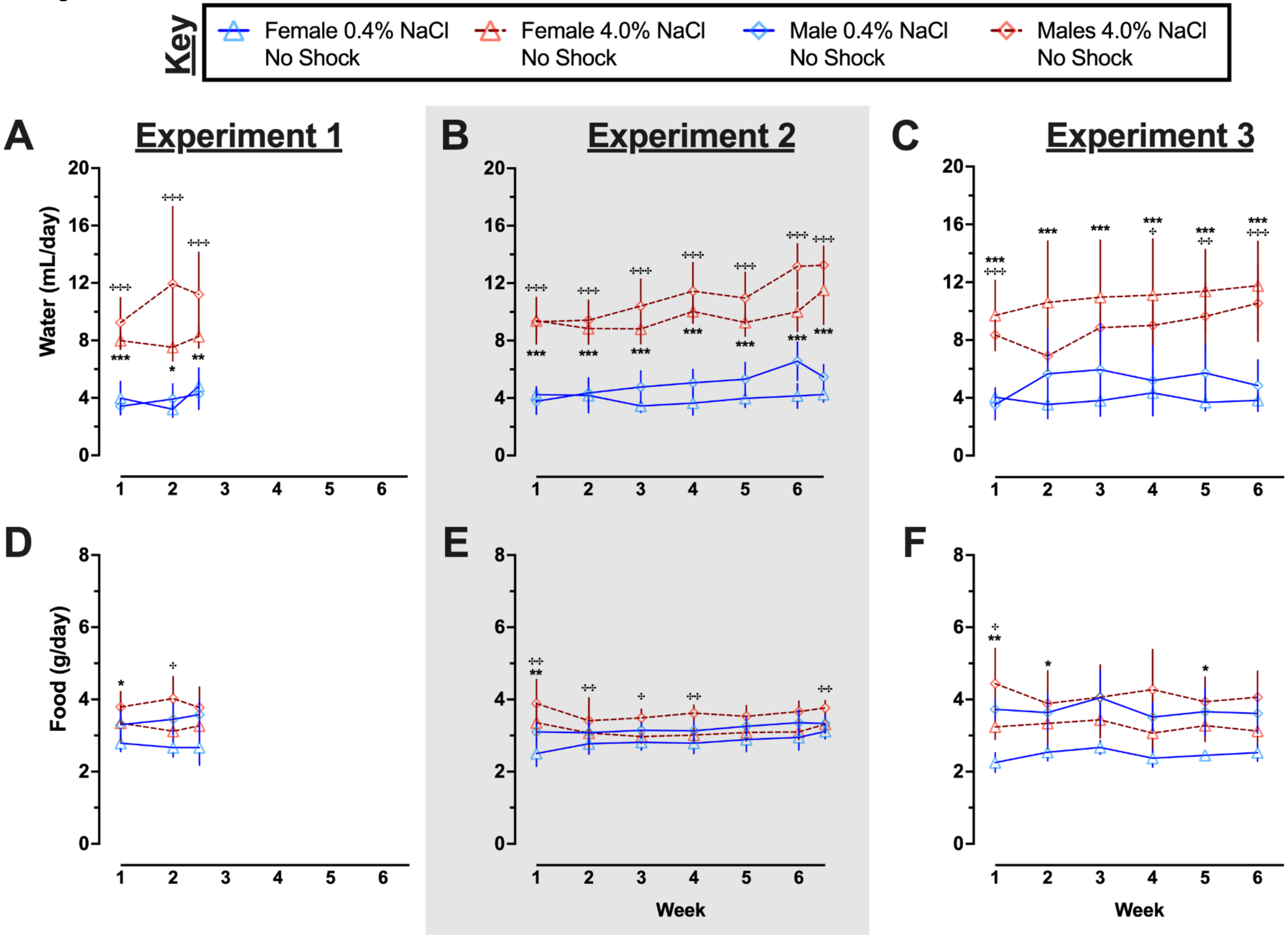
Key
Female 0.4% NaCl No Shock
Female 4.0% NaCl No Shock
Male 0.4% NaCl No Shock
Males 4.0% NaCl No Shock
A
Experiment 1
B
Experiment 2
C
Experiment 3
Water (mL/day)
D
E
F
Food (g/day)
Week
Week

**S13 Fig. Average water and food consumed per day by control no shock mice across Experiments.**

Females represented by triangles, males by diamonds; 0.4% NaCl represented by blue symbols and solid lines, 4.0% NaCl represented by red symbols and dashed lines. A, B, C) Water and D, E, F) food consumption was measured twice weekly, and full weeks were averaged; partial weeks at the conclusion of A, D) Experiment 1 and B, E) Experiment 2 (grey shading) are included in the graphs. Some data loss occurred on the very last weighing day for a subset of animals in C, F) Experiment 3, thus graphs and repeated measures statistical analyses for Experiment 3 consumption cease at week 6 to maximize inclusion of mice in repeated measures analyses. Experiment 1: 0.4% NaCl females, n=9; 4.0% NaCl females, n=9; 0.4% NaCl males, n=8; 4.0% NaCl males, n=9. Experiment 2: 0.4% NaCl females, n=8; 4.0% NaCl females, n=7; 0.4% NaCl males, n=9; 4.0% NaCl males, n=9. Experiment 3: 0.4% NaCl females, n=8; 4.0% NaCl females, n=8; 0.4% NaCl males, n=8; 4.0% NaCl males, n=8. Data are graphed as mean ± 95% confidence interval. *$p<0.05$, **$p<0.01$, ***$p<0.001$ indicate difference between females consuming 0.4% NaCl versus 4.0% NaCl. ‡$p<0.05$, ‡‡$p<0.01$, ‡‡‡$p<0.001$ indicate difference between males consuming 0.4% NaCl versus 4.0% NaCl.

**S21 Table. Three-way repeated measures ANOVAs on weekly body weight changes for control no shock mice across Experiments.**

S21A Table

| **Experiment 1** | **Body Weight Change** | | |
|---|---|---|---|
| Sex | F(1,31)=0.236 | p=0.631 | partial $\eta^2$=0.008 |
| Diet | F(1,31)=0.443 | p=0.511 | partial $\eta^2$=0.014 |
| Time | F(1.84,42.05)=59.12 | **p<0.001** | partial $\eta^2$=**0.656** |
| Time × Sex | F(1.84,42.05)=0.032 | p=0.960 | partial $\eta^2$=0.001 |
| Time × Diet | F(1.84,42.05)=1.319 | p=0.274 | partial $\eta^2$=0.041 |
| Sex × Diet | F(1,31)=6.755 | **p=0.014** | partial $\eta^2$=**0.179** |
| Time × Sex × Diet | F(1.84,42.05)=0.491 | p=0.599 | partial $\eta^2$=0.016 |

S21B Table

| **Experiment 2** | **Body Weight Change** | | |
|---|---|---|---|
| Sex | F(1,29)=0.966 | p=0.334 | partial $\eta^2$=0.032 |
| Diet | F(1,29)=1.596 | p=0.216 | partial $\eta^2$=0.052 |
| Time | F(4.00,115.9)=29.74 | p<0.001 | partial $\eta^2$=0.506 |
| Time × Sex | F(4.00,115.9)=4.637 | **p=0.002** | partial $\eta^2$=**0.138** |
| Time × Diet | F(4.00,115.9)=3.270 | **p=0.014** | partial $\eta^2$=**0.101** |
| Sex × Diet | F(1,29)=0.562 | p=0.460 | partial $\eta^2$=0.019 |
| Time × Sex × Diet | F(4.00,115.9)=0.530 | p=0.713 | partial $\eta^2$=0.018 |

S21C Table

| **Experiment 3** | **Body Weight Change** | | |
|---|---|---|---|
| Sex | F(1,28)=6.100 | p=0.020 | partial $\eta^2$=0.179 |
| Diet | F(1,28)=3.939 | p=0.057 | partial $\eta^2$=0.123 |
| Time | F(3.14,87.80)=18.35 | p<0.001 | partial $\eta^2$=0.396 |
| Time × Sex | F(3.14,87.80)=4.494 | **p=0.005** | partial $\eta^2$=**0.138** |
| Time × Diet | F(3.14,87.80)=4.231 | **p=0.007** | partial $\eta^2$=**0.131** |
| Sex × Diet | F(1,28)=0.136 | p=0.715 | partial $\eta^2$=0.005 |
| Time × Sex × Diet | F(3.14,87.80)=1.041 | p=0.380 | partial $\eta^2$=0.036 |

**S22 Table. Three-way repeated measures ANOVAs on twice weekly body weights of control no shock mice across Experiments.**

S22A Table

| Experiment 1 | Body Weight | | |
|---|---|---|---|
| Sex | $F(1,31)=25.84$ | $p<0.001$ | partial $\eta^2=0.455$ |
| Diet | $F(1,31)=0.018$ | $p=0.893$ | partial $\eta^2=0.001$ |
| Time | $F(2.31,71.60)=21.32$ | $p<0.001$ | partial $\eta^2=0.408$ |
| Time × Sex | $F(2.31,71.60)=0.174$ | $p=0.869$ | partial $\eta^2=0.006$ |
| Time × Diet | $F(2.31,71.60)=0.667$ | $p=0.537$ | partial $\eta^2=0.021$ |
| Sex × Diet | $F(1,31)=0.155$ | $p=0.697$ | partial $\eta^2=0.179$ |
| Time × Sex × Diet | $F(2.31,71.60)=3.707$ | **$p=0.024$** | partial $\eta^2=$**0.107** |

S22B Table

| Experiment 2 | Body Weight | | |
|---|---|---|---|
| Sex | $F(1,29)=38.62$ | $p<0.001$ | partial $\eta^2=0.571$ |
| Diet | $F(1,29)=2.447$ | $p=0.129$ | partial $\eta^2=0.078$ |
| Time | $F(3.30,95.56)=13.28$ | $p<0.001$ | partial $\eta^2=0.314$ |
| Time × Sex | $F(3.30,95.56)=4.031$ | **$p=0.008$** | partial $\eta^2=$**0.122** |
| Time × Diet | $F(3.30,95.56)=2.077$ | $p=0.102$ | partial $\eta^2=0.067$ |
| Sex × Diet | $F(1,29)=0.418$ | $p=0.523$ | partial $\eta^2=0.014$ |
| Time × Sex × Diet | $F(3.30,95.56)=0.961$ | $p=0.421$ | partial $\eta^2=0.032$ |

S22C Table

| Experiment 3 | Body Weight | | |
|---|---|---|---|
| Sex | $F(1,28)=59.03$ | $p<0.001$ | partial $\eta^2=0.678$ |
| Diet | $F(1,28)=1.095$ | $p=0.304$ | partial $\eta^2=0.038$ |
| Time | $F(2.64,73.82)=11.92$ | $p<0.001$ | partial $\eta^2=0.299$ |
| Time × Sex | $F(2.64,73.82)=3.879$ | **$p=0.016$** | partial $\eta^2=$**0.122** |
| Time × Diet | $F(2.64,73.82)=2.216$ | $p=0.101$ | partial $\eta^2=0.073$ |
| Sex × Diet | $F(1,28)=0.039$ | $p=0.845$ | partial $\eta^2=0.001$ |
| Time × Sex × Diet | $F(2.64,73.82)=0.564$ | $p=0.618$ | partial $\eta^2=0.020$ |

S14 Figure

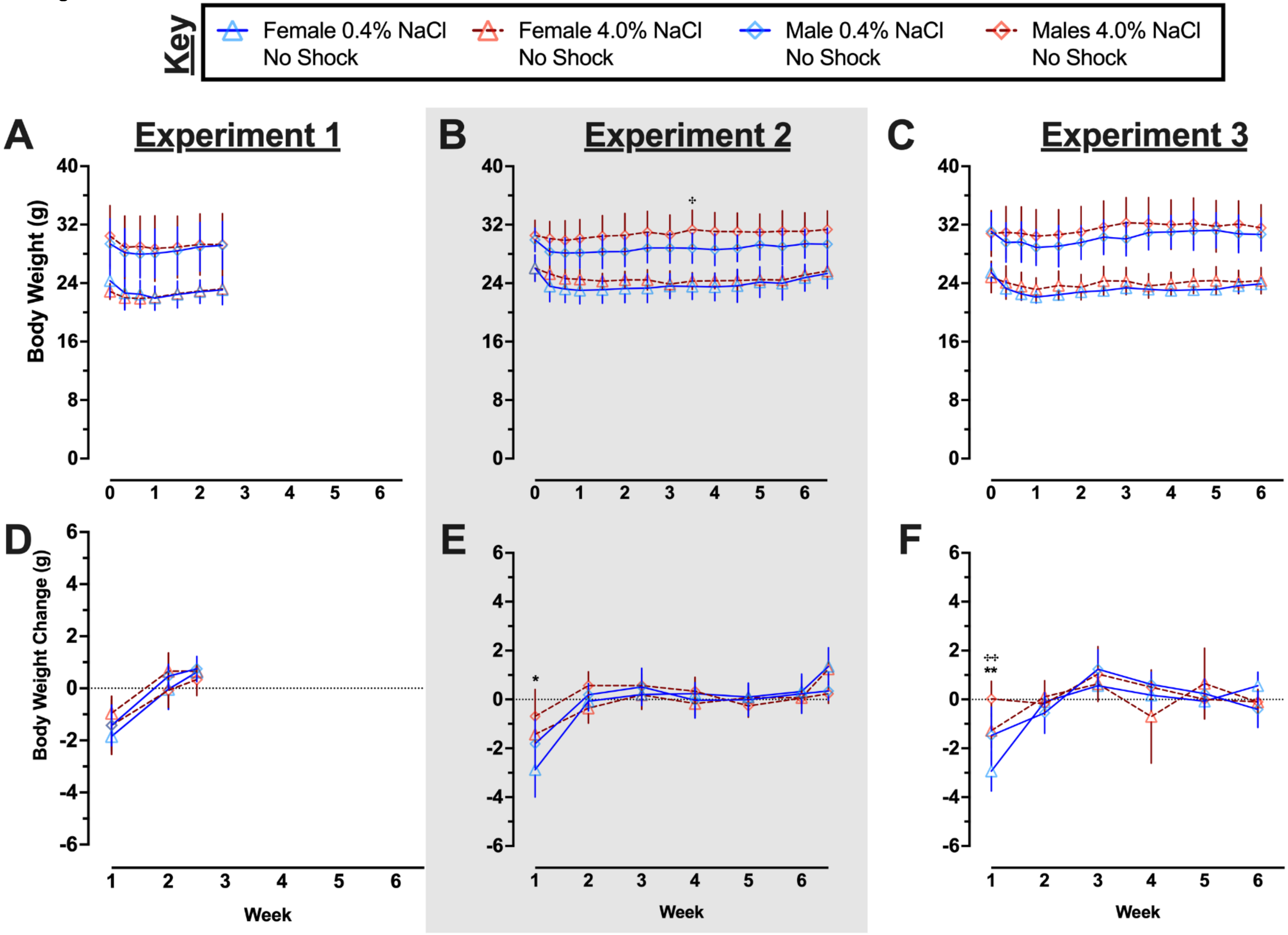

Key
Female 0.4% NaCl No Shock
Female 4.0% NaCl No Shock
Male 0.4% NaCl No Shock
Males 4.0% NaCl No Shock
A
Experiment 1
B
Experiment 2
C
Experiment 3
Body Weight (g)
40
32
24
16
8
0
0 1 2 3 4 5 6
D
E
F
Body Weight Change (g)
6
4
2
0
-2
-4
-6
1 2 3 4 5 6
Week
*
**

**S14 Fig. Body weights and body weight changes in control no shock mice across Experiments.**

Females represented by triangles, males by diamonds; 0.4% NaCl represented by blue symbols and solid lines, 4.0% NaCl represented by red symbols and dashed lines. A, B, C) Body weights were measured twice weekly, and D, E, F) body weight changes were calculated across full weeks. Partial weeks at the conclusion of A, D) Experiment 1 and B, E) Experiment 2 (grey shading) are included in the graphs. Some data loss occurred on the very last weighing day for a subset of animals in C, F) Experiment 3, thus graphs and repeated measures statistical analyses for Experiment 3 consumption cease at week 6 to maximize inclusion of mice in repeated measures analyses. Experiment 1: 0.4% NaCl females, n=9; 4.0% NaCl females, n=9; 0.4% NaCl males, n=8; 4.0% NaCl males, n=9. Experiment 2: 0.4% NaCl females, n=8; 4.0% NaCl females, n=7; 0.4% NaCl males, n=9; 4.0% NaCl males, n=9. Experiment 3: 0.4% NaCl females, n=8; 4.0% NaCl females, n=8; 0.4% NaCl males, n=8; 4.0% NaCl males, n=8. Data are graphed as mean ± 95% confidence interval. *p<0.05, **p<0.01, indicate difference between females consuming 0.4% NaCl versus 4.0% NaCl. $^{+}$p<0.05, $^{++}$p<0.01, indicate difference between males consuming 0.4% NaCl versus 4.0% NaCl.

**S23 Table. Three-way repeated measures ANOVAs on weekly average NaCl consumed per day by context fear conditioned mice across Experiments.**

S23A Table

| Females | Experiment 1 – NaCl/day | | |
|---|---|---|---|
| Diet | $F(1,30)=635.7$ | **$p<0.001$** | partial $\eta^2$=**0.955** |
| Context | $F(1,30)=0.004$ | $p=0.951$ | partial $\eta^2=0.000$ |
| Time | $F(1.61,48.36)=0.846$ | $p=0.413$ | partial $\eta^2=0.027$ |
| Time × Diet | $F(1.61,48.36)=0.834$ | $p=0.418$ | partial $\eta^2=0.027$ |
| Time × Context | $F(1.61,48.36)=0.698$ | $p=0.473$ | partial $\eta^2=0.023$ |
| Diet × Context | $F(1,30)=0.029$ | $p=0.866$ | partial $\eta^2=0.001$ |
| Time × Diet × Context | $F(1.61,48.36)=0.627$ | $p=0.505$ | partial $\eta^2=0.020$ |

S23B Table

| Males | Experiment 1 – NaCl/day | | |
|---|---|---|---|
| Diet | $F(1,29)=784.5$ | **$p<0.001$** | partial $\eta^2$=**0.964** |
| Context | $F(1,29)=0.017$ | $p=0.899$ | partial $\eta^2=0.001$ |
| Time | $F(1.64,47.50)=0.759$ | $p=0.449$ | partial $\eta^2=0.025$ |
| Time × Diet | $F(1.64,47.50)=0.377$ | $p=0.646$ | partial $\eta^2=0.013$ |
| Time × Context | $F(1.64,47.50)=0.312$ | $p=0.690$ | partial $\eta^2=0.011$ |
| Diet × Context | $F(1,29)=0.014$ | $p=0.906$ | partial $\eta^2=0.000$ |
| Time × Diet × Context | $F(1.64,47.50)=0.497$ | $p=0.574$ | partial $\eta^2=0.017$ |

S23C Table

| Females | Experiment 2 – NaCl/day | | |
|---|---|---|---|
| Diet | $F(1,30)=1412$ | $p<0.001$ | partial $\eta^2=0.979$ |
| Context | $F(1,30)=0.000$ | $p=0.988$ | partial $\eta^2=0.000$ |
| Time | $F(3.22,96.53)=6.807$ | $p<0.001$ | partial $\eta^2=0.185$ |
| Time × Diet | $F(3.22,96.53)=4.373$ | **$p=0.005$** | partial $\eta^2$=**0.127** |
| Time × Context | $F(3.22,96.53)=0.929$ | $p=0.435$ | partial $\eta^2=0.030$ |
| Diet × Context | $F(1,30)=0.002$ | $p=0.962$ | partial $\eta^2=0.000$ |
| Time × Diet × Context | $F(3.22,96.53)=0.760$ | $p=0.528$ | partial $\eta^2=0.025$ |

S23D Table

| Males | Experiment 2 – NaCl/day | | |
|---|---|---|---|
| Diet | $F(1,32)=2164$ | $p<0.001$ | partial $\eta^2=0.985$ |
| Context | $F(1,32)=0.904$ | $p=0.349$ | partial $\eta^2=0.027$ |
| Time | $F(4.11,131.4)=6.013$ | $p<0.001$ | partial $\eta^2=0.158$ |
| Time × Diet | $F(4.11,131.4)=4.350$ | **$p=0.002$** | partial $\eta^2$=**0.120** |
| Time × Context | $F(4.11,131.4)=0.943$ | $p=0.443$ | partial $\eta^2=0.029$ |

| | | | |
|---|---|---|---|
| Diet × Context | $F(1,32)=1.057$ | $p=0.312$ | partial $\eta^2=0.032$ |
| Time × Diet × Context | $F(4.11,131.4)=1.003$ | $p=0.410$ | partial $\eta^2=0.030$ |

S23E Table

| **Females** | **Experiment 3 – NaCl/day** | | |
|---|---|---|---|
| Diet | $F(1,30)=621.9$ | **$p<0.001$** | partial $\eta^2=$**0.954** |
| Context | $F(1,30)=0.301$ | $p=0.587$ | partial $\eta^2=0.010$ |
| Time | $F(2.27,68.08)=1.194$ | $p=0.313$ | partial $\eta^2=0.038$ |
| Time × Diet | $F(2.27,68.08)=0.710$ | $p=0.512$ | partial $\eta^2=0.023$ |
| Time × Context | $F(2.27,68.08)=0.192$ | $p=0.852$ | partial $\eta^2=0.006$ |
| Diet × Context | $F(1,30)=0.177$ | $p=0.677$ | partial $\eta^2=0.006$ |
| Time × Diet × Context | $F(2.27,68.08)=0.190$ | $p=0.853$ | partial $\eta^2=0.006$ |

S23F Table

| **Males** | **Experiment 3 – NaCl/day** | | |
|---|---|---|---|
| Diet | $F(1,28)=398.8$ | **$p<0.001$** | partial $\eta^2=$**0.934** |
| Context | $F(1,28)=0.186$ | $p=0.670$ | partial $\eta^2=0.007$ |
| Time | $F(2.94,82.41)=2.553$ | $p=0.062$ | partial $\eta^2=0.084$ |
| Time × Diet | $F(2.94,82.41)=1.989$ | $p=0.123$ | partial $\eta^2=0.066$ |
| Time × Context | $F(2.94,82.41)=0.723$ | $p=0.538$ | partial $\eta^2=0.025$ |
| Diet × Context | $F(1,28)=0.244$ | $p=0.625$ | partial $\eta^2=0.009$ |
| Time × Diet × Context | $F(2.94,82.41)=0.626$ | $p=0.597$ | partial $\eta^2=0.022$ |

S15 Figure

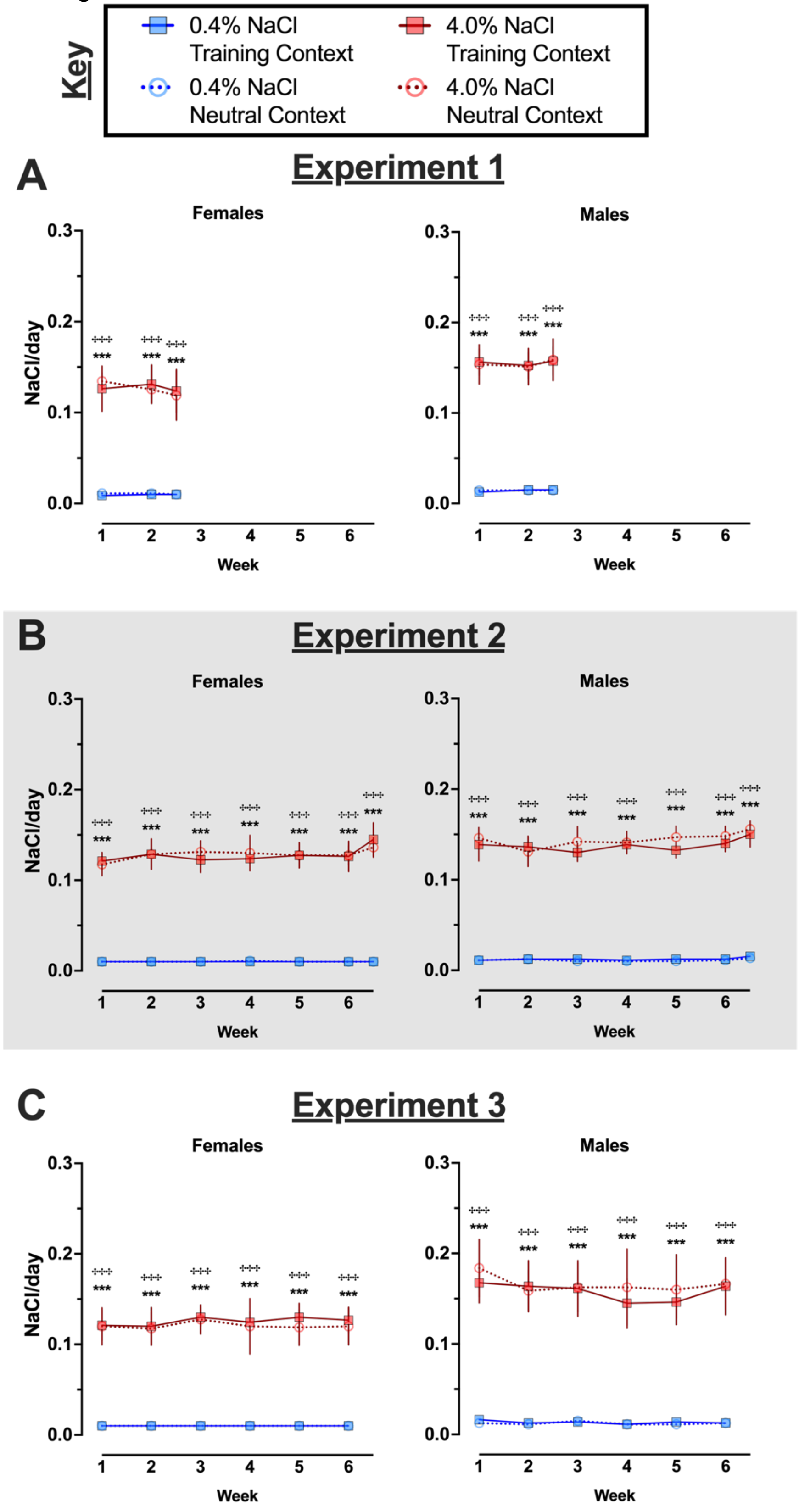

Key
0.4% NaCl Training Context
4.0% NaCl Training Context
0.4% NaCl Neutral Context
4.0% NaCl Neutral Context
A
Experiment 1
Females
Males
NaCl/day
Week
B
Experiment 2
Females
Males
NaCl/day
Week
C
Experiment 3
Females
Males
NaCl/day
Week

**S15 Fig. Average NaCl consumed per day by context fear conditioned mice across Experiments.**

Mice assigned to 0.4% NaCl represented by blue symbols, mice assigned to 4.0% NaCl represented by red symbols; mice to be tested in Training Context represented by squares and solid lines, mice to be tested in Neutral Context represented by circles and dotted lines. NaCl consumption was calculated based upon the amount of each diet consumed per day and the diet's respective NaCl percentage (0.4 or 4.0%, w/w). These calculations were made for each full week, plus the partial weeks at the conclusion of A) Experiment 1 and B) Experiment 2 (grey shading). Some data loss occurred on the very last weighing day for a subset of animals in C) Experiment 3, thus graphs and repeated measures statistical analyses for Experiment 3 calculations cease at week 6 to maximize inclusion of mice in repeated measures analyses. Experiment 1: 0.4% NaCl females Training Context, n=8; 0.4% NaCl females Neutral Context, n=9; 4.0% NaCl females Training Context, n=8; 4.0% NaCl females Neutral Context, n=9; 0.4% NaCl males Training Context, n=8; 0.4% NaCl males Neutral Context, n=9; 4.0% NaCl males Training Context, n=8; 4.0% NaCl males Neutral Context, n=8. Experiment 2: 0.4% NaCl females Training Context, n=9; 0.4% NaCl females Neutral Context, n=9; 4.0% NaCl females Training Context, n=8; 4.0% NaCl females Neutral Context, n=8; 0.4% NaCl males Training Context, n=9; 0.4% NaCl males Neutral Context, n=9; 4.0% NaCl males Training Context, n=8; 4.0% NaCl males Neutral Context, n=10. Experiment 3: 0.4% NaCl females Training Context, n=8; 0.4% NaCl females Neutral Context, n=9; 4.0% NaCl females Training Context, n=9; 4.0% NaCl females Neutral Context, n=8; 0.4% NaCl males Training Context, n=8; 0.4% NaCl males Neutral Context, n=8; 4.0% NaCl males Training Context, n=8; 4.0% NaCl males Neutral Context, n=8. Data are graphed as mean ± 95% confidence interval. *p<0.05, **p<0.01, ***p<0.001 indicate difference between mice within the same sex consuming 0.4% NaCl versus 4.0% NaCl and tested in Training Context. $^{+}p<0.05$, $^{++}p<0.01$, $^{+++}p<0.001$ indicate difference

between mice within the same sex consuming 0.4% NaCl versus 4.0% NaCl and tested in Neutral Context.

**S24 Table. Three-way repeated measures ANOVAs on weekly average NaCl consumed as a percentage of body weight in context fear conditioned mice across Experiments.**

S24A Table

| **Females** | **Experiment 1 – NaCl as % BW** | | |
|---|---|---|---|
| Diet | $F(1,30)=532.7$ | **$p<0.001$** | partial $\eta^2$=**0.947** |
| Context | $F(1,30)=0.408$ | $p=0.528$ | partial $\eta^2=0.013$ |
| Time | $F(1.51,45.38)=2.849$ | $p=0.082$ | partial $\eta^2=0.087$ |
| Time × Diet | $F(1.51,45.38)=2.372$ | $p=0.117$ | partial $\eta^2=0.073$ |
| Time × Context | $F(1.51,45.38)=0.648$ | $p=0.487$ | partial $\eta^2=0.021$ |
| Diet × Context | $F(1,30)=0.378$ | $p=0.544$ | partial $\eta^2=0.012$ |
| Time × Diet × Context | $F(1.51,45.38)=0.578$ | $p=0.519$ | partial $\eta^2=0.019$ |

S24B Table

| **Males** | **Experiment 1 – NaCl as % BW** | | |
|---|---|---|---|
| Diet | $F(1,29)=496.0$ | **$p<0.001$** | partial $\eta^2$=**0.945** |
| Context | $F(1,29)=0.002$ | $p=0.968$ | partial $\eta^2=0.000$ |
| Time | $F(1.75,50.83)=0.847$ | $p=0.421$ | partial $\eta^2=0.028$ |
| Time × Diet | $F(1.75,50.83)=0.841$ | $p=0.424$ | partial $\eta^2=0.028$ |
| Time × Context | $F(1.75,50.83)=0.937$ | $p=0.388$ | partial $\eta^2=0.031$ |
| Diet × Context | $F(1,29)=0.005$ | $p=0.944$ | partial $\eta^2=0.000$ |
| Time × Diet × Context | $F(1.75,50.83)=1.047$ | $p=0.351$ | partial $\eta^2=0.035$ |

S24C Table

| **Females** | **Experiment 2 – NaCl as % BW** | | |
|---|---|---|---|
| Diet | $F(1,30)=1804$ | **$p<0.001$** | partial $\eta^2$=**0.984** |
| Context | $F(1,30)=0.102$ | $p=0.752$ | partial $\eta^2=0.003$ |
| Time | $F(3.02,90.73)=2.635$ | $p=0.054$ | partial $\eta^2=0.081$ |
| Time × Diet | $F(3.02,90.73)=1.623$ | $p=0.189$ | partial $\eta^2=0.051$ |
| Time × Context | $F(3.02,90.73)=1.068$ | $p=0.367$ | partial $\eta^2=0.034$ |
| Diet × Context | $F(1,30)=0.162$ | $p=0.690$ | partial $\eta^2=0.005$ |
| Time × Diet × Context | $F(3.02,90.73)=0.897$ | $p=0.447$ | partial $\eta^2=0.029$ |

S24D Table

| **Males** | **Experiment 2 – NaCl as % BW** | | |
|---|---|---|---|
| Diet | $F(1,32)=819.4$ | $p<0.001$ | partial $\eta^2=0.962$ |
| Context | $F(1,32)=1.631$ | $p=0.211$ | partial $\eta^2=0.048$ |
| Time | $F(3.82,122.1)=4.817$ | $p=0.001$ | partial $\eta^2=0.131$ |
| Time × Diet | $F(3.82,122.1)=3.901$ | **$p=0.006$** | partial $\eta^2$=**0.109** |
| Time × Context | $F(3.82,122.1)=0.665$ | $p=0.610$ | partial $\eta^2=0.020$ |

| | | | |
|---|---|---|---|
| Diet × Context | $F(1,32)=1.895$ | $p=0.178$ | partial $\eta^2=0.056$ |
| Time × Diet × Context | $F(3.82,122.1)=0.665$ | $p=0.610$ | partial $\eta^2=0.020$ |

S24E Table

| **Females** | **Experiment 3 – NaCl as % BW** | | |
|---|---|---|---|
| Diet | $F(1,30)=873.8$ | **$p<0.001$** | partial $\eta^2=$**0.967** |
| Context | $F(1,30)=0.439$ | $p=0.513$ | partial $\eta^2=0.014$ |
| Time | $F(1.92,57.61)=1.531$ | $p=0.225$ | partial $\eta^2=0.049$ |
| Time × Diet | $F(1.92,57.61)=0.948$ | $p=0.390$ | partial $\eta^2=0.031$ |
| Time × Context | $F(1.92,57.61)=0.237$ | $p=0.781$ | partial $\eta^2=0.008$ |
| Diet × Context | $F(1,30)=0.621$ | $p=0.437$ | partial $\eta^2=0.020$ |
| Time × Diet × Context | $F(1.92,57.61)=0.197$ | $p=0.813$ | partial $\eta^2=0.007$ |

S24F Table

| **Males** | **Experiment 3 – NaCl as % BW** | | |
|---|---|---|---|
| Diet | $F(1,28)=772.2$ | **$p<0.001$** | partial $\eta^2=$**0.965** |
| Context | $F(1,28)=0.941$ | $p=0.340$ | partial $\eta^2=0.033$ |
| Time | $F(2.75,76.95)=3.738$ | **$p=0.017$** | partial $\eta^2=$**0.118** |
| Time × Diet | $F(2.75,76.95)=2.772$ | $p=0.052$ | partial $\eta^2=0.090$ |
| Time × Context | $F(2.75,76.95)=0.699$ | $p=0.544$ | partial $\eta^2=0.024$ |
| Diet × Context | $F(1,28)=0.807$ | $p=0.377$ | partial $\eta^2=0.028$ |
| Time × Diet × Context | $F(2.75,76.95)=0.627$ | $p=0.586$ | partial $\eta^2=0.022$ |

S16 Figure

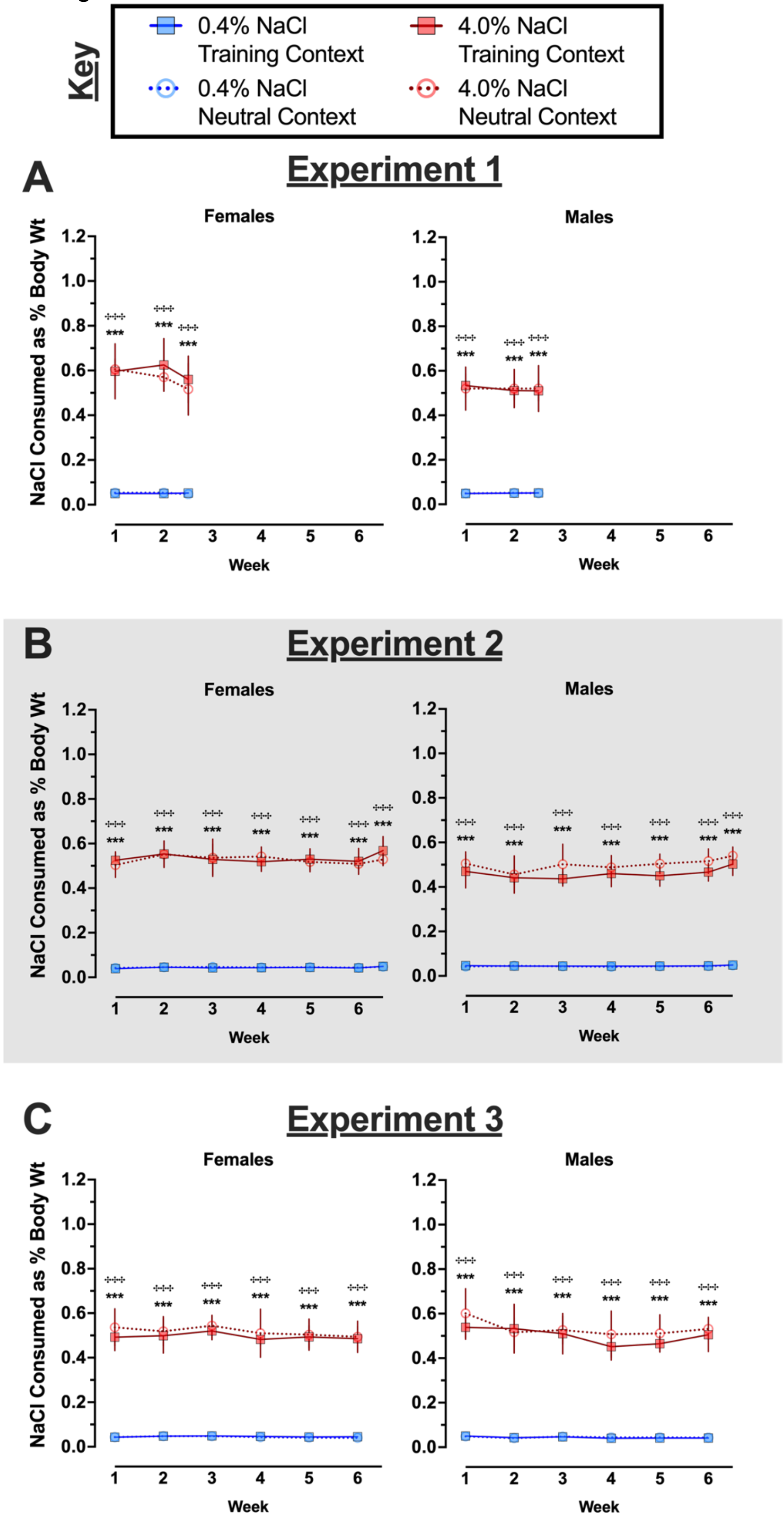

Key
0.4% NaCl Training Context
4.0% NaCl Training Context
0.4% NaCl Neutral Context
4.0% NaCl Neutral Context
A
Experiment 1
Females
Males
NaCl Consumed as % Body Wt
Week
B
Experiment 2
Females
Males
NaCl Consumed as % Body Wt
Week
C
Experiment 3
Females
Males
NaCl Consumed as % Body Wt
Week

**S16 Fig. NaCl consumed as a percentage of body weight by context fear conditioned mice across Experiments.**

Mice assigned to 0.4% NaCl represented by blue symbols, mice assigned to 4.0% NaCl represented by red symbols; mice to be tested in Training Context represented by squares and solid lines, mice to be tested in Neutral Context represented by circles and dotted lines. NaCl consumed as a percentage of body weight was calculated for each full week, plus the partial weeks at the conclusion of A) Experiment 1 and B) Experiment 2 (grey shading). Some data loss occurred on the very last weighing day for a subset of animals in C) Experiment 3, thus graphs and repeated measures statistical analyses for Experiment 3 calculations cease at week 6 to maximize inclusion of mice in repeated measures analyses. Experiment 1: 0.4% NaCl females Training Context, n=8; 0.4% NaCl females Neutral Context, n=9; 4.0% NaCl females Training Context, n=8; 4.0% NaCl females Neutral Context, n=9; 0.4% NaCl males Training Context, n=8; 0.4% NaCl males Neutral Context, n=9; 4.0% NaCl males Training Context, n=8; 4.0% NaCl males Neutral Context, n=8. Experiment 2: 0.4% NaCl females Training Context, n=9; 0.4% NaCl females Neutral Context, n=9; 4.0% NaCl females Training Context, n=8; 4.0% NaCl females Neutral Context, n=8; 0.4% NaCl males Training Context, n=9; 0.4% NaCl males Neutral Context, n=9; 4.0% NaCl males Training Context, n=8; 4.0% NaCl males Neutral Context, n=10. Experiment 3: 0.4% NaCl females Training Context, n=8; 0.4% NaCl females Neutral Context, n=9; 4.0% NaCl females Training Context, n=9; 4.0% NaCl females Neutral Context, n=8; 0.4% NaCl males Training Context, n=8; 0.4% NaCl males Neutral Context, n=8; 4.0% NaCl males Training Context, n=8; 4.0% NaCl males Neutral Context, n=8. Data are graphed as mean ± 95% confidence interval. *p<0.05, **p<0.01, ***p<0.001 indicate difference between mice within the same sex consuming 0.4% NaCl versus 4.0% NaCl and tested in Training Context. †p<0.05, ††p<0.01, †††p<0.001 indicate difference between mice within the same sex consuming 0.4% NaCl versus 4.0% NaCl and tested in Neutral Context.

**S25 Table. Three-way repeated measures ANOVAs on weekly ratio of water to NaCl consumed in context fear conditioned mice across Experiments.**

S25A Table

| **Females** | **Experiment 1 – Water:NaCl Ratio** | | |
|---|---|---|---|
| Diet | $F(1,30)=158.3$ | **$p<0.001$** | partial $\eta^2$=**0.841** |
| Context | $F(1,30)=0.020$ | $p=0.890$ | partial $\eta^2=0.001$ |
| Time | $F(1.86,55.66)=1.501$ | $p=0.232$ | partial $\eta^2=0.048$ |
| Time × Diet | $F(1.86,55.66)=1.044$ | $p=0.354$ | partial $\eta^2=0.034$ |
| Time × Context | $F(1.86,55.66)=0.511$ | $p=0.589$ | partial $\eta^2=0.017$ |
| Diet × Context | $F(1,30)=0.000$ | $p=0.996$ | partial $\eta^2=0.000$ |
| Time × Diet × Context | $F(1.86,55.66)=0.264$ | $p=0.752$ | partial $\eta^2=0.009$ |

S25B Table

| **Males** | **Experiment 1 – Water:NaCl Ratio** | | |
|---|---|---|---|
| Diet | $F(1,26)=88.66$ | $p<0.001$ | partial $\eta^2=0.773$ |
| Context | $F(1,26)=0.351$ | $p=0.559$ | partial $\eta^2=0.013$ |
| Time | $F(1.91,49.53)=13.16$ | $p<0.001$ | partial $\eta^2=0.336$ |
| Time × Diet | $F(1.91,49.53)=11.52$ | **$p<0.001$** | partial $\eta^2$=**0.307** |
| Time × Context | $F(1.91,49.53)=0.923$ | $p=0.400$ | partial $\eta^2=0.034$ |
| Diet × Context | $F(1,26)=0.896$ | $p=0.353$ | partial $\eta^2=0.033$ |
| Time × Diet × Context | $F(1.91,49.53)=1.376$ | $p=0.262$ | partial $\eta^2=0.050$ |

S25C Table

| **Females** | **Experiment 2 – Water:NaCl Ratio** | | |
|---|---|---|---|
| Diet | $F(1,29)=417.8$ | $p<0.001$ | partial $\eta^2=0.935$ |
| Context | $F(1,29)=0.839$ | $p=0.367$ | partial $\eta^2=0.028$ |
| Time | $F(3.96,114.7)=4.894$ | $p=0.001$ | partial $\eta^2=0.144$ |
| Time × Diet | $F(3.96,114.7)=4.698$ | **$p=0.002$** | partial $\eta^2$=**0.139** |
| Time × Context | $F(3.96,114.7)=0.952$ | $p=0.436$ | partial $\eta^2=0.032$ |
| Diet × Context | $F(1,29)=1.627$ | $p=0.212$ | partial $\eta^2=0.053$ |
| Time × Diet × Context | $F(3.96,114.7)=0.792$ | $p=0.532$ | partial $\eta^2=0.027$ |

S25D Table

| **Males** | **Experiment 2 – Water:NaCl Ratio** | | |
|---|---|---|---|
| Diet | $F(1,31)=154.6$ | $p<0.001$ | partial $\eta^2=0.833$ |
| Context | $F(1,31)=1.027$ | $p=0.319$ | partial $\eta^2=0.032$ |
| Time | $F(3.70,114.6)=8.018$ | $p<0.001$ | partial $\eta^2=0.205$ |
| Time × Diet | $F(1.17,37.97)=4.476$ | **$p=0.003$** | partial $\eta^2$=**0.126** |
| Time × Context | $F(1.17,37.97)=0.161$ | $p=0.949$ | partial $\eta^2=0.005$ |

| | |
|---|---|
| Diet × Context | F(1,31)=0.615 p=0.439 partial $\eta^2$=0.019 |
| Time × Diet × Context | F(1.17,37.97)=0.170 p=0.945 partial $\eta^2$=0.005 |

S25E Table

| **Females** | **Experiment 3 – Water:NaCl Ratio** |
|---|---|
| Diet | F(1,28)=225.1 p<0.001 partial $\eta^2$=0.889 |
| Context | F(1,28)=0.184 p=0.671 partial $\eta^2$=0.007 |
| Time | F(2.64,73.98)=4.532 p=0.008 partial $\eta^2$=0.139 |
| Time × Diet | F(2.64,73.98)=3.094 **p=0.038** partial $\eta^2$=**0.100** |
| Time × Context | F(2.64,73.98)=0.358 p=0.758 partial $\eta^2$=0.013 |
| Diet × Context | F(1,28)=0.165 p=0.687 partial $\eta^2$=0.006 |
| Time × Diet × Context | F(2.64,73.98)=0.394 p=0.732 partial $\eta^2$=0.014 |

S25F Table

| **Males** | **Experiment 3 – Water:NaCl Ratio** |
|---|---|
| Diet | F(1,27)=212.8 p<0.001 partial $\eta^2$=0.887 |
| Context | F(1,27)=0.999 p=0.326 partial $\eta^2$=0.036 |
| Time | F(2.82,76.01)=8.667 p<0.001 partial $\eta^2$=0.243 |
| Time × Diet | F(2.82,76.01)=4.683 **p=0.006** partial $\eta^2$=**0.148** |
| Time × Context | F(2.82,76.01)=1.027 p=0.382 partial $\eta^2$=0.037 |
| Diet × Context | F(1,27)=1.140 p=0.295 partial $\eta^2$=0.041 |
| Time × Diet × Context | F(2.82,76.01)=0.879 p=0.450 partial $\eta^2$=0.032 |

S17 Figure

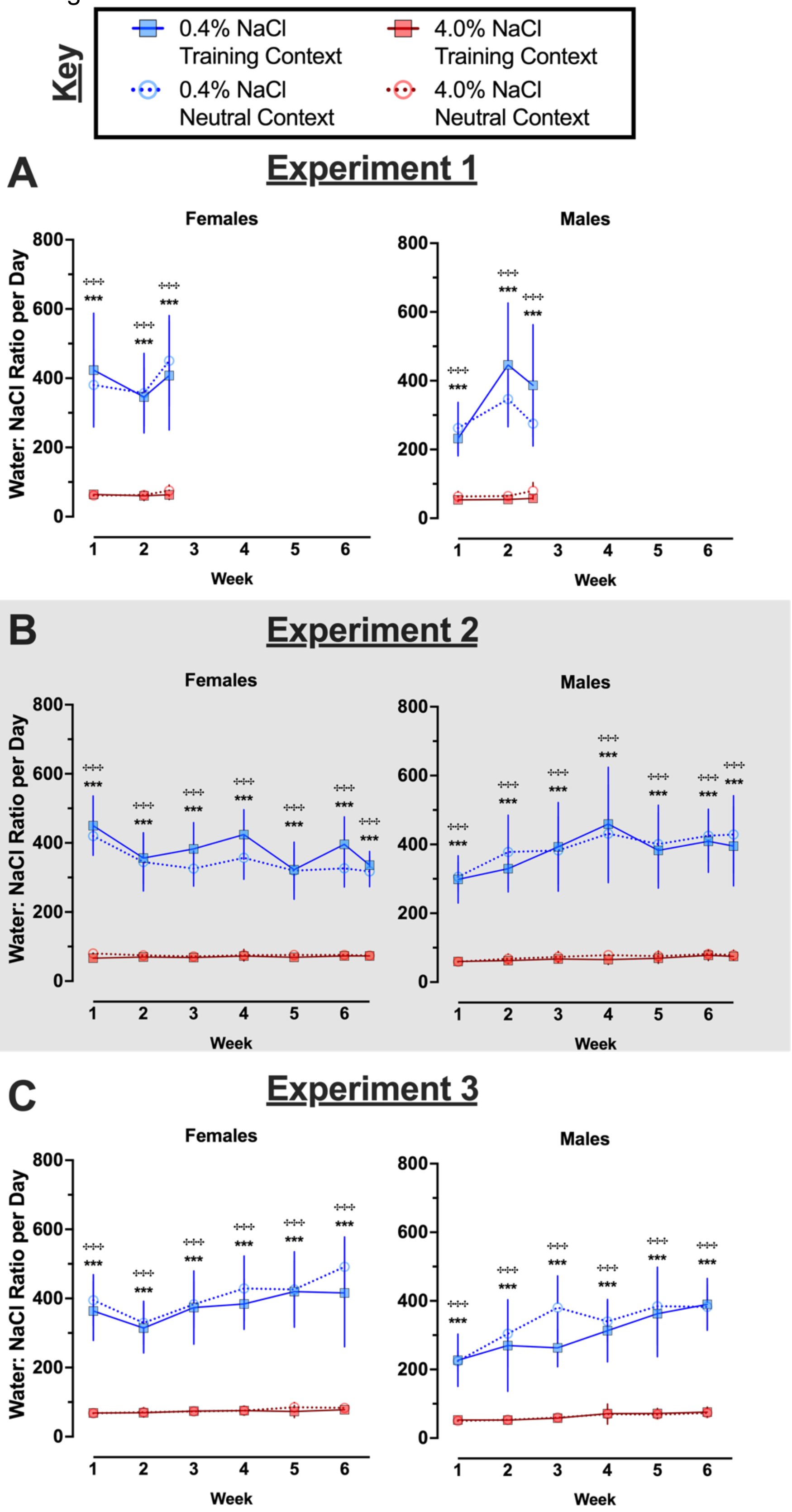
Key
0.4% NaCl Training Context
4.0% NaCl Training Context
0.4% NaCl Neutral Context
4.0% NaCl Neutral Context
A
Experiment 1
Females
Males
Water: NaCl Ratio per Day
Week
B
Experiment 2
Females
Males
Water: NaCl Ratio per Day
Week
C
Experiment 3
Females
Males
Water: NaCl Ratio per Day
Week

**S17 Fig. Water to NaCl consumed ratio for context fear conditioned mice across Experiments.**

Mice assigned to 0.4% NaCl represented by blue symbols, mice assigned to 4.0% NaCl represented by red symbols; mice to be tested in Training Context represented by squares and solid lines, mice to be tested in Neutral Context represented by circles and dotted lines. Water to NaCl consumed ratio was calculated for each full week, plus the partial weeks at the conclusion of A) Experiment 1 and B) Experiment 2 (grey shading). Some data loss occurred on the very last weighing day for a subset of animals in C) Experiment 3, thus graphs and repeated measures statistical analyses for Experiment 3 calculations cease at week 6 to maximize inclusion of mice in repeated measures analyses. Experiment 1: 0.4% NaCl females Training Context, n=8; 0.4% NaCl females Neutral Context, n=9; 4.0% NaCl females Training Context, n=8; 4.0% NaCl females Neutral Context, n=9; 0.4% NaCl males Training Context, n=7; 0.4% NaCl males Neutral Context, n=8; 4.0% NaCl males Training Context, n=8; 4.0% NaCl males Neutral Context, n=7. Experiment 2: 0.4% NaCl females Training Context, n=8; 0.4% NaCl females Neutral Context, n=9; 4.0% NaCl females Training Context, n=8; 4.0% NaCl females Neutral Context, n=8; 0.4% NaCl males Training Context, n=9; 0.4% NaCl males Neutral Context, n=8; 4.0% NaCl males Training Context, n=8; 4.0% NaCl males Neutral Context, n=10. Experiment 3: 0.4% NaCl females Training Context, n=8; 0.4% NaCl females Neutral Context, n=8; 4.0% NaCl females Training Context, n=9; 4.0% NaCl females Neutral Context, n=7; 0.4% NaCl males Training Context, n=8; 0.4% NaCl males Neutral Context, n=7; 4.0% NaCl males Training Context, n=8; 4.0% NaCl males Neutral Context, n=8. Data are graphed as mean ± 95% confidence interval. *p<0.05, **p<0.01, ***p<0.001 indicate difference between mice within the same sex consuming 0.4% NaCl versus 4.0% NaCl and tested in Training Context. $^{+}$p<0.05, $^{++}$p<0.01, $^{+++}$p<0.001 indicate difference between mice within the same sex consuming 0.4% NaCl versus 4.0% NaCl and tested in Neutral Context.

**S26 Table. Three-way repeated measures ANOVAs on weekly average kcal consumed per day by context fear conditioned mice across Experiments.**

S26A Table

| Females | Experiment 1 – kcal/day | | |
|---|---|---|---|
| Diet | $F(1,30)=3.622$ | $p=$**0.067** | partial $\eta^2=$**0.108** |
| Context | $F(1,30)=0.105$ | $p=0.749$ | partial $\eta^2=0.003$ |
| Time | $F(1.67,50.22)=0.640$ | $p=0.505$ | partial $\eta^2=0.021$ |
| Time × Diet | $F(1.67,50.22)=0.568$ | $p=0.540$ | partial $\eta^2=0.019$ |
| Time × Context | $F(1.67,50.22)=1.251$ | $p=0.290$ | partial $\eta^2=0.040$ |
| Diet × Context | $F(1,30)=0.244$ | $p=0.625$ | partial $\eta^2=0.008$ |
| Time × Diet × Context | $F(1.67,50.22)=0.823$ | $p=0.426$ | partial $\eta^2=0.027$ |

S26B Table

| Males | Experiment 1 – kcal/day | | |
|---|---|---|---|
| Diet | $F(1,29)=0.204$ | $p=0.655$ | partial $\eta^2=0.007$ |
| Context | $F(1,29)=0.013$ | $p=0.911$ | partial $\eta^2=0.000$ |
| Time | $F(1.85,53.70)=1.833$ | $p=0.172$ | partial $\eta^2=0.059$ |
| Time × Diet | $F(1.85,53.70)=0.354$ | $p=0.687$ | partial $\eta^2=0.012$ |
| Time × Context | $F(1.85,53.70)=0.148$ | $p=0.847$ | partial $\eta^2=0.005$ |
| Diet × Context | $F(1,29)=0.003$ | $p=0.959$ | partial $\eta^2=0.000$ |
| Time × Diet × Context | $F(1.85,53.70)=0.866$ | $p=0.419$ | partial $\eta^2=0.029$ |

S26C Table

| Females | Experiment 2 – kcal/day | | |
|---|---|---|---|
| Diet | $F(1,30)=27.67$ | **$p<0.001$** | partial $\eta^2=$**0.480** |
| Context | $F(1,30)=0.016$ | $p=0.901$ | partial $\eta^2=0.001$ |
| Time | $F(3.72,111.7)=13.06$ | **$p<0.001$** | partial $\eta^2=$**0.303** |
| Time × Diet | $F(3.72,111.7)=0.221$ | $p=0.916$ | partial $\eta^2=0.007$ |
| Time × Context | $F(3.72,111.7)=1.325$ | $p=0.267$ | partial $\eta^2=0.042$ |
| Diet × Context | $F(1,30)=0.034$ | $p=0.856$ | partial $\eta^2=0.001$ |
| Time × Diet × Context | $F(3.72,111.7)=0.434$ | $p=0.771$ | partial $\eta^2=0.014$ |

S26D Table

| Males | Experiment 2 – kcal/day | | |
|---|---|---|---|
| Diet | $F(1,32)=0.756$ | $p=0.391$ | partial $\eta^2=0.023$ |
| Context | $F(1,32)=0.117$ | $p=0.735$ | partial $\eta^2=0.004$ |
| Time | $F(4.32,138.1)=10.33$ | **$p<0.001$** | partial $\eta^2=$**0.244** |
| Time × Diet | $F(4.32,138.1)=1.334$ | $p=0.258$ | partial $\eta^2=0.040$ |
| Time × Context | $F(4.32,138.1)=0.436$ | $p=0.797$ | partial $\eta^2=0.013$ |

| | |
|---|---|
| Diet × Context | F(1,32)=0.654 p=0.425 partial $\eta^2$=0.020 |
| Time × Diet × Context | F(4.32,138.1)=0.759 p=0.563 partial $\eta^2$=0.023 |

S26E Table

| **Females** | **Experiment 3 – kcal/day** |
|---|---|
| Diet | F(1,30)=10.63 **p=0.003** partial $\eta^2$=**0.262** |
| Context | F(1,30)=1.166 p=0.289 partial $\eta^2$=0.037 |
| Time | F(2.81,84.25)=5.167 **p=0.003** partial $\eta^2$=**0.147** |
| Time × Diet | F(2.81,84.25)=1.684 p=0.180 partial $\eta^2$=0.053 |
| Time × Context | F(2.81,84.25)=0.462 p=0.697 partial $\eta^2$=0.015 |
| Diet × Context | F(1,30)=0.029 p=0.866 partial $\eta^2$=0.001 |
| Time × Diet × Context | F(2.81,84.25)=0.453 p=0.703 partial $\eta^2$=0.015 |

S26F Table

| **Males** | **Experiment 3 – kcal/day** |
|---|---|
| Diet | F(1,28)=4.534 **p=0.042** partial $\eta^2$=**0.139** |
| Context | F(1,28)=0.014 p=0.906 partial $\eta^2$=0.001 |
| Time | F(3.42,95.83)=5.161 **p=0.002** partial $\eta^2$=**0.156** |
| Time × Diet | F(3.42,95.83)=0.990 p=0.408 partial $\eta^2$=0.034 |
| Time × Context | F(3.42,95.83)=1.274 p=0.287 partial $\eta^2$=0.044 |
| Diet × Context | F(1,28)=0.489 p=0.490 partial $\eta^2$=0.017 |
| Time × Diet × Context | F(3.42,95.83)=0.552 p=0.671 partial $\eta^2$=0.019 |

S18 Figure

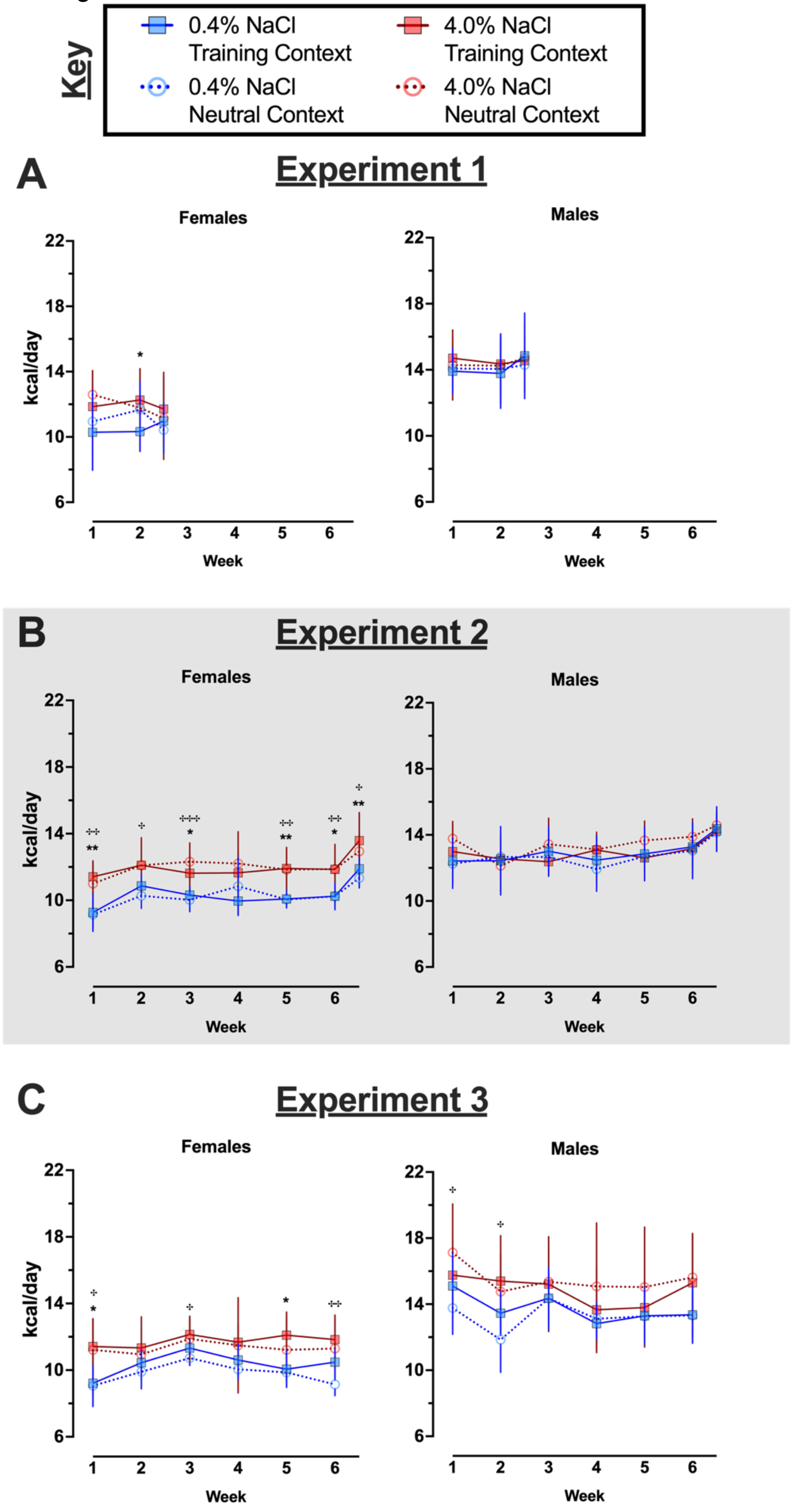

Key
0.4% NaCl Training Context
4.0% NaCl Training Context
0.4% NaCl Neutral Context
4.0% NaCl Neutral Context
A
Experiment 1
Females
Males
kcal/day
Week
B
Experiment 2
Females
Males
kcal/day
Week
C
Experiment 3
Females
Males
kcal/day
Week

**S18 Fig. Average kcal consumed per day by context fear conditioned mice across Experiments.**

Mice assigned to 0.4% NaCl represented by blue symbols, mice assigned to 4.0% NaCl represented by red symbols; mice to be tested in Training Context represented by squares and solid lines, mice to be tested in Neutral Context represented by circles and dotted lines. Consumption of kcal was calculated based upon the amount of each diet consumed per day and the diet's respective kcal (3.89 kcal/g for 0.4% NaCl diet; 3.75 kcal/g for 4.0% NaCl diet). These calculations were made for each full week, plus the partial weeks at the conclusion of A) Experiment 1 and B) Experiment 2 (grey shading). Some data loss occurred on the very last weighing day for a subset of animals in C) Experiment 3, thus graphs and repeated measures statistical analyses for Experiment 3 calculations cease at week 6 to maximize inclusion of mice in repeated measures analyses. Experiment 1: 0.4% NaCl females Training Context, n=8; 0.4% NaCl females Neutral Context, n=9; 4.0% NaCl females Training Context, n=8; 4.0% NaCl females Neutral Context, n=9; 0.4% NaCl males Training Context, n=8; 0.4% NaCl males Neutral Context, n=9; 4.0% NaCl males Training Context, n=8; 4.0% NaCl males Neutral Context, n=8. Experiment 2: 0.4% NaCl females Training Context, n=9; 0.4% NaCl females Neutral Context, n=9; 4.0% NaCl females Training Context, n=8; 4.0% NaCl females Neutral Context, n=8; 0.4% NaCl males Training Context, n=9; 0.4% NaCl males Neutral Context, n=9; 4.0% NaCl males Training Context, n=8; 4.0% NaCl males Neutral Context, n=10. Experiment 3: 0.4% NaCl females Training Context, n=8; 0.4% NaCl females Neutral Context, n=9; 4.0% NaCl females Training Context, n=9; 4.0% NaCl females Neutral Context, n=8; 0.4% NaCl males Training Context, n=8; 0.4% NaCl males Neutral Context, n=8; 4.0% NaCl males Training Context, n=8; 4.0% NaCl males Neutral Context, n=8. Data are graphed as mean ± 95% confidence interval. *$p<0.05$, **$p<0.01$, ***$p<0.001$ indicate difference between mice within the same sex consuming 0.4% NaCl versus 4.0% NaCl and tested in Training Context. ‡$p<0.05$,

‡‡p<0.01, ‡‡‡p<0.001 indicate difference between mice within the same sex consuming 0.4% NaCl versus 4.0% NaCl and tested in Neutral Context.

**S27 Table. Three-way repeated measures ANOVAs on weekly kcal consumed as a percentage of body weight in context fear conditioned mice across Experiments.**

S27A Table

| **Females** | **Experiment 1 – kcal as % BW** |
|---|---|
| Diet | $F(1,30)=2.043$ $p=0.163$ partial $\eta^2=0.064$ |
| Context | $F(1,30)=0.320$ $p=0.576$ partial $\eta^2=0.011$ |
| Time | $F(1.61,48.41)=3.279$ $p=0.056$ partial $\eta^2=0.099$ |
| Time × Diet | $F(1.61,48.41)=0.462$ $p=0.591$ partial $\eta^2=0.015$ |
| Time × Context | $F(1.61,48.41)=1.166$ $p=0.311$ partial $\eta^2=0.037$ |
| Diet × Context | $F(1,30)=0.140$ $p=0.710$ partial $\eta^2=0.005$ |
| Time × Diet × Context | $F(1.61,48.41)=0.756$ $p=0.448$ partial $\eta^2=0.025$ |

S27B Table

| **Males** | **Experiment 1 – kcal as % BW** |
|---|---|
| Diet | $F(1,29)=0.000$ $p=0.993$ partial $\eta^2=0.000$ |
| Context | $F(1,29)=0.006$ $p=0.941$ partial $\eta^2=0.000$ |
| Time | $F(1.71,49.67)=0.479$ $p=0.593$ partial $\eta^2=0.016$ |
| Time × Diet | $F(1.71,49.67)=0.459$ $p=0.605$ partial $\eta^2=0.016$ |
| Time × Context | $F(1.71,49.67)=0.272$ $p=0.729$ partial $\eta^2=0.009$ |
| Diet × Context | $F(1,29)=0.024$ $p=0.878$ partial $\eta^2=0.001$ |
| Time × Diet × Context | $F(1.71,49.67)=0.631$ $p=0.512$ partial $\eta^2=0.021$ |

S27C Table

| **Females** | **Experiment 2 – kcal as % BW** |
|---|---|
| Diet | $F(1,30)=25.14$ $\mathbf{p<0.001}$ partial $\eta^2=\mathbf{0.456}$ |
| Context | $F(1,30)=0.003$ $p=0.953$ partial $\eta^2=0.000$ |
| Time | $F(3.40,102.1)=5.508$ $\mathbf{p<0.001}$ partial $\eta^2=\mathbf{0.155}$ |
| Time × Diet | $F(3.40,102.1)=0.541$ $p=0.677$ partial $\eta^2=0.018$ |
| Time × Context | $F(3.40,102.1)=1.330$ $p=0.267$ partial $\eta^2=0.042$ |
| Diet × Context | $F(1,30)=0.428$ $p=0.518$ partial $\eta^2=0.014$ |
| Time × Diet × Context | $F(3.40,102.1)=0.487$ $p=0.487$ partial $\eta^2=0.016$ |

S27D Table

| **Males** | **Experiment 2 – kcal as % BW** |
|---|---|
| Diet | $F(1,32)=0.793$ $p=0.380$ partial $\eta^2=0.024$ |
| Context | $F(1,32)=0.424$ $p=0.520$ partial $\eta^2=0.013$ |
| Time | $F(3.71,118.6)=6.216$ $\mathbf{p<0.001}$ partial $\eta^2=\mathbf{0.163}$ |
| Time × Diet | $F(3.71,118.6)=1.178$ $p=0.324$ partial $\eta^2=0.036$ |
| Time × Context | $F(3.71,118.6)=0.414$ $p=0.784$ partial $\eta^2=0.013$ |

| | | | |
|---|---|---|---|
| Diet × Context | $F(1,32)=2.189$ | $p=0.149$ | partial $\eta^2=0.064$ |
| Time × Diet × Context | $F(3.71,118.6)=0.413$ | $p=0.785$ | partial $\eta^2=0.013$ |

S27E Table

| **Females** | **Experiment 3 – kcal as % BW** | | |
|---|---|---|---|
| Diet | $F(1,30)=6.312$ | **p=0.018** | partial $\eta^2=$**0.174** |
| Context | $F(1,30)=0.005$ | $p=0.947$ | partial $\eta^2=0.000$ |
| Time | $F(2.59,77.66)=5.137$ | **p=0.004** | partial $\eta^2=$**0.146** |
| Time × Diet | $F(2.59,77.66)=1.132$ | $p=0.337$ | partial $\eta^2=0.036$ |
| Time × Context | $F(2.59,77.66)=0.494$ | $p=0.660$ | partial $\eta^2=0.016$ |
| Diet × Context | $F(1,30)=1.458$ | $p=0.237$ | partial $\eta^2=0.046$ |
| Time × Diet × Context | $F(2.59,77.66)=0.217$ | $p=0.858$ | partial $\eta^2=0.007$ |

S27F Table

| **Males** | **Experiment 3 – kcal as % BW** | | |
|---|---|---|---|
| Diet | $F(1,28)=12.08$ | **p=0.002** | partial $\eta^2=$**0.301** |
| Context | $F(1,28)=1.490$ | $p=0.232$ | partial $\eta^2=0.051$ |
| Time | $F(3.06,85.77)=8.143$ | **p<0.001** | partial $\eta^2=$**0.225** |
| Time × Diet | $F(3.06,85.77)=0.870$ | $p=0.462$ | partial $\eta^2=0.030$ |
| Time × Context | $F(3.06,85.77)=1.053$ | $p=0.374$ | partial $\eta^2=0.036$ |
| Diet × Context | $F(1,28)=0.277$ | $p=0.603$ | partial $\eta^2=0.010$ |
| Time × Diet × Context | $F(3.06,85.77)=0.509$ | $p=0.681$ | partial $\eta^2=0.018$ |

S19 Figure

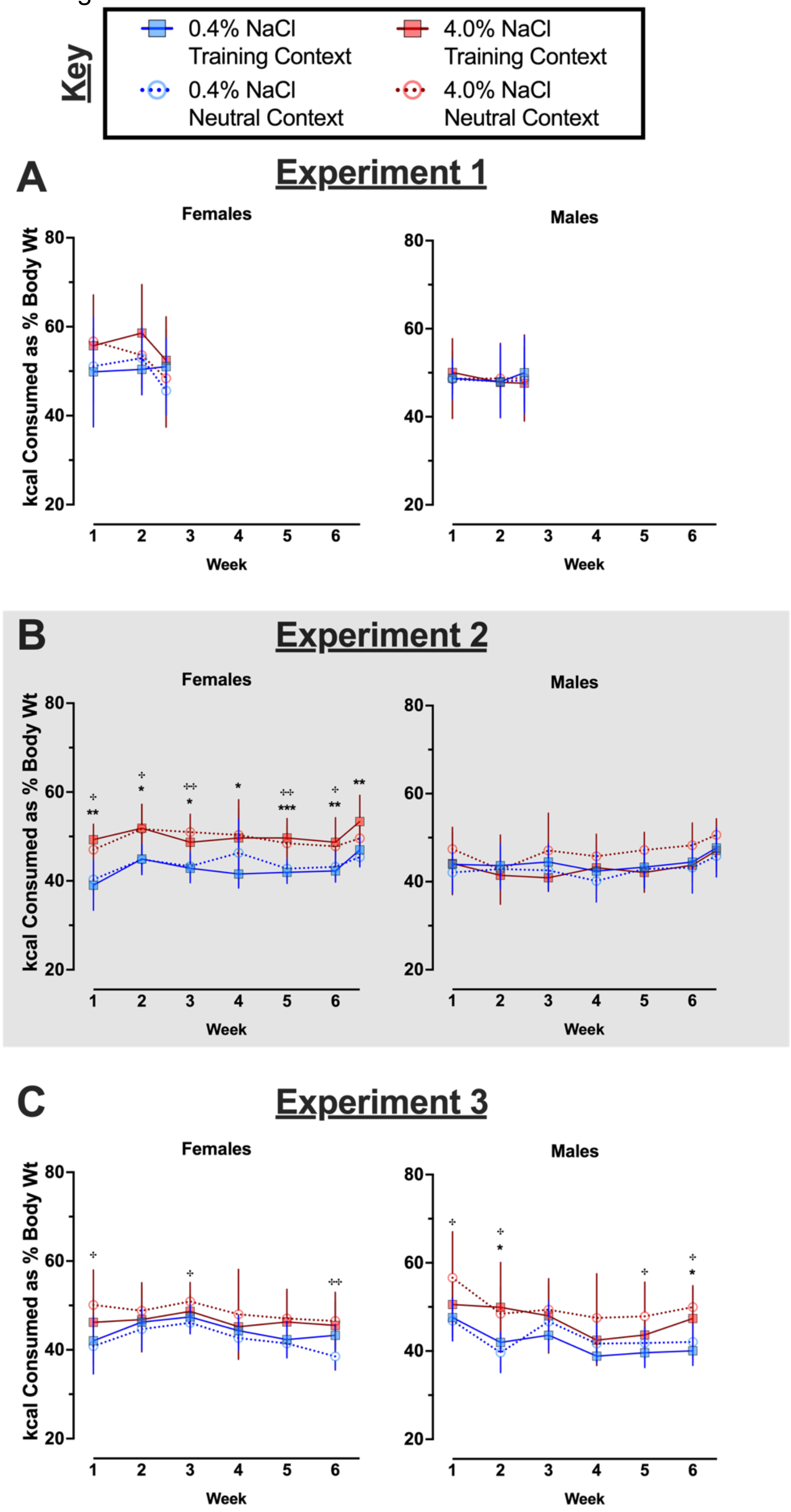
Key
0.4% NaCl Training Context
4.0% NaCl Training Context
0.4% NaCl Neutral Context
4.0% NaCl Neutral Context
A
Experiment 1
Females
Males
kcal Consumed as % Body Wt
Week
B
Experiment 2
Females
Males
kcal Consumed as % Body Wt
Week
C
Experiment 3
Females
Males
kcal Consumed as % Body Wt
Week

**S19 Fig. Consumed kcal as a percentage of body weight by context fear conditioned mice across Experiments.**

Mice assigned to 0.4% NaCl represented by blue symbols, mice assigned to 4.0% NaCl represented by red symbols; mice to be tested in Training Context represented by squares and solid lines, mice to be tested in Neutral Context represented by circles and dotted lines. Consumed kcal as a percentage of body weight was calculated for each full week, plus the partial weeks at the conclusion of A) Experiment 1 and B) Experiment 2 (grey shading). Some data loss occurred on the very last weighing day for a subset of animals in C) Experiment 3, thus graphs and repeated measures statistical analyses for Experiment 3 calculations cease at week 6 to maximize inclusion of mice in repeated measures analyses. Experiment 1: 0.4% NaCl females Training Context, n=8; 0.4% NaCl females Neutral Context, n=9; 4.0% NaCl females Training Context, n=8; 4.0% NaCl females Neutral Context, n=9; 0.4% NaCl males Training Context, n=8; 0.4% NaCl males Neutral Context, n=9; 4.0% NaCl males Training Context, n=8; 4.0% NaCl males Neutral Context, n=8. Experiment 2: 0.4% NaCl females Training Context, n=9; 0.4% NaCl females Neutral Context, n=9; 4.0% NaCl females Training Context, n=8; 4.0% NaCl females Neutral Context, n=8; 0.4% NaCl males Training Context, n=9; 0.4% NaCl males Neutral Context, n=9; 4.0% NaCl males Training Context, n=8; 4.0% NaCl males Neutral Context, n=10. Experiment 3: 0.4% NaCl females Training Context, n=8; 0.4% NaCl females Neutral Context, n=9; 4.0% NaCl females Training Context, n=9; 4.0% NaCl females Neutral Context, n=8; 0.4% NaCl males Training Context, n=8; 0.4% NaCl males Neutral Context, n=8; 4.0% NaCl males Training Context, n=8; 4.0% NaCl males Neutral Context, n=8. Data are graphed as mean ± 95% confidence interval. *p<0.05, **p<0.01, ***p<0.001 indicate difference between mice within the same sex consuming 0.4% NaCl versus 4.0% NaCl and tested in Training Context. $^{+}$p<0.05, $^{++}$p<0.01, $^{+++}$p<0.001 indicate difference

between mice within the same sex consuming 0.4% NaCl versus 4.0% NaCl and tested in Neutral Context.

**S28 Table. Three-way repeated measures ANOVAs on weekly average NaCl consumed per day by control no shock mice across Experiments.**

S28A Table

| **Experiment 1** | **NaCl/day** | | |
|---|---|---|---|
| Sex | $F(1,31)=6.870$ | $p=0.013$ | partial $\eta^2=0.181$ |
| Diet | $F(1,31)=585.5$ | $p<0.001$ | partial $\eta^2=0.950$ |
| Time | $F(1.55,47.89)=0.069$ | $p=0.890$ | partial $\eta^2=0.002$ |
| Time × Sex | $F(1.55,47.89)=1.907$ | $p=0.168$ | partial $\eta^2=0.058$ |
| Time × Diet | $F(1.55,47.89)=0.103$ | $p=0.853$ | partial $\eta^2=0.003$ |
| Sex × Diet | $F(1,31)=4.262$ | **$p=0.047$** | partial $\eta^2=$**0.121** |
| Time × Sex × Diet | $F(1.55,47.89)=1.733$ | $p=0.193$ | partial $\eta^2=0.053$ |

S28B Table

| **Experiment 2** | **NaCl/day** | | |
|---|---|---|---|
| Sex | $F(1,29)=14.80$ | $p<0.001$ | partial $\eta^2=0.338$ |
| Diet | $F(1,29)=1821$ | $p<0.001$ | partial $\eta^2=0.984$ |
| Time | $F(2.40,69.59)=3.232$ | $p=0.037$ | partial $\eta^2=0.100$ |
| Time × Sex | $F(2.40,69.59)=0.157$ | $p=0.889$ | partial $\eta^2=0.005$ |
| Time × Diet | $F(2.40,69.59)=3.330$ | **$p=0.033$** | partial $\eta^2=$**0.103** |
| Sex × Diet | $F(1,29)=11.10$ | **$p=0.002$** | partial $\eta^2=$**0.277** |
| Time × Sex × Diet | $F(2.40,69.59)=0.132$ | $p=0.909$ | partial $\eta^2=0.005$ |

S28C Table

| **Experiment 3** | **NaCl/day** | | |
|---|---|---|---|
| Sex | $F(1,28)=6.847$ | $p=0.014$ | partial $\eta^2=0.196$ |
| Diet | $F(1,28)=318.3$ | $p<0.001$ | partial $\eta^2=0.919$ |
| Time | $F(2.37,66.44)=0.759$ | $p=0.493$ | partial $\eta^2=0.026$ |
| Time × Sex | $F(2.37,66.44)=1.633$ | $p=0.199$ | partial $\eta^2=0.055$ |
| Time × Diet | $F(2.37,66.44)=0.729$ | $p=0.508$ | partial $\eta^2=0.025$ |
| Sex × Diet | $F(1,28)=3.864$ | **$p=0.059$** | partial $\eta^2=$**0.121** |
| Time × Sex × Diet | $F(2.37,66.44)=1.580$ | $p=0.210$ | partial $\eta^2=0.053$ |

**S29 Table. Three-way repeated measures ANOVAs on weekly NaCl consumed as a percentage of body weight by control no shock mice across Experiments.**

S29A Table

| **Experiment 1** | **NaCl as % BW** | | |
|---|---|---|---|
| Sex | $F(1,31)=0.304$ | $p=0.585$ | partial $\eta^2=0.010$ |
| Diet | $F(1,31)=323.9$ | $p<0.001$ | partial $\eta^2=0.913$ |
| Time | $F(1.52,47.12)=0.936$ | $p=0.376$ | partial $\eta^2=0.029$ |
| Time × Sex | $F(1.52,47.12)=3.195$ | $p=0.063$ | partial $\eta^2=0.093$ |
| Time × Diet | $F(1.52,47.12)=0.743$ | $p=0.447$ | partial $\eta^2=0.023$ |
| Sex × Diet | $F(1,31)=0.329$ | $p=0.570$ | partial $\eta^2=0.011$ |
| Time × Sex × Diet | $F(1.52,47.12)=2.829$ | **$p=0.083$** | partial **$\eta^2=0.084$** |

S29B Table

| **Experiment 2** | **NaCl as % BW** | | |
|---|---|---|---|
| Sex | $F(1,29)=4.453$ | $p=0.044$ | partial $\eta^2=0.133$ |
| Diet | $F(1,29)=2475$ | $p<0.001$ | partial $\eta^2=0.988$ |
| Time | $F(1.92,55.69)=2.794$ | $p=0.072$ | partial $\eta^2=0.088$ |
| Time × Sex | $F(1.92,55.69)=0.109$ | $p=0.889$ | partial $\eta^2=0.004$ |
| Time × Diet | $F(1.92,55.69)=3.222$ | **$p=0.049$** | partial $\eta^2=$**$0.100$** |
| Sex × Diet | $F(1,29)=3.125$ | **$p=0.088$** | partial $\eta^2=$**$0.097$** |
| Time × Sex × Diet | $F(1.92,55.69)=0.074$ | $p=0.923$ | partial $\eta^2=0.003$ |

S29C Table

| **Experiment 3** | **NaCl as % BW** | | |
|---|---|---|---|
| Sex | $F(1,28)=0.211$ | $p=0.649$ | partial $\eta^2=0.007$ |
| Diet | $F(1,28)=438.4$ | **$p<0.001$** | partial $\eta^2=$**$0.940$** |
| Time | $F(2.91,81.55)=1.500$ | $p=0.222$ | partial $\eta^2=0.051$ |
| Time × Sex | $F(2.91,81.55)=1.040$ | $p=0.378$ | partial $\eta^2=0.036$ |
| Time × Diet | $F(2.91,81.55)=1.330$ | $p=0.271$ | partial $\eta^2=0.045$ |
| Sex × Diet | $F(1,28)=0.521$ | $p=0.476$ | partial $\eta^2=0.018$ |
| Time × Sex × Diet | $F(2.91,81.55)=0.940$ | $p=0.423$ | partial $\eta^2=0.032$ |

S20 Figure

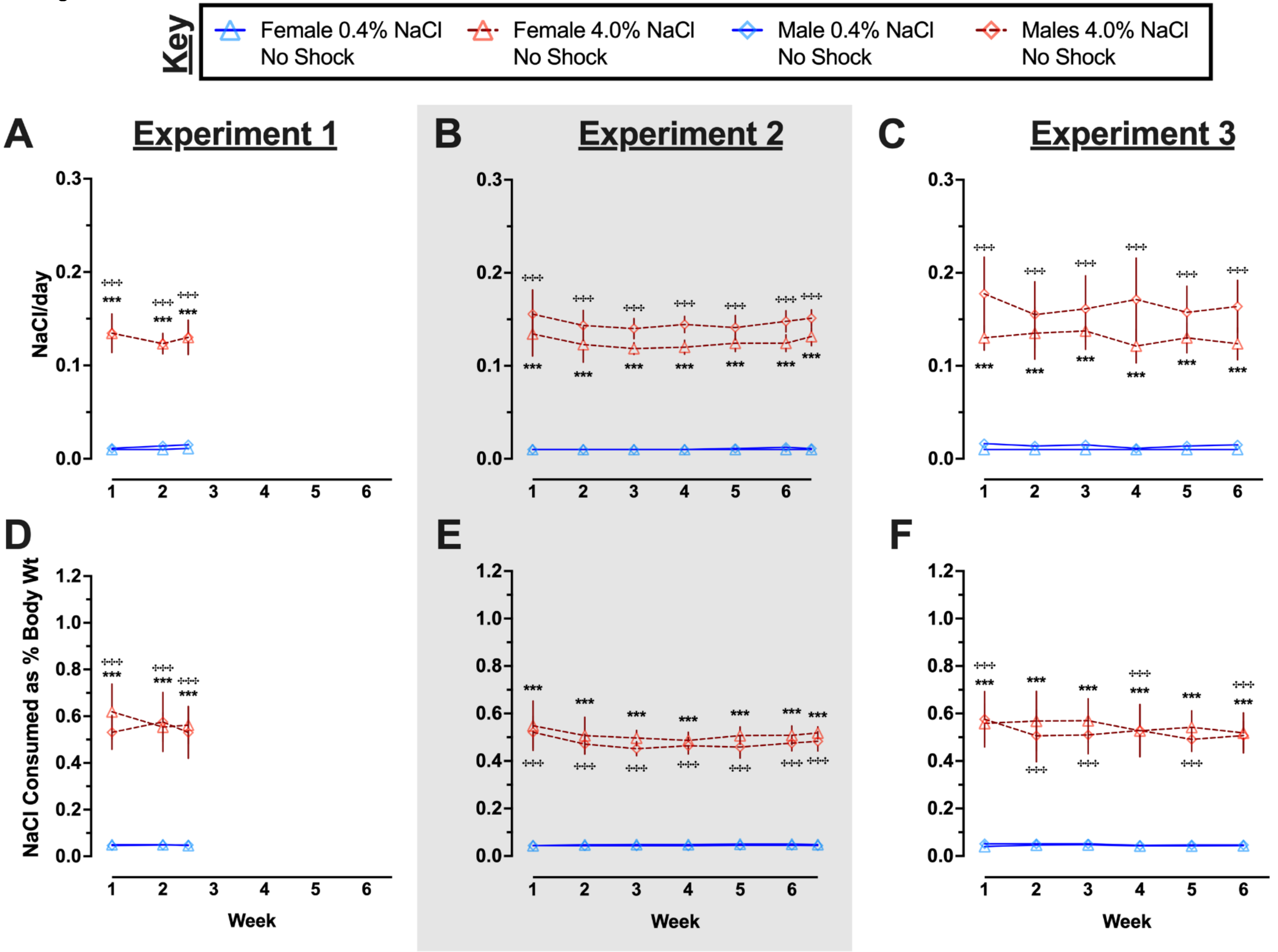

Key
Female 0.4% NaCl No Shock
Female 4.0% NaCl No Shock
Male 0.4% NaCl No Shock
Males 4.0% NaCl No Shock
A
Experiment 1
B
Experiment 2
C
Experiment 3
D
E
F
NaCl/day
NaCl Consumed as % Body Wt
Week

**S20 Fig. Average NaCl consumed per day and NaCl consumed as a percentage of body weight by control no shock mice across Experiments.**

Females represented by triangles, males by diamonds; 0.4% NaCl represented by blue symbols and solid lines, 4.0% NaCl represented by red symbols and dashed lines. A, B, C) NaCl consumed per day and D, E, F) NaCl consumption as a percentage of body weight were calculated for each full week, and for the partial week at the conclusion of A, D) Experiment 1 and B, E) Experiment 2 (grey shading). Some data loss occurred on the very last weighing day for a subset of animals in C, F) Experiment 3, thus graphs and repeated measures statistical analyses for Experiment 3 calculations cease at week 6 to maximize inclusion of mice in repeated measures analyses. Experiment 1: 0.4% NaCl females, n=9; 4.0% NaCl females, n=9; 0.4% NaCl males, n=8; 4.0% NaCl males, n=9. Experiment 2: 0.4% NaCl females, n=8; 4.0% NaCl females, n=7; 0.4% NaCl males, n=9; 4.0% NaCl males, n=9. Experiment 3: 0.4% NaCl females, n=8; 4.0% NaCl females, n=8; 0.4% NaCl males, n=8; 4.0% NaCl males, n=8. Data are graphed as mean ± 95% confidence interval. *$p<0.05$, **$p<0.01$, ***$p<0.001$ indicate difference between females consuming 0.4% NaCl versus 4.0% NaCl. $^{\ddagger}p<0.05$, $^{\ddagger\ddagger}p<0.01$, $^{\ddagger\ddagger\ddagger}p<0.001$ indicate difference between males consuming 0.4% NaCl versus 4.0% NaCl.

**S30 Table. Three-way repeated measures ANOVAs on weekly water to NaCl ratio consumed by control no shock mice across Experiments.**

S30A Table

| **Experiment 1** | **Water:NaCl Ratio** | | |
|---|---|---|---|
| Sex | $F(1,31)=13.68$ | $p<0.001$ | partial $\eta^2=0.306$ |
| Diet | $F(1,31)=487.2$ | $p<0.001$ | partial $\eta^2=0.940$ |
| Time | $F(1.39,43.09)=3.731$ | $\mathbf{p=0.047}$ | partial $\eta^2=\mathbf{0.107}$ |
| Time × Sex | $F(1.39,43.09)=2.248$ | $p=0.133$ | partial $\eta^2=0.068$ |
| Time × Diet | $F(1.39,43.09)=3.027$ | $p=0.076$ | partial $\eta^2=0.089$ |
| Sex × Diet | $F(1,31)=18.22$ | $\mathbf{p<0.001}$ | partial $\eta^2=\mathbf{0.370}$ |
| Time × Sex × Diet | $F(1.39,43.09)=2.048$ | $p=0.154$ | partial $\eta^2=0.062$ |

S30B Table

| **Experiment 2** | **Water:NaCl Ratio** | | |
|---|---|---|---|
| Sex | $F(1,29)=1.913$ | $p=0.177$ | partial $\eta^2=0.062$ |
| Diet | $F(1,29)=589.6$ | $p<0.001$ | partial $\eta^2=0.953$ |
| Time | $F(3.93,113.8)=2.126$ | $p=0.083$ | partial $\eta^2=0.068$ |
| Time × Sex | $F(3.93,113.8)=5.567$ | $p<0.001$ | partial $\eta^2=0.161$ |
| Time × Diet | $F(3.93,113.8)=1.161$ | $p=0.332$ | partial $\eta^2=0.038$ |
| Sex × Diet | $F(1,29)=2.208$ | $p=0.148$ | partial $\eta^2=0.071$ |
| Time × Sex × Diet | $F(3.93,113.8)=4.342$ | $\mathbf{p=0.003}$ | partial $\eta^2=\mathbf{0.130}$ |

S30C Table

| **Experiment 3** | **Water:NaCl Ratio** | | |
|---|---|---|---|
| Sex | $F(1,28)=2.707$ | $p=0.111$ | partial $\eta^2=0.088$ |
| Diet | $F(1,28)=137.4$ | $p<0.001$ | partial $\eta^2=0.831$ |
| Time | $F(2.85,79.82)=1.662$ | $p=0.184$ | partial $\eta^2=0.056$ |
| Time × Sex | $F(2.85,79.82)=4.474$ | $p=0.007$ | partial $\eta^2=0.138$ |
| Time × Diet | $F(2.85,79.82)=1.050$ | $p=0.373$ | partial $\eta^2=0.036$ |
| Sex × Diet | $F(1,28)=0.347$ | $p=0.561$ | partial $\eta^2=0.012$ |
| Time × Sex × Diet | $F(2.85,79.82)=4.428$ | $\mathbf{p=0.007}$ | partial $\eta^2=\mathbf{0.137}$ |

S21 Figure

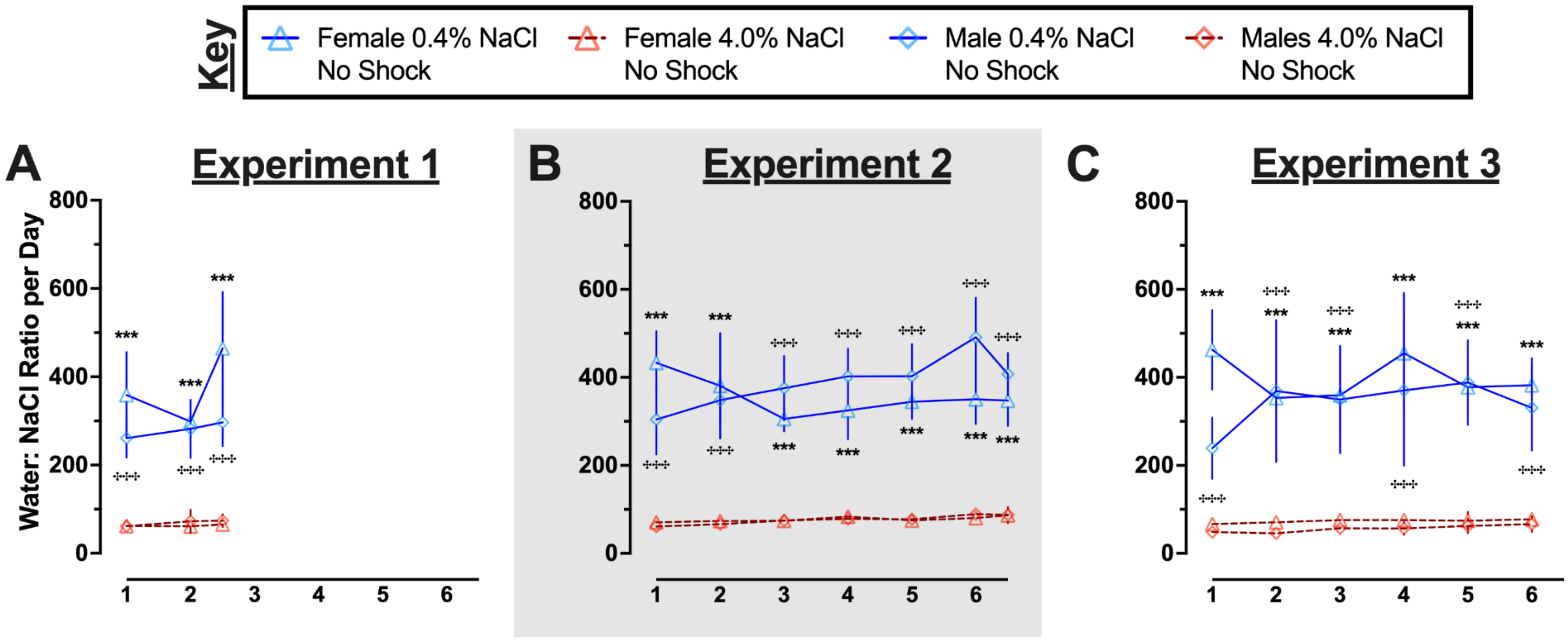
Key
Female 0.4% NaCl No Shock
Female 4.0% NaCl No Shock
Male 0.4% NaCl No Shock
Males 4.0% NaCl No Shock
A
Experiment 1
B
Experiment 2
C
Experiment 3
Water: NaCl Ratio per Day
800
600
400
200
0
1
2
3
4
5
6
***

**S21 Fig. Ratio of water to NaCl consumed by control no shock mice across Experiments.** Females represented by triangles, males by diamonds; 0.4% NaCl represented by blue symbols and solid lines, 4.0% NaCl represented by red symbols and dashed lines. Water to NaCl consumption ratio was calculated for each full week, and for the partial week at the conclusion of A) Experiment 1 and B) Experiment 2 (grey shading). Some data loss occurred on the very last weighing day for a subset of animals in C) Experiment 3, thus graphs and repeated measures statistical analyses for Experiment 3 calculations cease at week 6 to maximize inclusion of mice in repeated measures analyses. Experiment 1: 0.4% NaCl females, n=9; 4.0% NaCl females, n=9; 0.4% NaCl males, n=8; 4.0% NaCl males, n=9. Experiment 2: 0.4% NaCl females, n=8; 4.0% NaCl females, n=7; 0.4% NaCl males, n=9; 4.0% NaCl males, n=9. Experiment 3: 0.4% NaCl females, n=8; 4.0% NaCl females, n=8; 0.4% NaCl males, n=8; 4.0% NaCl males, n=8. Data are graphed as mean ± 95% confidence interval. *p<0.05, **p<0.01, ***p<0.001 indicate difference between females consuming 0.4% NaCl versus 4.0% NaCl. ‡p<0.05, ‡‡p<0.01, ‡‡‡p<0.001 indicate difference between males consuming 0.4% NaCl versus 4.0% NaCl.

**S31 Table. Three-way repeated measures ANOVAs on weekly average kcal consumed per day by control no shock mice across Experiments.**

S31A Table

| Experiment 1 | kcal/day | | |
|---|---|---|---|
| Sex | $F(1,31)=17.84$ | **$p<0.001$** | partial $\eta^2$=**0.365** |
| Diet | $F(1,31)=4.655$ | **$p=0.039$** | partial $\eta^2$=**0.131** |
| Time | $F(1.55,48.15)=0.003$ | $p=0.990$ | partial $\eta^2=0.000$ |
| Time × Sex | $F(1.55,48.15)=2.408$ | $p=0.113$ | partial $\eta^2=0.072$ |
| Time × Diet | $F(1.55,48.15)=0.225$ | $p=0.743$ | partial $\eta^2=0.007$ |
| Sex × Diet | $F(1,31)=0.193$ | $p=0.663$ | partial $\eta^2=0.006$ |
| Time × Sex × Diet | $F(1.55,48.15)=1.253$ | $p=0.288$ | partial $\eta^2=0.039$ |

S31B Table

| Experiment 2 | kcal/day | | |
|---|---|---|---|
| Sex | $F(1,29)=20.20$ | **$p<0.001$** | partial $\eta^2$=**0.411** |
| Diet | $F(1,29)=6.500$ | $p=0.016$ | partial $\eta^2=0.183$ |
| Time | $F(3.19,92.45)=3.755$ | $p=0.012$ | partial $\eta^2=0.115$ |
| Time × Sex | $F(3.19,92.45)=0.524$ | $p=0.678$ | partial $\eta^2=0.018$ |
| Time × Diet | $F(3.19,92.45)=4.385$ | **$p=0.005$** | partial $\eta^2$=**0.131** |
| Sex × Diet | $F(1,29)=0.433$ | $p=0.516$ | partial $\eta^2=0.015$ |
| Time × Sex × Diet | $F(3.19,92.45)=0.359$ | $p=0.795$ | partial $\eta^2=0.012$ |

S31C Table

| Experiment 3 | kcal/day | | |
|---|---|---|---|
| Sex | $F(1,28)=24.96$ | **$p<0.001$** | partial $\eta^2$=**0.471** |
| Diet | $F(1,28)=5.135$ | **$p=0.031$** | partial $\eta^2$=**0.155** |
| Time | $F(2.77,77.66)=1.733$ | $p=0.171$ | partial $\eta^2=0.058$ |
| Time × Sex | $F(2.77,77.66)=1.512$ | $p=0.220$ | partial $\eta^2=0.051$ |
| Time × Diet | $F(2.77,77.66)=1.539$ | $p=0.214$ | partial $\eta^2=0.052$ |
| Sex × Diet | $F(1,28)=0.916$ | $p=0.347$ | partial $\eta^2=0.032$ |
| Time × Sex × Diet | $F(2.77,77.66)=1.162$ | $p=0.328$ | partial $\eta^2=0.040$ |

**S32 Table. Three-way repeated measures ANOVAs on weekly average kcal consumed as a percentage of body weight by control no shock mice across Experiments.**

S32A Table

| **Experiment 1** | **Kcal as % BW** |
|---|---|
| Sex | $F(1,31)=0.151$ $p=0.700$ partial $\eta^2=0.005$ |
| Diet | $F(1,31)=3.279$ $p=0.080$ partial $\eta^2=0.096$ |
| Time | $F(1.65,51.04)=1.353$ $p=0.265$ partial $\eta^2=0.042$ |
| Time × Sex | $F(1.65,51.04)=3.592$ $\mathbf{p=0.043}$ partial $\eta^2=\mathbf{0.104}$ |
| Time × Diet | $F(1.65,51.04)=0.089$ $p=0.880$ partial $\eta^2=0.003$ |
| Sex × Diet | $F(1,31)=0.348$ $p=0.560$ partial $\eta^2=0.011$ |
| Time × Sex × Diet | $F(1.65,51.04)=1.194$ $p=0.304$ partial $\eta^2=0.037$ |

S32B Table

| **Experiment 2** | **Kcal as % BW** |
|---|---|
| Sex | $F(1,29)=7.328$ $\mathbf{p=0.011}$ partial $\eta^2=\mathbf{0.202}$ |
| Diet | $F(1,29)=1.017$ $p=0.322$ partial $\eta^2=0.034$ |
| Time | $F(2.59,75.20)=1.345$ $p=0.267$ partial $\eta^2=0.044$ |
| Time × Sex | $F(2.59,75.20)=0.667$ $p=0.554$ partial $\eta^2=0.022$ |
| Time × Diet | $F(2.59,75.20)=4.208$ $\mathbf{p=0.011}$ partial $\eta^2=\mathbf{0.127}$ |
| Sex × Diet | $F(1,29)=0.014$ $p=0.906$ partial $\eta^2=0.000$ |
| Time × Sex × Diet | $F(2.59,75.20)=0.430$ $p=0.704$ partial $\eta^2=0.015$ |

S32C Table

| **Experiment 3** | **Kcal as % BW** |
|---|---|
| Sex | $F(1,28)=0.431$ $p=0.517$ partial $\eta^2=0.015$ |
| Diet | $F(1,28)=4.291$ $\mathbf{p=0.048}$ partial $\eta^2=\mathbf{0.133}$ |
| Time | $F(3.02,84.44)=3.121$ $\mathbf{p=0.030}$ partial $\eta^2=\mathbf{0.100}$ |
| Time × Sex | $F(3.02,84.44)=1.336$ $p=0.268$ partial $\eta^2=0.046$ |
| Time × Diet | $F(3.02,84.44)=1.945$ $p=0.128$ partial $\eta^2=0.065$ |
| Sex × Diet | $F(1,28)=2.776$ $p=0.107$ partial $\eta^2=0.090$ |
| Time × Sex × Diet | $F(3.02,84.44)=0.648$ $p=0.587$ partial $\eta^2=0.023$ |

S22 Figure

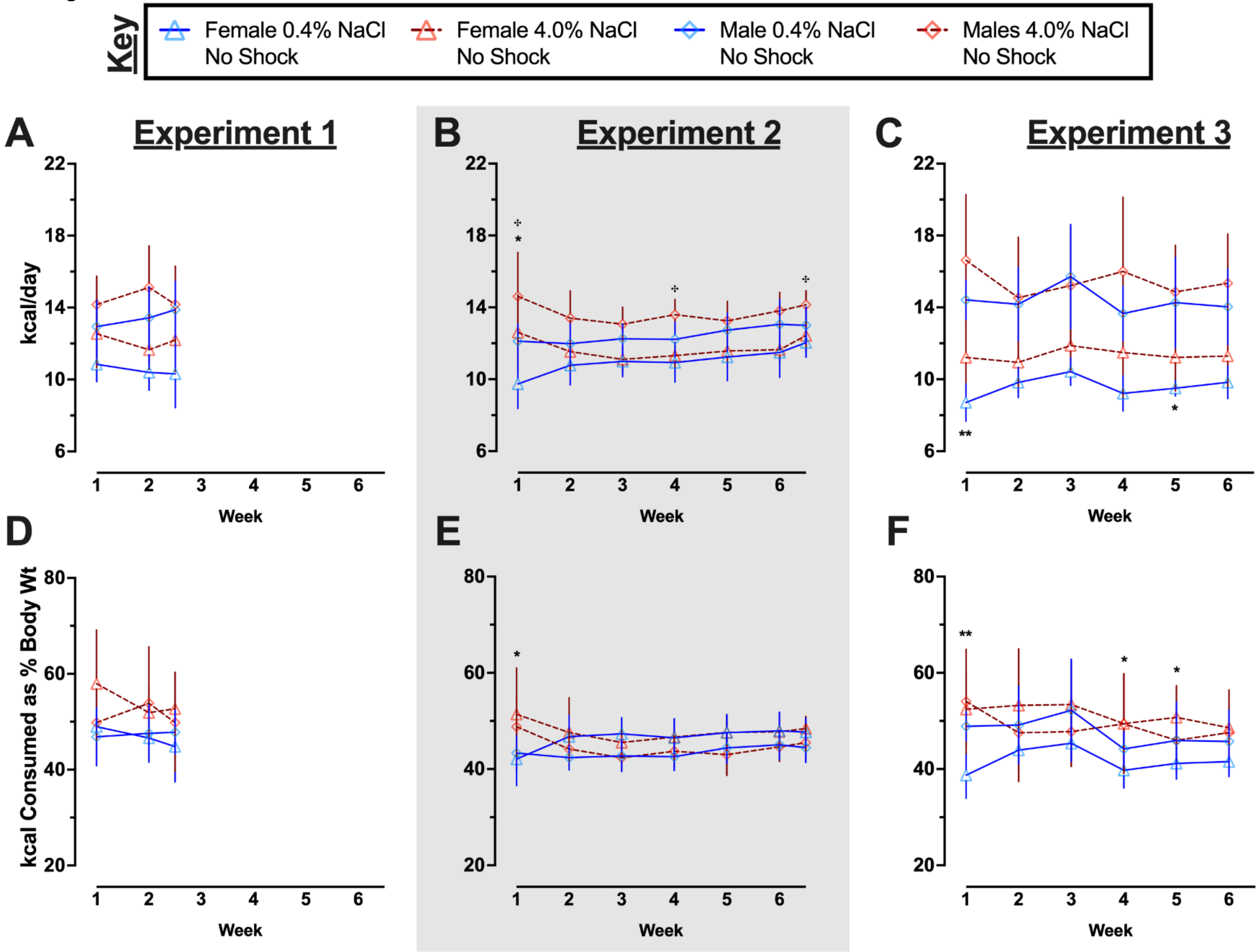
Key
Female 0.4% NaCl No Shock
Female 4.0% NaCl No Shock
Male 0.4% NaCl No Shock
Males 4.0% NaCl No Shock
A
Experiment 1
B
Experiment 2
C
Experiment 3
kcal/day
22
18
14
10
6
1 2 3 4 5 6
Week
D
E
F
kcal Consumed as % Body Wt
80
60
40
20
**
*

**S22 Fig. Average kcal consumed per day and kcal consumed as a percentage of body weight by control no shock mice across Experiments.**

Females represented by triangles, males by diamonds; 0.4% NaCl represented by blue symbols and solid lines, 4.0% NaCl represented by red symbols and dashed lines. A, B, C) Consumption of kcal per day and D, E, F) kcal consumption as a percentage of body weight were calculated for each full week, and for the partial week at the conclusion of A, D) Experiment 1 and B, E) Experiment 2 (grey shading). Some data loss occurred on the very last weighing day for a subset of animals in C, F) Experiment 3, thus graphs and repeated measures statistical analyses for Experiment 3 calculations cease at week 6 to maximize inclusion of mice in repeated measures analyses. Experiment 1: 0.4% NaCl females, n=9; 4.0% NaCl females, n=9; 0.4% NaCl males, n=8; 4.0% NaCl males, n=9. Experiment 2: 0.4% NaCl females, n=8; 4.0% NaCl females, n=7; 0.4% NaCl males, n=9; 4.0% NaCl males, n=9. Experiment 3: 0.4% NaCl females, n=8; 4.0% NaCl females, n=8; 0.4% NaCl males, n=8; 4.0% NaCl males, n=8. Data are graphed as mean ± 95% confidence interval. $^{*}p<0.05$, $^{**}p<0.01$, $^{***}p<0.001$ indicate difference between females consuming 0.4% NaCl versus 4.0% NaCl. $^{+}p<0.05$, $^{++}p<0.01$, $^{+++}p<0.001$ indicate difference between males consuming 0.4% NaCl versus 4.0% NaCl.

S23 Figure

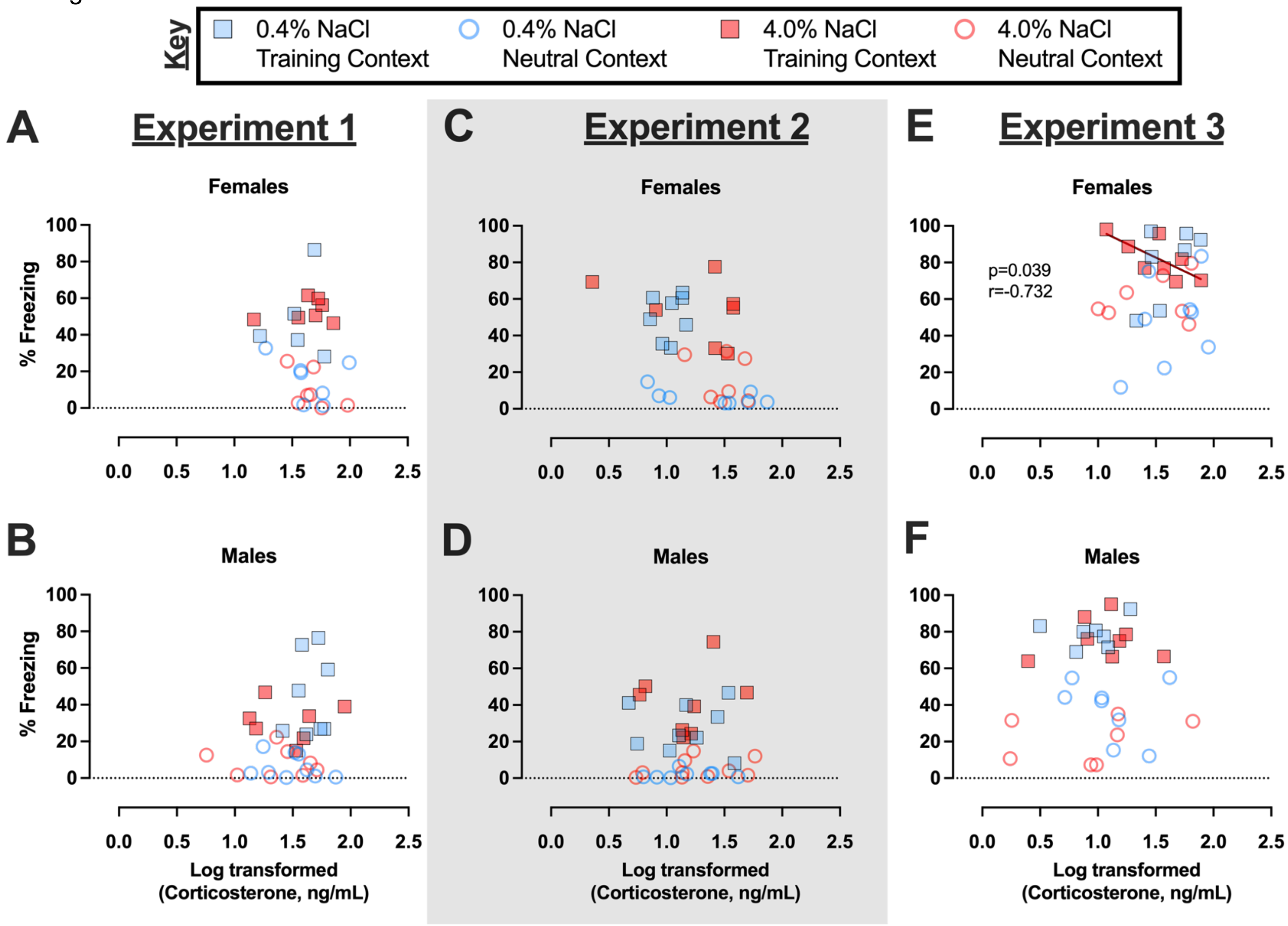

Key
0.4% NaCl Training Context
0.4% NaCl Neutral Context
4.0% NaCl Training Context
4.0% NaCl Neutral Context
A
Experiment 1
Females
B
Males
C
Experiment 2
Females
D
Males
E
Experiment 3
Females
p=0.039
r=-0.732
F
Males
% Freezing
0
20
40
60
80
100
0.0
0.5
1.0
1.5
2.0
2.5
Log transformed
(Corticosterone, ng/mL)

**S23 Fig. Correlations between context fear expression and log-transformed serum corticosterone levels across contexts and Experiments.**

Mice assigned to 0.4% NaCl represented by blue symbols, mice assigned to 4.0% NaCl represented by red symbols; mice tested in Training Context represented by squares, mice tested in Neutral Context represented by circles. Individual log-transformed serum corticosterone levels were plotted on the x-axis against the same individual's contextual fear expression during minutes two through six of the 10 min testing session plotted on the y-axis. Data for each sex were graphed for A, B) Experiment 1, C, D) Experiment 2 (grey shading), and E, F) Experiment 3. Significant correlation indicated with solid line (female). Experiment 1: 0.4% NaCl females Training Context, n=5; 0.4% NaCl females Neutral Context, n=7; 4.0% NaCl females Training Context, n=7; 4.0% NaCl females Neutral Context, n=7; 0.4% NaCl males Training Context, n=8; 0.4% NaCl males Neutral Context, n=9; 4.0% NaCl males Training Context, n=7; 4.0% NaCl males Neutral Context, n=8. Experiment 2: 0.4% NaCl females Training Context, n=8; 0.4% NaCl females Neutral Context, n=8; 4.0% NaCl females Training Context, n=7; 4.0% NaCl females Neutral Context, n=7; 0.4% NaCl males Training Context, n=9; 0.4% NaCl males Neutral Context, n=8; 4.0% NaCl males Training Context, n=8; 4.0% NaCl males Neutral Context, n=10. Experiment 3: 0.4% NaCl females Training Context, n=7; 0.4% NaCl females Neutral Context, n=8; 4.0% NaCl females Training Context, n=8; 4.0% NaCl females Neutral Context, n=7; 0.4% NaCl males Training Context, n=7; 0.4% NaCl males Neutral Context, n=8; 4.0% NaCl males Training Context, n=8; 4.0% NaCl males Neutral Context, n=7.

S24 Figure

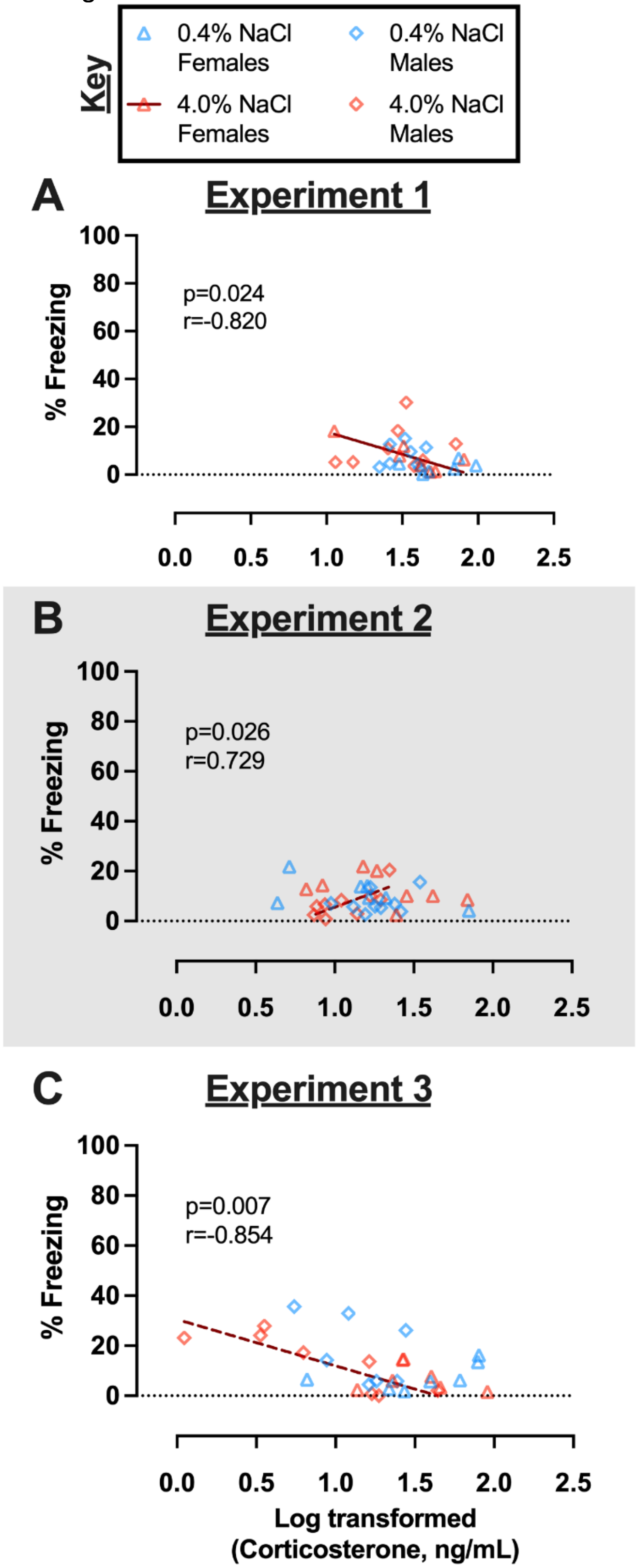
Key
0.4% NaCl Females
0.4% NaCl Males
4.0% NaCl Females
4.0% NaCl Males
A
Experiment 1
p=0.024
r=-0.820
% Freezing
100
80
60
40
20
0
0.0
0.5
1.0
1.5
2.0
2.5
B
Experiment 2
p=0.026
r=0.729
% Freezing
100
80
60
40
20
0
0.0
0.5
1.0
1.5
2.0
2.5
C
Experiment 3
p=0.007
r=-0.854
% Freezing
100
80
60
40
20
0
0.0
0.5
1.0
1.5
2.0
2.5
Log transformed
(Corticosterone, ng/mL)

**S24 Figure. Correlations between context fear expression and log-transformed serum corticosterone levels in control no shock mice across Experiments.**

Females represented by triangles, males by diamonds; 0.4% NaCl represented by blue symbols, 4.0% NaCl represented by red symbols. Individual log-transformed serum corticosterone levels were plotted on the x-axis against the same individual's contextual fear expression during minutes two through six of the 10 min testing session plotted on the y-axis. Data for each sex of control no shock mice were graphed for A) Experiment 1, B) Experiment 2 (grey shading), and C) Experiment 3. Significant correlations indicated with solid (female) or dashed (male) lines. Experiment 1: 0.4% NaCl females, n=6; 4.0% NaCl females, n=7; 0.4% NaCl males, n=8; 4.0% NaCl males, n=8. Experiment 2: 0.4% NaCl females, n=7; 4.0% NaCl females, n=8; 0.4% NaCl males, n=9; 4.0% NaCl males, n=9. Experiment 3: 0.4% NaCl females, n=7; 4.0% NaCl females, n=7; 0.4% NaCl males, n=7; 4.0% NaCl males, n=8.

**S33 Table. Details of confirmed outliers (> 5 standard deviations (SD) ± average of data set sans suspected outlier) within respective data sets. Outliers are marked with red font.**

S33A Table

| Male • Experiment 3 • 0.4%NaCl<br>No Shock • Training Context • Training baseline | | | |
|---|---|---|---|
| **Animal ID** | **Data** | **Outlier Determination** | |
| 0445 | 6.36 | | |
| 0447 | 8.93 | | |
| 0514 | 24.01 | >6 SD beyond mean | |
| 0519 | 0.88 | 2.66 | *Avg* |
| 0523 | 0.69 | 3.54 | *SD* |
| 0529 | 0.00 | 23.88 | *Avg + 6 SD* |
| 0697 | 0.00 | | |
| 0796 | 1.75 | | |

S33B Table

| Male • Experiment 1 • 4.0%NaCl<br>No Shock • Training Context • Testing | | | |
|---|---|---|---|
| **Animal ID** | **Data** | **Outlier Determination** | |
| 0331 | 30.21 | | |
| 0345 | 18.41 | | |
| 0352 | 75.65 | >7 SD beyond mean | |
| 0377 | 3.63 | | |
| 0388 | 10.92 | 11.56 | *Avg* |
| 0391 | 6.03 | 9.04 | *SD* |
| 0409 | 5.15 | 74.81 | *Avg + 7 SD* |
| 0415 | 12.92 | | |
| 0422 | 5.24 | | |

S33C Table

| Male • Experiment 1 • 4.0%NaCl<br>Shock • Training Context • Testing | | |
|---|---|---|
| **Animal ID** | **Data** | **Outlier Determination** |
| 0342 | 86.01 | >5 SD beyond mean |

| | | | |
|---|---|---|---|
| 0344 | 39.07 | 30.84 | *Avg* |
| 0357 | 33.78 | 10.65 | *SD* |
| 0367 | 21.75 | 84.09 | *Avg + 5 SD* |
| 0387 | 14.97 | | |
| 0413 | 32.57 | | |
| 0416 | 46.73 | | |
| 0423 | 27.04 | | |

S33D Table

| Female・Experiment 2・0.4%NaCl<br>Shock・Neutral Context・Testing | | | |
|---|---|---|---|
| **Animal ID** | **Data** | **Outlier Determination** | |
| 0608 | 3.56 | | |
| 0610 | 9.26 | | |
| 0613 | 3.16 | | |
| 0622 | 6.15 | | |
| 0626 | 3.12 | 6.35 | *Avg* |
| 0667 | 3.71 | 4.05 | *SD* |
| 0677 | 7.10 | 30.63 | *Avg + 6 SD* |
| 0678 | 14.73 | | |
| 0682 | 32.69 | >6 SD beyond mean | |

S33E Table

| Male・Experiment 2・0.4%NaCl<br>Shock・Neutral Context・Testing | | | |
|---|---|---|---|
| **Animal ID** | **Data** | **Outlier Determination** | |
| 0445 | 2.31 | | |
| 0447 | 2.58 | | |
| 0514 | 0.58 | 2.04 | *Avg* |
| 0519 | 6.52 | 2.05 | *SD* |
| 0523 | 0.76 | 26.61 | *Avg + 12 SD* |
| 0529 | 2.57 | | |
| 0697 | 27.97 | >12 SD beyond mean | |
| 0796 | 0.27 | | |
| 0807 | 0.75 | | |

S33F Table

| Male・Experiment 3・0.4%NaCl<br>Shock・Training Context・Testing | | | |
|---|---|---|---|
| **Animal ID** | **Data** | **Outlier Determination** | |
| 0073 | 69.01 | 79.18 | *Avg* |
| 0187 | 80.66 | 7.74 | *SD* |
| 0278 | 83.16 | 40.46 | *Avg - 5 SD* |
| 0312 | 80.07 | | |
| 0324 | 77.38 | | |
| 0428 | 92.45 | | |
| 0434 | 38.13 | >5 SD beyond mean | |
| 0436 | 71.55 | | |

S33G Table

| Male・Experiment 3・4.0%NaCl<br>Osmolality | | | |
|---|---|---|---|
| **Animal ID** | **Data** | **Outlier Determination** | |
| 0077 | 338 | >7 SD beyond mean | |
| 0188 | 315 | | |
| 0306 | 316 | | |
| 0311 | 322 | 316.29 | *Avg* |
| 0322 | 314 | 2.87 | *SD* |
| 0427 | 318 | 336.38 | *Avg + 7 SD* |
| 0435 | 314 | | |
| 0438 | 315 | | |

S33H Table

| Female・Experiment 1・0.4%NaCl<br>No Shock・Training Context・Corticosterone Log | | | |
|---|---|---|---|
| **Animal ID** | **Data** | **Outlier Determination** | |
| 0394 | 1.67 | | |
| 0531 | 1.84 | | |
| 0539 | 1.99 | | |
| 0544 | 1.64 | 1.75 | *Avg* |
| 0550 | 1.48 | 0.19 | *SD* |
| 0556 | 1.87 | 0.82 | *Avg - 5 SD* |
| 0642 | 0.79 | >5 SD beyond mean | |

S33I Table

| Female・Experiment 2・0.4%NaCl Shock・Training Context・Corticosterone Log | | | |
|---|---|---|---|
| **Animal ID** | **Data** | **Outlier Determination** | |
| 0616 | 1.13 | | |
| 0618 | 1.94 | >7 SD beyond mean | |
| 0620 | 1.17 | | |
| 0625 | 0.86 | 1.03 | *Avg* |
| 0628 | 0.88 | 0.12 | *SD* |
| 0676 | 1.05 | 1.86 | *Avg + 7 SD* |
| 0681 | 1.14 | | |
| 0683 | 1.04 | | |
| 0771 | 0.96 | | |

S33J Table

| Male・Experiment 1・0.4%NaCl・Shock Training Context・Water:NaCl Ratio Wk 1 | | | |
|---|---|---|---|
| **Animal ID** | **Data** | **Outlier Determination** | |
| 0341 | 333.63 | | |
| 0353 | 153.57 | | |
| 0354 | 441.7 | >6 SD beyond mean | |
| 0368 | 229.87 | | |
| 0389 | 218.90 | 231.82 | *Avg* |
| 0407 | 218.37 | 34.28 | *SD* |
| 0417 | 257.84 | 437.51 | *Avg + 6 SD* |
| 0419 | 210.53 | | |

S33K Table

| Male・Experiment 1・0.4%NaCl・Shock Neutral Context・Water:NaCl Ratio Wk 2 | | | |
|---|---|---|---|
| **Animal ID** | **Data** | **Outlier Determination** | |
| 0333 | 349.81 | 346.46 | *Avg* |
| 0343 | 342.23 | 84.92 | *SD* |
| 0350 | 301.87 | 1110.78 | *Avg + 9 SD* |
| 0358 | 262.82 | | |
| 0376 | 383.72 | | |
| 0390 | 486.90 | | |
| 0408 | 1127.25 | >5 SD beyond mean | |
| 0418 | 224.22 | | |

| | |
|---|---|
| 0421 | 420.07 |

S33L Table

| Male • Experiment 1 • 0.4%NaCl • Shock Neutral Context • Water:NaCl Ratio Wk 3 | | | |
|---|---|---|---|
| **Animal ID** | **Data** | **Outlier Determination** | |
| 0333 | 302.97 | 275.09 | *Avg* |
| 0343 | 258.75 | 47.54 | *SD* |
| 0350 | 272.09 | 1225.95 | *Avg + 20 SD* |
| 0358 | 188.51 | | |
| 0376 | 320.86 | | |
| 0390 | 258.38 | | |
| 0408 | 1228.7 | >20 SD beyond mean | |
| 0418 | 256.58 | | |
| 0421 | 342.59 | | |

S33M Table

| Male • Experiment 1 • 4.0%NaCl • Shock Neutral Context • Water:NaCl Ratio Wk 2.5 | | | |
|---|---|---|---|
| **Animal ID** | **Data** | **Outlier Determination** | |
| 0329 | 75.44 | 64.79 | *Avg* |
| 0332 | 59.85 | 9.38 | *SD* |
| 0340 | 65.41 | 139.82 | *Avg + 8 SD* |
| 0351 | 140.65 | >8 SD beyond mean | |
| 0371 | 75.72 | | |
| 0411 | 68.95 | | |
| 0412 | 57.52 | | |
| 0420 | 50.65 | | |

S33N Table

| Female • Experiment 2 • 0.4%NaCl • Shock Training Context • Water:NaCl Ratio Wk 4 | | |
|---|---|---|
| **Animal ID** | **Data** | **Outlier Determination** |
| 0616 | 375.63 | |
| 0618 | 900.00 | >5 SD beyond mean |
| 0620 | 433.96 | |
| 0625 | 309.68 | |

| | | | |
|---|---|---|---|
| 0628 | 376.56 | 424.32 | *Avg* |
| 0676 | 474.04 | 86.14 | *SD* |
| 0681 | 377.40 | 855.01 | *Avg + 5 SD* |
| 0683 | 454.14 | | |
| 0771 | 593.11 | | |

S33O Table

| Male・Experiment 2・0.4%NaCl・Shock Training Context・Water:NaCl Ratio Wk 2 | | | |
|---|---|---|---|
| **Animal ID** | **Data** | **Outlier Determination** | |
| 0451 | 310.26 | 329.64 | *Avg* |
| 0453 | 364.10 | 81.02 | *SD* |
| 0802 | 243.72 | 815.77 | *Avg + 6 SD* |
| 0512 | 257.58 | | |
| 0516 | 234.38 | | |
| 0520 | 459.49 | | |
| 0526 | 381.22 | | |
| 0702 | 386.36 | | |
| 0795 | 848.89 | >6 SD beyond mean | |

S33P Table

| Female・Experiment 3・0.4%NaCl・Shock Neutral Context・Water:NaCl Ratio Wk 2 | | | |
|---|---|---|---|
| **Animal ID** | **Data** | **Outlier Determination** | |
| 0559 | 777.14 | >5 SD beyond mean | |
| 0568 | 443.72 | 329.34 | *Avg* |
| 0570 | 416.26 | 74.83 | *SD* |
| 0589 | 245.56 | 703.51 | *Avg + 5 SD* |
| 0591 | 257.72 | | |
| 0651 | 270.49 | | |
| 0656 | 341.22 | | |
| 0663 | 367.35 | | |
| 0664 | 292.40 | | |

S33Q Table

| Female・Experiment 3・4.0%NaCl・Shock Neutral Context・Water:NaCl Ratio Wk 4 |
|---|

| Animal ID | Data | Outlier Determination | |
|---|---|---|---|
| 0561 | 75.48 | 75.50 | *Avg* |
| 0563 | 80.97 | 7.02 | *SD* |
| 0567 | 70.87 | 110.58 | *Avg + 5 SD* |
| 0573 | 111.94 | >5 SD beyond mean | |
| 0586 | 76.37 | | |
| 0660 | 87.56 | | |
| 0662 | 67.32 | | |
| 0665 | 69.94 | | |
